# Top predators govern multitrophic diversity effects in tritrophic food webs

**DOI:** 10.1101/2020.07.31.230375

**Authors:** Ruben Ceulemans, Christian Guill, Ursula Gaedke

**Affiliations:** University of Potsdam, Germany

## Abstract

It is well known that functional diversity strongly affects ecosystem functioning. However, even in rather simple model communities consisting of only two or, at best, three trophic levels, the relationship between multitrophic functional diversity and ecosystem functioning appears difficult to generalize, due to its high contextuality. In this study, we considered several differently structured tritrophic food webs, in which the amount of functional diversity was varied independently on each trophic level. To achieve generalizable results, largely independent of parametrization, we examined the outcomes of 128, 000 parameter combinations sampled from ecologically plausible intervals, with each tested for 200 randomly sampled initial conditions. Analysis of our data was done by training a Random Forest model. This method enables the identification of complex patterns in the data through partial dependence graphs, and the comparison of the relative influence of model parameters, including the degree of diversity, on food web properties. We found that bottom-up and top-down effects cascade simultaneously throughout the food web, intimately linking the effects of functional diversity of any trophic level to the amount of diversity of other trophic levels, which may explain the difficulty in unifying results from previous studies. Strikingly, only with high diversity throughout the whole food web, different interactions synergize to ensure efficient exploitation of the available nutrients and efficient biomass transfer, ultimately leading to a high biomass and production on the top level. The temporal variation of biomass showed a more complex pattern with increasing multitrophic diversity: while the system initially became less variable, eventually the temporal variation rose again due to the increasingly complex dynamical patterns. Importantly, top predator diversity and food web parameters affecting the top trophic level were of highest importance to determine the biomass and temporal variability of any trophic level. Overall, our study reveals that the mechanisms by which diversity influences ecosystem functioning are affected by every part of the food web, hampering the extrapolation of insights from simple monotrophic or bitrophic systems to complex natural food webs.

## Introduction

In the face of rapid global biodiversity loss (Pimm et al., 2014), investigating the influence of biodiversity on ecosystem functioning is a highly important area of research. It has become clear that biodiversity is a predominant factor in determining relevant functions of ecosystems such as biomass production, resource use efficiency, and stability (Hooper et al., 2005; Tilman et al., 2006; Worm et al., 2006). A major factor affecting the link between biodiversity and these ecosystem functions is functional diversity, i.e., the range of differences between the functions of species contained within the ecosystem (Petchey and Gaston, 2006).

Mechanistically motivated studies into the role of functional diversity have mainly been performed in the context of simple communities consisting only of one or, at best, two trophic levels. Many of these studies restricted their focus to primary producer diversity, and were able to show its correlation with relevant ecosystem functions (reviewed by Cardinale et al. (2011)). However, during the last two decades, more sophisticated theoretical and experimental studies linking both plant and consumer diversity to these ecosystem functions were conducted (see Thébault and Loreau (2003); Tirok and Gaedke (2010); Borer et al. (2012); Filip et al. (2014); Klauschies et al. (2016); Schneider et al. (2016); Seabloom et al. (2017); Flöder et al. (2018), and reviews by Duffy et al. (2007); Griffin et al. (2013); Barnes et al. (2018)). In a recent experimental study, Wohlgemuth et al. (2017) demonstrated that producer diversity effects on the biomass distribution and production at higher trophic levels crucially depends on particular traits of the consumer level, such as specialization and selectivity. Such studies highlight how the links between multitrophic functional diversity and ecosystem functioning are difficult to generalize, due to their high contextuality. The specific food webs that are studied, and the theoretical models used to study them, are often too different to enable a meaningful attempt at synthesis of their findings (Thébault and Loreau, 2003; Barnes et al., 2018).

For this reason there is a clear need to understand the effects of diversity on ecosystem functions in a setting that is as general and context-free as possible. In addition, the high degree of interplay already observed between diversity of the primary producer and herbivore consumer level underlines the importance of including diversity of even higher trophic levels. In this study, we want to advance our understanding of how functional diversity affects ecosystem functioning in model communities by including a diverse third trophic level. While it has often been highlighted how important the effects of the third trophic level on ecosystem functions are (Bruno and O’Connor, 2005; Duffy et al., 2007; Daam et al., 2019; Abdala-Roberts et al., 2019; Ehrlich and Gaedke, 2020), relatively few studies have attempted to explicitly take these effects into account. Ceulemans et al. (2019) showed that functional diversity increases the biomass production, temporal stability, and biomass transfer efficiency to higher trophic levels of a tritrophic food web, when diversity is increased simultaneously at all three trophic levels. This model analyzed one particular food web structure in detail, which raises the question whether the observed trends are to be expected in general, or whether they are context-dependent as well.

Our study tackles this issue by investigating several different tritrophic food web configurations with respect to the same ecosystem functions. Such a method has been applied successfully in the past (Gilman et al., 2010; Kovach-Orr and Fussmann, 2013; Poisot et al., 2013), but this study is the first where the diversity can be independently controlled on three trophic levels. We investigated eight different food web configurations (Figure 1), which differ in the trophic location at which functional diversity may be present. We measured functional diversity of a trophic level by the difference between the functional traits of the two species residing there: when the trait difference between the species is large, so is the functional diversity, and vice-versa. In this way, we were able to change the functional diversity of a trophic level without changing the number of species. Adopting such a trait-based rather than species-specific approach by analyzing functional diversity through trait differences, instead of using non-functional metrics of biodiversity such as species number, allows us to produce results of high generality (McGill et al., 2006; Hillebrand and Matthiessen, 2009; Krause et al., 2014). Furthermore, the relatively simple and general structure of our food webs (see Figure 1) makes our results accessible for verification by experimental studies, as they are often limited in how much complexity can be included.

**Figure 1:**
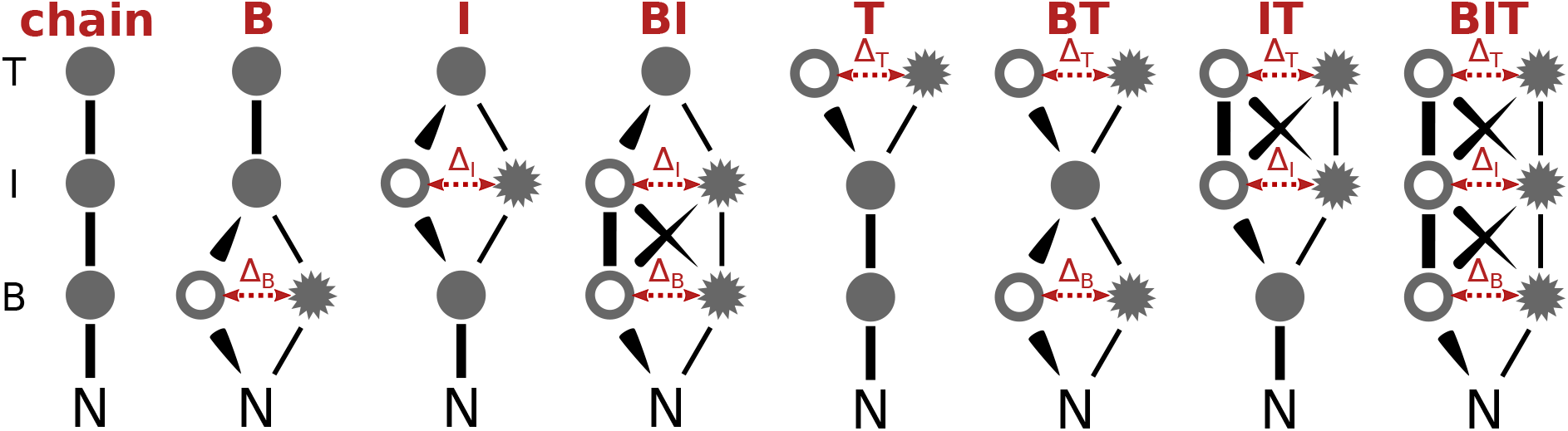
Schematic overview of the 8 different food webs compared in this study, which differ by the trophic levels (*B* for basal, *I* for intermediate, and *T* for top) on which diversity is possible (indicated above). In this way, chain refers to the linear chain which contains no diversity, B to the food web on which only the basal level is diverse, etc., and finally BIT denotes the food web which contains diversity on all trophic levels. The thickness of the connections between the nodes illustrates the comparative intensity of the trophic interaction, which is determined by the amount of diversity, or the trait difference, between the species on each trophic level (Δ_*B*_, Δ_*I*_, and Δ_*T*_). Each of these food webs are analyzed as general as possible, with independently varying amounts of trait differences and parameters drawn randomly from biologically plausible intervals.

Our model rests on few very general assumptions. The first is allometry, which states that larger organisms tend to grow slower than smaller ones (Kalinkat et al., 2013). Combined with the assumption that consumers tend to be larger than their prey we obtain the general property that the mean growth rate should decrease as the trophic level increases. This strictly holds for pelagic systems (Gaedke and Kamjunke, 2006), but also for other ones, except for the plant-herbivore interface (Brose et al., 2006). The third basic assumption is the frequently established trade-off between growth rate and defense (Herms and Mattson, 1992; Hillebrand et al., 2000; Kneitel and Chase, 2004; Ehrlich et al., 2017, 2020). It implies that slow growing species are generally less affected by grazing than faster growing species, which invest less in defense mechanisms due to energetic limitations. In addition, the non-grazing mortality terms (see Eq. (8)) are of general nature and may be due to several different processes, such as basal respiration, the influence of parasites and viruses, or outflow in an experimental microcosm.

Importantly, food web dynamics do not only depend on the topology of the food web, but also on the specific parametrization used, regarding both external environmental parameters as well as internal parameters such as growth rates, attacks rates, and handling times. To sufficiently capture the potentially high variation in biomass dynamics, we randomly selected a total of 128, 000 parameter combinations from ecologically plausible intervals for the eight different food webs, as well as tested 200 initial conditions per parameter combination. These parameter values were drawn from intervals geometrically centered around values which are particularly relevant for planktonic systems (Ceulemans et al., 2019), but are sufficiently wide to capture the behavior of many different types of food webs (see Table 1). This procedure allows us to obtain results of high generality, as they apply to the average behavior of tritrophic systems, independent of its parametrization.

**Table 1:** Name and meaning of the parameters that were used in the study, along with the range from which they were sampled. For example, the nutrient inflow concentration *N*_0_ was randomly sampled from the interval [1/2, 2]·1120 ≈ [560, 2240] *μg* N *l*. In this table, B-I refers to the functional response between the Basal (B) and the Intermediate (I) trophic level, and I-T to the Intermediate and Top (T) level. The bottom three parameters were kept at fixed values.

| Parameter | Meaning | Range |
| --- | --- | --- |
| $N_0$ | nutrient inflow concentration | $[1/2, 2] \cdot 1120 \mu g N/l$ |
| $h_N$ | nutrient uptake half-saturation const. | $[1/2, 2] \cdot 10 \mu g N/l$ |
| $r'_0$ | basal growth rate | $[1/2, 2] \cdot 1/\text{day}$ |
| $a_0$ | B-I attack rate | $[1/2, 2] \cdot 1.04 \cdot 10^{-3}/(\text{day} \cdot \mu g C/l)$ |
| $h_0$ | B-I handling time | $[1/2, 2] \cdot 1.15 \cdot \text{day}$ |
| $\alpha_0$ | I-T attack rate | $[1/2, 2] \cdot 4.48 \cdot 10^{-4}/(\text{day} \cdot \mu g C/l)$ |
| $\eta_0$ | I-T handling time | $[1/2, 2] \cdot 2.62 \cdot \text{day}$ |
| $\delta$ | inflow rate | $[0.03, 0.06] \cdot 1/\text{day}$ |
| $a_{\text{scale}}$ | cross link scaling factor | $[1, 500]$ |
| $n$ | B-I Hill exponent | $[1, 2]$ |
| $\nu$ | I-T Hill exponent | $[1, 2]$ |
| Parameter | Meaning | Value |
| $e$ | biomass conversion efficiency | 0.33 (not varied) |
| $c_N/c_C$ | basal nitrogen-to-carbon ratio | 0.175 (not varied) |
| $\tau_{\text{inc}}$ | maximal trait increase | 1/2 (not varied) |

## Methods

The numerical data used in our study was obtained by storing the mean biomasses and coefficients of variation (*CV*s) of the following ordinary differential equation model:

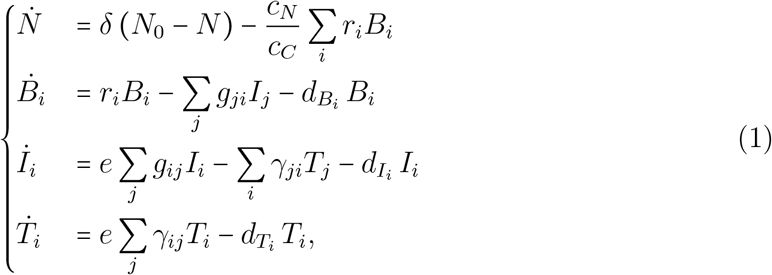

where the indices *i, j* ∊ {1, 2}. *N* describes the free inorganic nutrients in the system, with the inflow concentration *N*_0_, inflow rate *δ*, and nutrient-to-carbon ratio 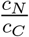. The loss rates 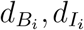 and 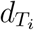 represent losses proportional to the biomass present, such as basal respiration, sedimentation, or wash-out. The basal species’ *B_i_* uptake rate *r_i_* is described by their maximal growth rate 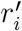 and nutrient uptake half-saturation constant *h_N_*:

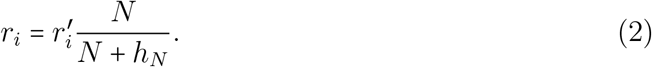

The interaction between the intermediate species *I_i_* and the basal species *B_j_* is described by a Holling-Type-III functional response which is determined by the attack rate *a_ij_*, handling time *h_ij_* and the Hill exponent *n*:

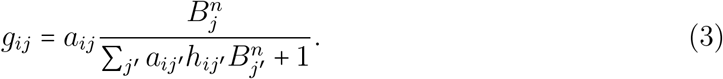

In the same way, the interaction between top species *T_i_* and intermediate species *I_j_* is given by:

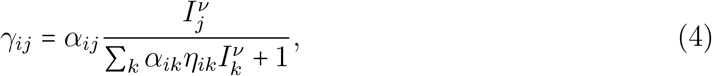

with attack rate *α_ij_*, handling time *η_ij_*, and Hill exponent *ν*. Finally, the biomass conversion efficiency for the intermediate and top species is described by *e*.

### Influence of trait differences on trait parameters

The parameters 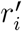, *h_ij_, a_ij_, η_ij_, α_ij_*, and all death rates are determined by the trait differences Δ_*B*_, Δ_*I*_, and Δ_*T*_, which each can vary from 0 (the two species at each trophic level are equal) to 1 (maximal trait differences). As trait differences increase, the species *B*_1_, *I*_1_, and *T*_1_ will be metabolically more active, whereas *B*_2_, *I*_2_, and *T*_2_ will be less active through modifying their maximal feeding rates (which equal the inverse of the handling times *h_ij_* and *η_ij_* for the intermediate and top species).

In our model, trait differences affect the relevant species’ parameters symmetrically, such that an increase for species 1 leads to a decrease for species 2 by the same factor. Explicitly:

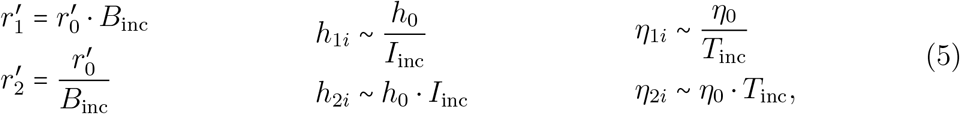

with

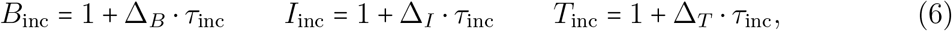

so that they are unity for Δ_*i*_ = 0, leaving the species’ parameters unaffected, and maximal for Δ*i* = 1, where *τ*_inc_ determines their maximal increase. Note that the handling times *h_ij_* and *η_ij_* depend on the trait differences of both the predator and the prey level, hence the proportional relationship (~) instead of equality (more information is provided below, cf. Equations (7) & (9)).

The universality of trade-offs in natural systems (Kneitel and Chase, 2004; Ehrlich et al., 2017) implies that for any increase or decrease in growth rates, the species’ loss rates must change correspondingly. Time and/or energy that is invested towards a certain defense strategy cannot be used for resource uptake, and thus, comes at the cost of a lower growth rate (and thus a higher handling time). Conversely, investing in a higher growth rate (lower handling time) tends to make a species more vulnerable to predation as it leaves less time and/or energy for employing defense strategies. For simplicity, the loss rates are affected in the same way as the growth rates. Thus, *B*_inc_ affects the handling times *h_ij_* as well as the death rates 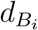, *I*_inc_ affects *η_ij_* and 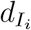, and *T*_inc_ affects 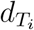, in the following way:

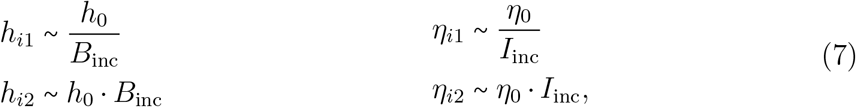

and,

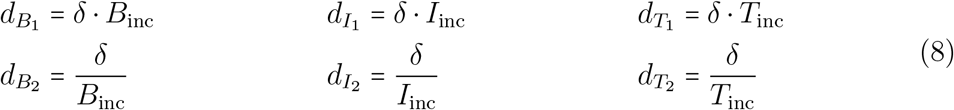

The handling times *h_ij_* and *η_ij_* are thus dependent on both *B*_inc_ & *I*_inc_, or *I*_inc_ & *T*_inc_, respectively. While the linear relationship that describes this dependence is almost certainly a simplification of biological reality, specifying a more complex relationship might make our model unnecessarily more complicated. As described in the next section, multiple parameter combinations will be investigated, which means that our approach is not limited to one single distinct trade-off curve.

Summarizing:

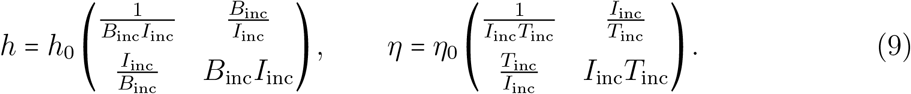

The interaction between predator-prey pairs is not only determined by the handling times *h_ij_* and *η_ij_*, but also by the attack rates *a_ij_* and *α_ij_*. In our model, these are responsible for determining the relative strength of the “cross” links between two adjacent trophic levels (e.g. *B*_1_ → *I*_2_, etc.). As the functional diversity on adjacent trophic levels increases, these “cross” links will decrease in strength relative to the “parallel” links (e.g. *B*_1_ → *I*_1_, etc.). The rate at which their strength decreases is determined by the attack rate scaling parameter *a*_scale_. For details see Appendix S1. In this way, it is possible to describe a tightly linked food web for *a*_scale_ ≈ 1, two largely separated tritrophic chains for *a*_scale_ ≫ 1, or an intermediate situation.

### Parameter selection

In order to capture a high diversity of dynamical outcomes, within a plausible ecological setting, the parameters of the food web were sampled uniformly from certain intervals determined by a standard value from which the boundaries are calculated (Table 1). These standard values are based on Ceulemans et al. (2019), and describe an ecologically realistic planktonic system with three trophic levels. In particular, the maximal growth rates (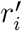, *e/h*_0_, & *e/η*_0_) were set to correspond to an allometrically scaled food chain with the body mass ratios between adjacent trophic levels of 10^3^, with an allometric scaling exponent of −0.15. However, due to the spread of the intervals the actual ratio between body masses (assuming the same scaling exponent *λ*) can vary between approximately 1 and 10, 000, 000 (for details, see Appendix S2). In this way, a good balance is made between capturing a high amount of dynamical variation, while still being ecologically realistic.

The trait difference parameters can take the following values:

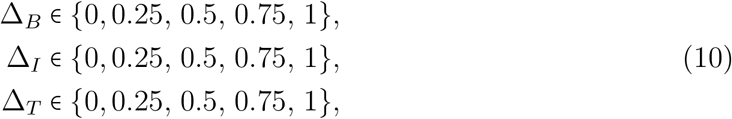

so that there are 125 combinations possible. These determine both the specific food web topology and the amount of functional diversity present. For example, Δ_*B*_ = 1, Δ_*I*_ = 0.25, and Δ_*T*_ = 0 implies that we are investigating the BI food web (Figure 1), where the basal level is highly diverse, but the species on the intermediate level are still relatively similar.

In order to sample a large part of all the possible dynamical outcomes that can be exhibited by our model, we randomly sampled 1024 different parameter combinations, for each selection of Δ_*B*_, Δ_*I*_, and Δ_*T*_ (Eq. 10). Moreover, for every parameter combination, 200 different initial conditions were tested to capture potential alternative stable states. These initial values were randomly sampled such that the total amount of biomass in the initial state did not exceed 2 · *N*_0_. The system was allowed to relax to its attractor before the mean biomasses and the *CV* of each species, and of each trophic level, were recorded for a sufficiently long time period. More detailed information on this procedure can be found in Appendix S3. Numerical integration of the ordinary differential equations in Eq. (1) was done in *C* with the SUNDIALS CVODE solver version 2.7.0 (Hindmarsh et al., 2005). Subsequent analysis of the food web data was performed in Python 3.6 using NumPy (Van Der Walt et al., 2011), pandas (McKinney, 2010), and Matplotlib (Hunter, 2007). Further details on our computational procedure, as well as the code itself and the data required to produce Figures 3–6 and various Appendix Figures can be found as an electronic supplement.

### Random Forest Model

In order to simplify the presentation of our results, and to easily extract additional relevant information, we trained a Random Forest model on our dataset. A detailed description of how this works can be found in Appendix S3. Essentially, Random Forests are a class of machine learning models which are popularly used due to their relatively simple structure and high versatility (Breiman, 2001; Cutler et al., 2007; Thomas et al., 2018).

For each quantity of interest (see Results), an Extremely Random Forest consisting of 2000 trees was trained using the Scikit-learn (Pedregosa et al., 2011) package in Python. Using only those parameter combinations that lead to coexistence of all species in the food web, the training dataset consisted of the 14 different parameters (see Table 1 and Eqs. 10) as input values, and the mean biomass and *CV* s of each trophic level as output values. During training, the random forest algorithm performed cross-validation by calculating the Out-Of-Bag (OOB) score, to estimate its accuracy. After training the random forest model, we used it to investigate how the basal, intermediate, and top diversity (Δ_*B*_, Δ_*I*_, and Δ_*T*_) affect the quantities of interest, independently of all other parameters, by examining the partial dependency graphs. Finally, the random forest also provided us with a measure of the importance of each of the input parameters in determining the desired outcome (relative importance).

## Results

In order to understand in which ways diversity of different trophic levels affects tritrophic systems, we analyzed the solutions of the ordinary differential equation model presented in the Methods (Equation (1)) for 128, 000 different parameter combinations. For each parameter combination, we saved the mean nutrient concentration and biomass density (in short biomass) and coefficient of variation (CV) of each individual population and trophic level over a long period of time (see also Appendix S3). In the main text, we will focus in particular on diversity effects on:

- the nutrient concentration *N* and biomass per trophic level *B_B_, B_I_, B_T_* (see Figure 3); and
- the *CV* of the nutrient concentration and biomass per trophic level *CV_N_, CV_B_, CV_I_, CV_T_*(see Figure 4).

Based on the mean biomasses, we also calculated several quantities related to the flow of energy through the food web. The following ones are shown in the main text (see Figure 5, and Equation (1) and Appendix S5 for more information):

- the biomass production on the top level 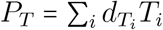;
- the amount of basal biomass flowing upward to the intermediate level *B*_up_;
- the production to biomass ratio of the basal level 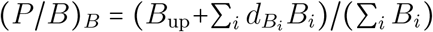; and
- the food web efficiency, defined as the ratio between the biomass production at the top and the basal level: *P_T_/P_B_*.

Figures 3–5 are partial dependence graphs revealing how trait differences on the basal (Δ_*B*_), intermediate (Δ_*I*_), and top (Δ_*T*_) level affect the quantity of interest. Such partial dependence graphs are calculated from the Random Forest model trained on the food web data, and show the average value of the quantity of interest, independent of all other model parameters (see Methods). This presentation allows us to concisely capture the full behavior of all food webs, as they each occupy a certain location in the partial dependency graphs (Figure 2). A concise summary of our main findings is presented in Table 2.

**Figure 2:**
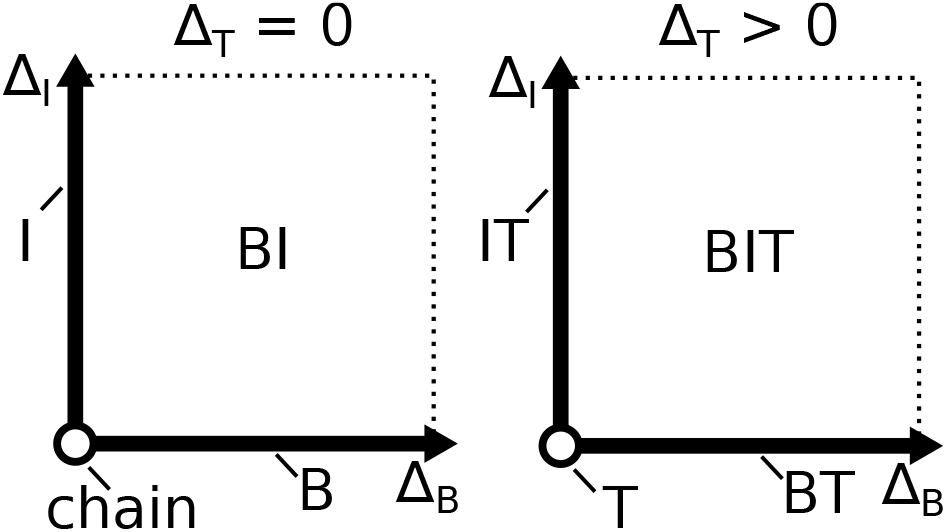
Pictorial representation of the location of the different food webs (Figure 1) in the partial dependence graphs in Figures 3–5. On the left-side graph (Δ_*T*_ = 0, i.e., no diversity at the top level), the chain is on the point (0, 0) (Δ_*B*_ = Δ_*I*_ = 0), the B food web is located on the line Δ_*I*_ = 0, the I food web is located on the line Δ_*B*_ = 0, and the BI web is located in the plane where both Δ_*B*_ and Δ_*I*_ are non-zero. Similarly, on the right-side graph where Δ_*T*_ > 0 (either *low* or *high* in Figs. 3–5), the T web is located on the point (0; 0) (Δ_*B*_ = Δ_*I*_ = 0), the BT food web is located on the line Δ_*I*_ = 0, the IT food web is located on the line Δ_*B*_ = 0, and finally the BIT web is located in the plane where Δ_*B*_, Δ_*I*_, and Δ_*T*_ are non-zero.

**Figure 3:**
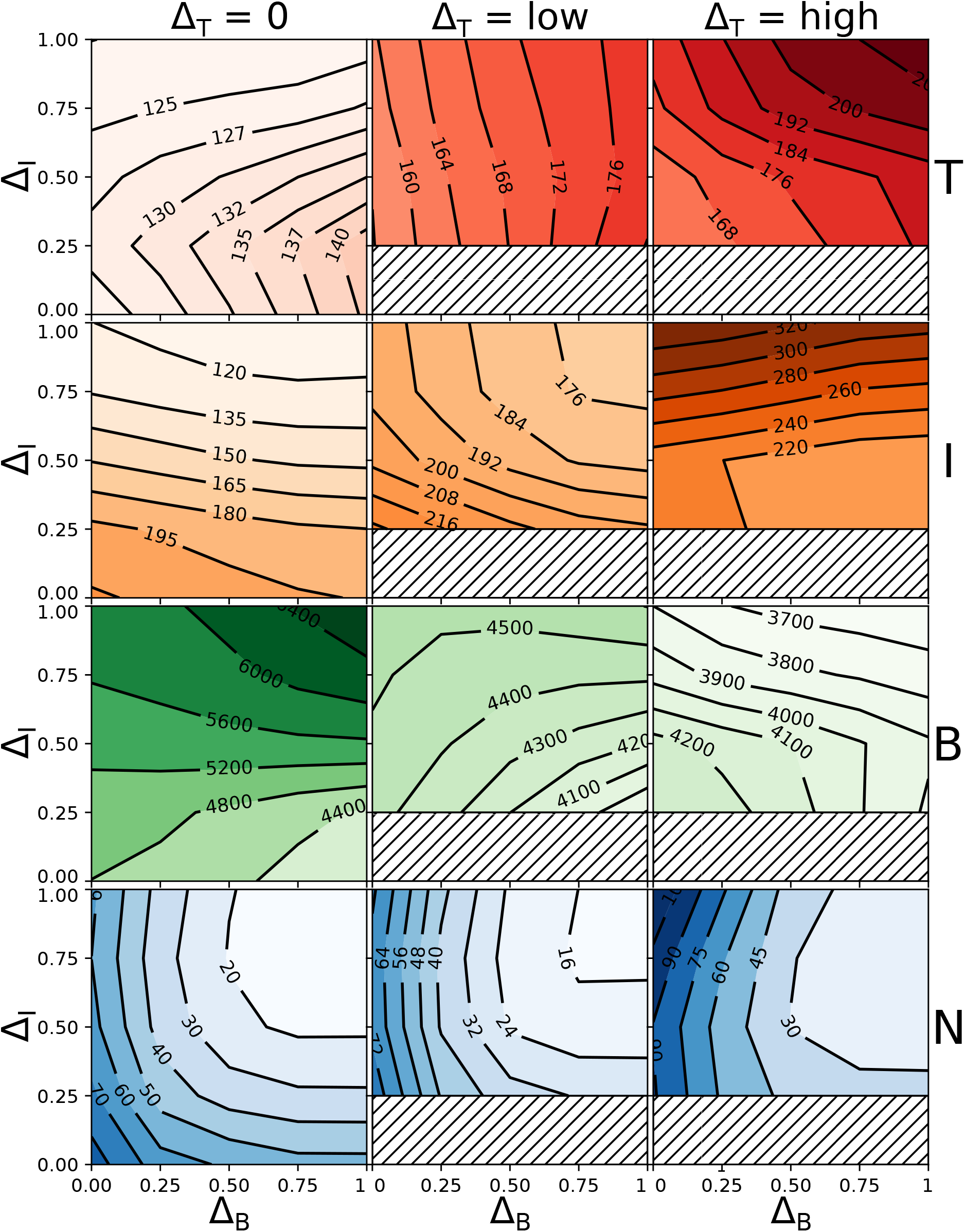
Effect of trait differences at the basal (Δ_*B*_), intermediate (Δ_*I*_), and top (Δ_*T*_) trophic level on the free nutrient concentration (*N*, blue), and the biomasses on the basal (*B*, green), intermediate (*I*, orange), and top (*T*, red) trophic level, displayed as partial dependence graphs. To simplify the presentation, the effects of Δ_*B*_ and Δ_*I*_ are shown separately for three levels of Δ_*T*_: Δ_*T*_ = 0 (left), low Δ_*T*_ (0.25 and 0.5, middle), and high Δ_*T*_ (0.75 and 1, right). Figure 2 shows a detailed explanation on how to read this figure. These graphs show the expected trends of *N, B, I,* & *T* as the amount of diversity on any trophic level is varied (for more information see Methods and Appendix S5). For example, in the chain (lower left corner of each subplot for Δ_*T*_ = 0), *T* is expected to be much lower than in the highly diverse BIT web (upper right corner for Δ_*T*_ = *high*). When Δ_*T*_ is nonzero, the region below Δ_*I*_ = 0.25 (T and BT webs) cannot be shown as no coexisting parameter combination exists here due to the two distinct top species sharing only one resource.

**Figure 4:**
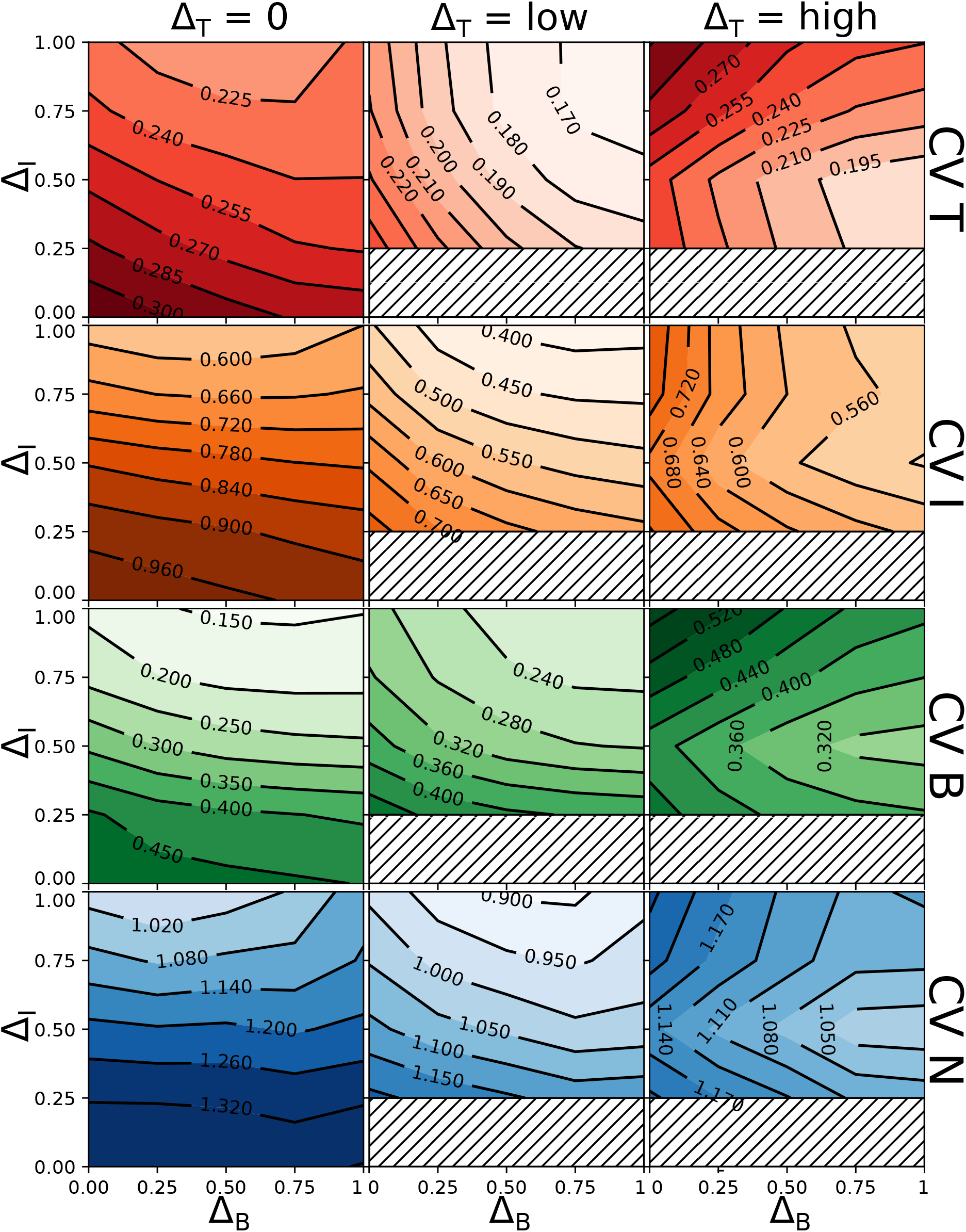
Effect of trait differences at the basal (Δ_*B*_), intermediate (Δ_*I*_), and top (Δ_*T*_) trophic level on the coefficient of variation (*CV*) of the free nutrient concentration (*N*, blue), the total biomass at the basal (*B*, green), intermediate (*I*, orange), and top (*T*, red) trophic levels, displayed as partial dependence graphs. Consult Figures 2 and 3 for a detailed description on how to read this Figure. Strikingly, we can see that Δ_*T*_ has a non-monotonous effect on the temporal variability of the whole food web: a moderate amount of top predator diversity tends to decrease the temporal variation, but adding yet more diversity to the food web causes it to increase again.

**Figure 5:**
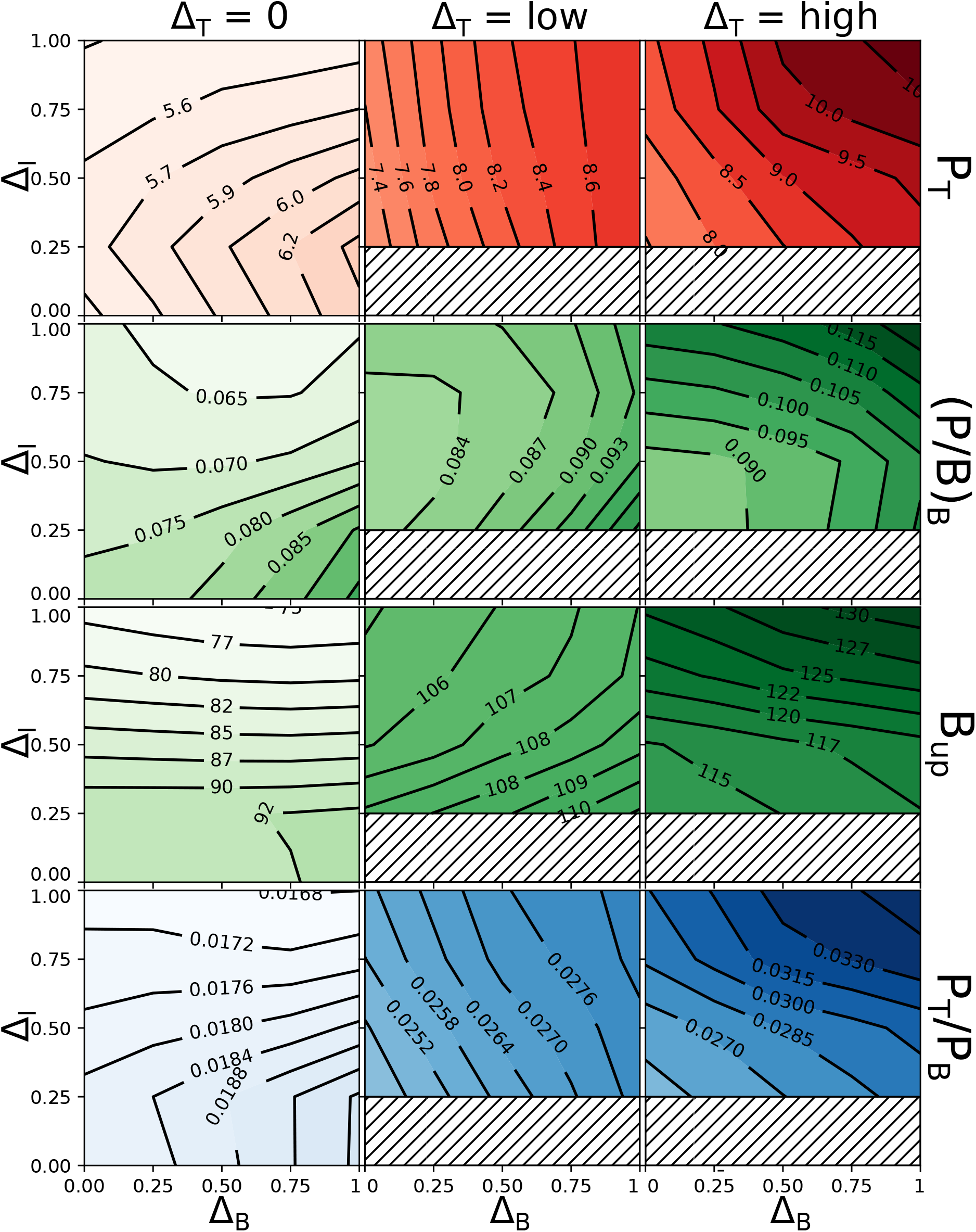
Effect of trait differences at the basal (Δ_*B*_), intermediate (Δ_*I*_), and top (Δ_*T*_) trophic level on several different metrics related to the flow of biomass and energy through the food web, displayed as partial dependence graphs. From top to bottom: top biomass production *P_T_*, basal production to biomass ratio ((*P/B*)_*B*_), basal biomass owing to *I* (*B*_up_), and the food web efficiency *P_T_*/*P_B_*. Consult the Results and Appendix S5 for more information on these quantities, and Figures 2 and 3 for a detailed description on how to read this Figure. In the chain (lower left corner for Δ_*T*_ = 0), we observe for example a much lower *P_T_*/*P_B_* than in the highly diverse BIT web (upper right corner for Δ_*T*_ = *high*), which means biomass produced by the basal trophic level is transferred much more efficiently to the top level.

**Table 2:**
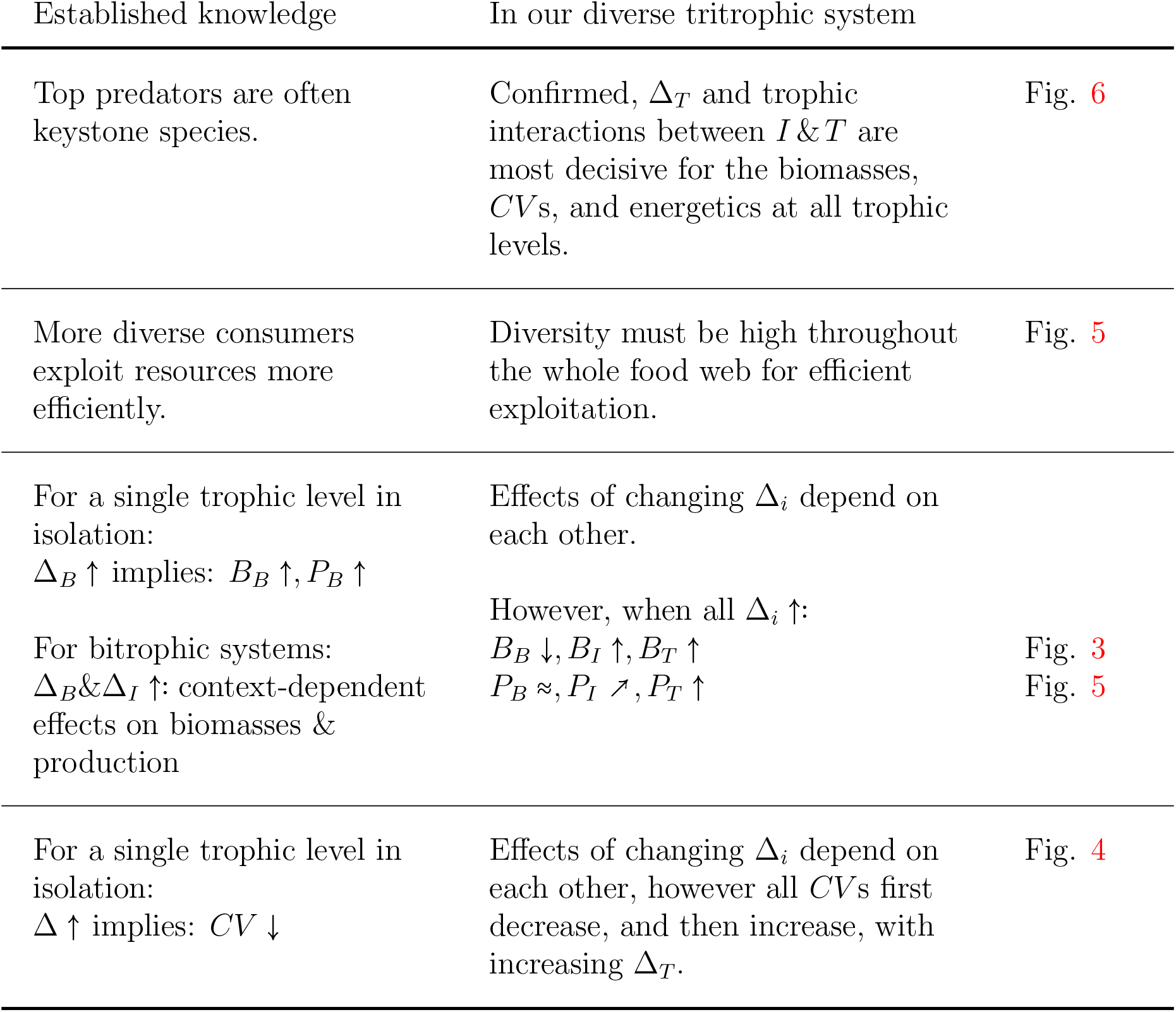
Comparison of established knowledge of the link between the functional diversity and certain ecosystem functions of communities consisting of one or two trophic levels (see text for references), to our model where diversity can be changed at three trophic levels (see Figure 1). Δ_*i*_, B_*i*_, and *P_i_* refer to the diversity, biomass, and biomass production at trophic level *i* ∈ {*B, I, T*}, respectively (see Figure 1 and Results), and the arrows indicate the direction in which these quantities are changing (↑:increase, ↗: moderate increase, ≈: approximately constant, ↓ decrease). Our model enables us to understand the mechanisms responsible for these top-down and bottom-up effects, which simultaneously cascade through the food web.

In most cases the OOB scores, which measure the accuracy of the Random Forest models, were above 0.60, with some exceptions (Table 3). Such scores indicate a sufficient model accuracy as we focus on the average trends in the predicted quantity as a function of the functional diversity of different trophic levels, rather than on predictions for specific parameter values.

**Table 3:**
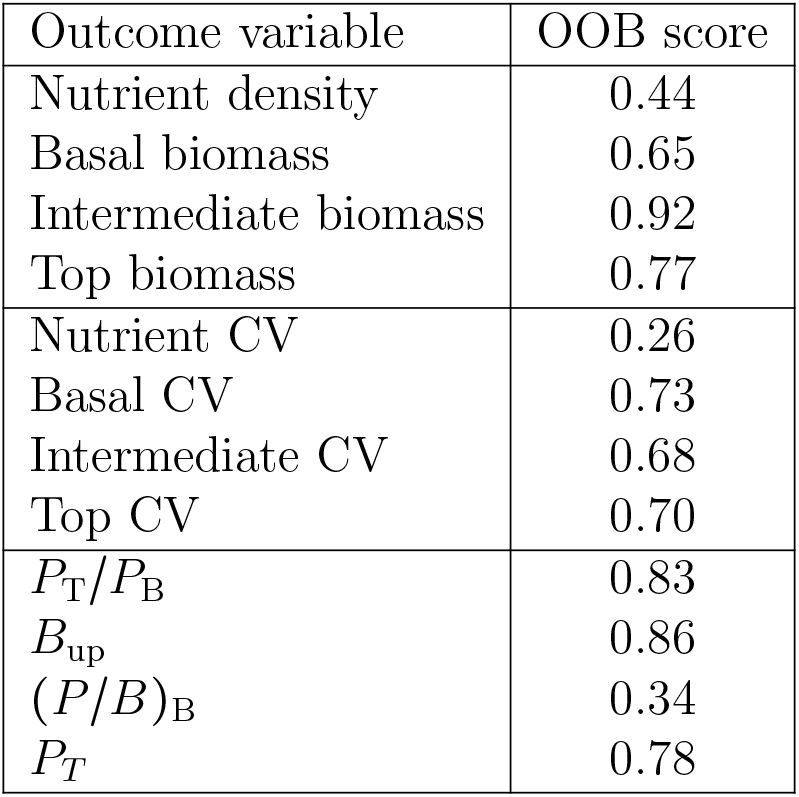
OOB scores estimating the accuracy of the random forest model, for all outcome quantities. An OOB score of 1 represents a perfect model prediction, whereas an OOB score of 0 means that the model is as accurate as simply predicting the mean outcome value every time.

The quantities of interest were only examined for those initial conditions and parameter combinations that actually led to coexistence of all species originally present (see Figure 1). Interestingly, there were only very few parameter combinations that led to coexistence for the Τ (1 combination) and BΤ (8 combinations) food webs (see Appendix S4). One of the two top species almost always outcompeted the other in these webs. As we cannot reliably investigate the behavior of these food webs in general, we did not include these parameter combinations in our dataset. This implies that our dataset contains no data points with Δ_*T*_ > 0, Δ_*I*_ = 0, and therefore the region below Δ_*I*_ = 0.25 for Δ_*T*_ > 0 in Figures 3–5 remains empty.

### Nutrient concentration and biomasses

The partial dependency graphs of the free nutrient concentration and the biomasses on each trophic level on the trait differences Δ_*B*_, Δ_*I*_, and Δ_*T*_ (Figure 3) reveal strong differences between the simple chain without any diversity (Δ_*B*_ = Δ_*I*_ = Δ_*T*_ = 0), and the food web with high trait differences at every trophic level (Δ_*B*_ = Δ_*I*_ = 1, and Δ_*T*_ = high). Comparing these two points shows that the linear chain has a higher average free nutrient concentration and a lower intermediate and top biomass than the diverse food web.

In between these two extremes, the tritrophic structure of our model gives rise to several interesting patterns. Comparing the chain and the B, I and BI food webs (i.e. Δ_*T*_ = 0, Figure 3, left panels) shows that when Δ_*I*_ is 0 or low, increasing Δ_*B*_ leads to a decrease in basal biomass, whereas if Δ_*I*_ is high, this pattern reverses as the basal biomass increases with Δ_*B*_. In other words, if functional diversity is only present on the basal level, basal biomass tends to decrease with Δ_*B*_. However, taking consumer diversity into consideration in the BI food web shows that this pattern is not general and strongly depends on the actual level of consumer diversity (Δ_*I*_).

Investigating the effect of Δ_*I*_ and Δ_*T*_ on the intermediate and top level biomasses shows exactly the same pattern. When Δ_*T*_ is 0, intermediate biomass tends to decrease as Δ_*I*_ increases, whereas when Δ_*T*_ is high, it increases with Δ_*I*_ (independently of Δ_*B*_). Additionally, it is clear that top biomass increases with Δ_*T*_ in a gradual fashion.

The location and strength of trophic cascading in the food web is also affected by the amount of functional diversity present on the different trophic levels. For example, when Δ_*T*_ is zero, an inverse relationship between the biomass on the intermediate and basal level can be observed, whereas the top level biomass seems hardly affected by Δ_*B*_ and Δ_*I*_ (Figure 3). When Δ_*T*_ is low, biomasses at the top and intermediate level are strongly negatively correlated, indicating that a diverse top level is able to exert a stronger influence on the whole food web as compared to a non-diverse top level. This negative relationship does not cascade downwards to the basal level, potentially due to the buffering properties of a diverse intermediate level. However, for Δ_*T*_ = high, the strong inverse relationship between top and intermediate biomass is replaced by a rather positive one, due to the sharp increase in intermediate biomass as Δ_*I*_ is increased.

### Temporal variation

We also examined how the functional diversity at each trophic level (Δ_*B*_, Δ_*I*_, and Δ_*T*_) influences the temporal variation of the nutrients and biomasses per trophic level, by calculating the coefficient of variation (*CV*) (Figure 4). One clear overarching pattern is the covariation of the *CV* s along the different trophic levels. Temporal fluctuations at any trophic level propagate through the whole food web, affecting all other levels.

The left column shows how Δ_*B*_ and Δ_*I*_ affect the *CV* s of the food webs without top diversity (Δ_*T*_ = 0). In this case, the *CV* of any trophic level depends almost solely on Δ_*I*_. Only the *CV* of the nutrient concentration depends strongly on Δ_*B*_.

These results are strongly affected by the top diversity. By increasing Δ_*T*_ from 0 to *low*, all *CV* s are considerably dampened. However, this trend reverses as Δ_*T*_ is increased further, as all *CV* s tend to increase again (Δ_*T*_ = high). Hence, while comparing the simple chain (Δ_*B*_ = Δ_*I*_ = Δ_*T*_ = 0) to the food web with high trait differences ((Δ_*B*_ = Δ_*I*_ = 1, and Δ_*T*_ = high) does not immediately show any notable differences, it is clear that temporal variability is strongly affected in an intricate way by the amount of functional diversity at the different trophic levels.

Additionally, there is a strong correlation between the *CV* of the basal trophic level, and the free nutrient concentration (Figure 3, bottom row). A low temporal variability on the basal level leads to a strong increase in nutrient exploitation efficiency, and therefore low nutrient concentrations.

### Biomass production and food web energetics

We also analyzed metrics related to biomass production and food web energetics: biomass production on the top level *P_T_*, basal biomass flowing to the intermediate level *B*_up_, basal biomass to production ratio (*P/B*)_B_, and the food web efficiency *P*_T_/*P*_B_ (Figure 5, and Appendix S5 for more information on these quantities). Examining these (and related, see Appendix S5) quantities helped us to understand why the biomass at the top level is highest when functional diversity everywhere is high (top right corner in Figure 3 for Δ_*T*_ = *high*). Importantly, we can infer the quantities *P_I_* (total biomass production of the intermediate level) and *I*_up_ (biomass flowing from the intermediate to the top level) from *B*_up_ and *P_T_*: *P_I_ = e · B*_up_, and *I*_up_ = *P_T_/e* (see also Appendix S5).

The biomass production by the basal level *P_B_* varies only little, as this quantity is completely determined by the interaction with the free nutrients (see Appendix S6). This property lies at the basis for explaining the increase in top biomass and food web efficiency as functional diversity increases everywhere.

When Δ_*T*_ = 0, the absence of a diverse top trophic level creates a slight relative advantage for the fast growing species *I*_1_ (see Appendix Figure S7.1). Its effects on the basal level strongly depend on Δ_*I*_. For high Δ_*I*_, the fast growing *B*_1_ is heavily suppressed and the basal biomass is concentrated in *B*_2_, which is less edible for the prominent *I*_1_. For low Δ_*I*_ (i.e., *I*_1_ and *I*_2_ are functionally similar and less specialized), the dominant *I*_1_ can also graze significantly on the slow growing *B*_2_, which strongly promotes the fast-growing *B*_1_. The higher growth rate of *B*_1_ causes strong fluctuations of the basal biomass (Figure 4), which, in turn, leads to less efficient nutrient exploitation (Figure 3). Thus, for both low and high intermediate diversity, the basal level is unevenly exploited, which leads to a relatively high proportion of basal biomass being lost from the system, instead of being transferred up the food web (see also Appendix S5). The rather low basal biomass that is transferred to the intermediate level supports only a modest amount of intermediate biomass, and hence, a low biomass and biomass production on the top level, and a low food web efficiency.

In contrast, when the top level is highly diverse (Δ_*T*_ = *high*), the intermediate level is more evenly exploited, leading to a balanced presence of both intermediate species. In turn, this leads to an efficient exploitation of the basal level, especially when Δ_*I*_ is also high, which is reflected by high values of *(P/B)_B_* (Figure 5). Even though *P_B_* remains roughly the same (Appendix Figure S5.2, and Appendix S6), *B*_up_ is increased (Figure 5) which leads to a significantly higher intermediate biomass and biomass production (Figures 3, S5.2), and, ultimately an increase in biomass on the top level. This increase subsequently explains the increase in food web efficiency through an increased top biomass production (Figure 5).

### Relative importance of parameters

The Random Forest model provides an estimate for the importance of each of the food web parameters in predicting the outcome (see Methods). Figure 6 shows them for the different biomasses and *CV* s for each of the 14 model parameters (for the relative importance of the different production metrics, see Appendix Figure S5.4).

**Figure 6:**
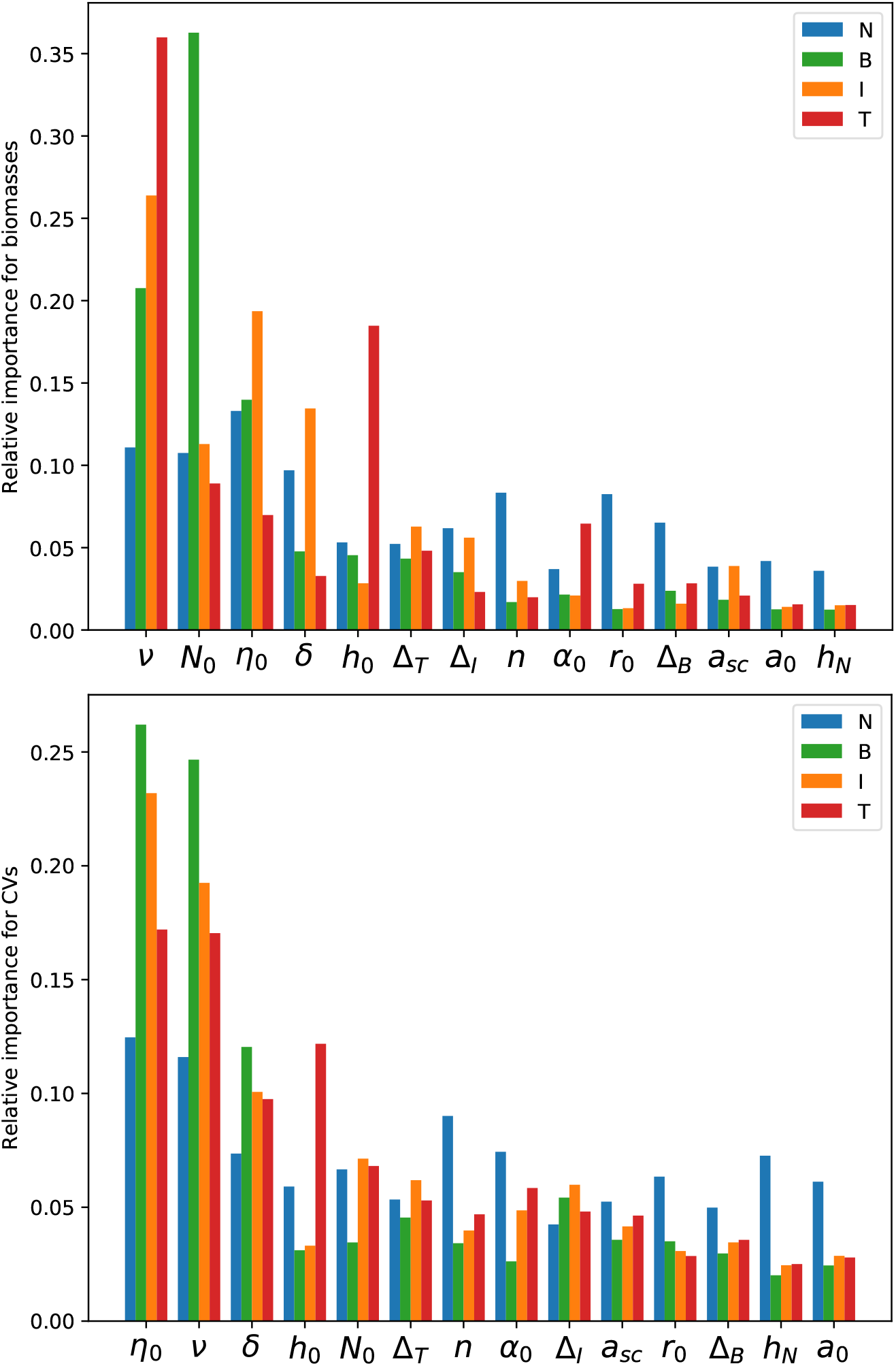
Relative importance of the different model parameters (see Table 1 on determining the biomasses and CVs of the different trophic levels. The relative importance quantifies how important the value of a certain parameter is to accurately predict the desired quantity, and they sum up to 1. The higher the relative importance of a parameter, the more relevant it is to make a prediction. In these graphs, the model parameters are ordered by their mean importance for each group of quantities (biomasses and CVs); for each parameter, the individual bars are ordered by trophic level.

The parameters in each graph are grouped by their mean importance in descending order. For example, the Hill-exponent of the functional response describing the intermediate-top interaction (*ν*) has the highest mean relative importance for predicting the biomasses on each trophic level (Figure 6, top). In particular, it is very important for predicting the biomass on the top and intermediate level. On the other hand, the nutrient-uptake half saturation constant *h_N_* is the least important.

One important observation in all three panels is that while the three possible trait differences Δ_*B*_, Δ_*I*_, and Δ_*T*_ have a strong influence on all the different quantities we have investigated (see Figs. 3–5), they are never amongst the most important parameters. However, this is not very surprising given the nature of the other parameters in our model: for example, it is very natural that increasing the nutrient inflow concentration *N*_0_ has a very strong influence on species’ biomasses.

Our results also show a balance between the relative importance of parameters affecting the external environment (such as the nutrient inflow concentration *N*_0_ and the inflow rate *δ*), and internal parameters affecting the ecological dynamics within the food web (such as the handling times *h*_0_*, η*_0_, and Hill-exponents *n, ν*). Remarkably, parameters affecting the intermediate-top interaction (*ν*, *η*_0_, *α*_0_) are of higher importance than their intermediate-basal analogues (*n*, *h*_0_, *a*_0_). In particular, the importance of the different diversity measures Δ_*T*_, Δ_*I*_, Δ_*B*_ is ranked by trophic level. In this way, it is clear that food web parameters affecting the top level of are of highest importance.

## Discussion

The food web model analyzed in this manuscript was built with the aim of being as general as possible, while still being ecologically realistic. Given the expansive range of different environmental and ecological situations that are effectively covered by the model, we did not intend to answer research questions about specific environmental or ecological conditions. Rather, we focused on how the *average* behavior of tritrophic systems depends on the diversity of each trophic level separately. In particular, we studied how functional diversity in tritrophic food webs affects the biomass distribution, temporal variability, and production, on average. The partial dependence graphs provided by training a Random Forest model on our data served as an ideal tool to answer these questions. Given the large amount of parameters that were randomly sampled, it is to be expected that the output data contains a large amount of variation. For example, parameters like the inflow nutrient concentration *N*_0_, or the inflow rate *δ* naturally have a very strong influence on the trophic level biomasses and temporal variation. Partial dependence graphs revealed how the predicted outcome changes as a function of one particular parameter, *on average*, i.e., independently of all other model parameters.

### Absence of coexistence in some webs

In two of the food webs we investigated, Τ and BΤ (see Figure 1), coexistence of all species was extremely rare (see Appendix S4). In almost every case, one of the two top species outcompeted the other one, as expected when applying the competitive exclusion principle (Hardin, 1960; Armstrong and McGehee, 1980; Klauschies and Gaedke, 2019). For less than 0.1% of the parameter combinations, both top species still co-occurred at the end of the simulation time. The structure of these two food webs entails that the two top species are competing with each other for only one resource, *I*, with no other density dependent interaction.

In contrast, coexistence of all species is very likely in the I and IΤ food webs, even though the two intermediate species also share a single resource, *B*. This is due to an additional density dependent interaction acting on the intermediate species, by the presence of the top level (which may or may not be diverse). Therefore, more than one species can exist at the intermediate level without the necessity of fine-tuning their interaction parameters (Huntly, 1991; Brose, 2008; van Velzen, 2020).

Viewed in this way, it is clear that the amount of functional diversity of one trophic level can drastically influence that of other trophic levels: a loss of functional diversity at the intermediate level in the IΤ or BIΤ food webs leads to a loss of functional diversity at the top level as well. It is therefore crucial to safeguard functional diversity of lower trophic levels to enable a diverse top predator community.

### Relative parameter importance

The random forest model trained on the output data of our simulations (see Methods) provides information on which of the input parameters (see Table 1) are most important for estimating the predicted biomasses, CVs, and production metrics. In short, a parameter is of high importance when it tends to appear high up in many different trees in the forest. Conversely, when a parameter only appears near the end of the trees, it is of low importance in estimating the desired outcome. These relative importances are ranked from highest to lowest in Figure 6.

Remarkably, parameters directly affecting the top trophic level tend to be of high importance, whereas parameters influencing the nutrient uptake by the basal species are all situated near the bottom end. The different diversity indices Δ_*B*_, Δ_*I*_, and Δ_*T*_ are also ranked by trophic level. This hierarchy shows how important the higher trophic levels are in determining the biomass distributions, temporal variation of biomass dynamics, and energetics of whole food webs. Our model is thus able to mechanistically support the general observation that changing the diversity of the top trophic level often has far-reaching consequences (Ripple et al., 2014).

In addition to most of the parameters governing the trophic interactions of the top trophic level, the nutrient inflow rate *δ* and concentration *N*_0_ are also of high importance. As *δ* determines the death rates of all the species in the model (see Eq. 8), and in particular those of the top level, it has a strong influence on the quantities we have investigated (Kath et al., 2018). The nutrient inflow concentration is unsurprisingly also of high importance in estimating these quantities. Its level, representing the total biomass carrying capacity of our system, affects the basal trophic level most strongly (Figure 6), which is in line with field observations (Gaedke, 1998).

This analysis shows that the relative importance measures provided by the random forest model provide useful information to uncover the underlying mechanisms that govern the dynamics of more complex models. Our results clearly show how external and internal food web parameters do not overpower each other. Information on both types is required for accurately predicting biomasses, biomass variability and food web energetics.

### The complex relationships between diversity and ecosystem functioning

Our results show that functional diversity robustly increases biomass and production efficiency (Figure 5) at high trophic levels (Figure 3), and generally decreases temporal variation (Figure 4), as summarized by Table 2. In addition, we reveal intricate and complicated interactions between the degree of diversity at different trophic levels and these ecosystem functions. These interactions complicate comparison of the numerous studies on the links between diversity and functioning to each other (Filip et al., 2014; Wohlgemuth et al., 2017; Flöder et al., 2018; Daam et al., 2019).

For instance, our model shows that the effect of increasing producer diversity on the biomasses of each trophic level highly depends on the amount of functional diversity of the other trophic levels (Figure 3). When the top level is not functionally diverse (Δ_*T*_ = 0), the direction of the effect of Δ_*B*_ on the basal biomass is determined by the amount of functional diversity of the intermediate level (Δ_*I*_). When Δ_*I*_ is low (low functional diversity), basal and intermediate biomass tend to decrease with increasing Δ_*B*_, whereas this trend reverses as Δ_*I*_ becomes higher. A recent experimental study revealed that the effects of producer diversity on food web functioning also depend on the trait values on the consumer level in a bitrophic system (Wohlgemuth et al., 2017). Our results indicate that this interdependency is of a very general nature, and moreover, is expected to hold for higher trophic levels as well, which are less manageable in experimental settings. Indeed, our model shows a similar pattern when investigating the effect of Δ_*I*_ and Δ_*T*_ on the intermediate and top biomasses. Starting from Δ_*T*_ = 0, increasing Δ_*I*_ leads to a reduction in intermediate biomass, compared to an increase in intermediate biomass when Δ_*T*_ is high. Our tritrophic food web comparison also shows that, when functional diversity is increased everywhere, the biomass of the intermediate and top species increases significantly, whereas the basal biomass stays roughly constant. The same pattern was found in a modeling study comparing food webs of up to 100 animal species (Schneider et al., 2016). This correspondency gives credibility to considering the effects of biodiversity on food web functioning through changing the functional diversity in simpler food webs, instead of changing the species number which significantly increases food web complexity.

The effect of functional diversity on the temporal variability (*CV*) of the biomasses at the different trophic levels also exhibited a complex dependency on the functional diversity of every single trophic level (Figure 4). One particularly robust result, however, is the non-monotonous relationship between top diversity (Δ_*T*_) and the *CV* of any trophic level. When Δ_*T*_ is increased from 0 to *low*, the *CV* s tended to strongly decrease. Such a reduction in the *CV* of a community as diversity increases has often been observed (Tilman, 1996), and can often be attributed to the presence of compensatory dynamical patterns (Gonzalez and Loreau, 2009; Bauer et al., 2014). However, as Δ_*T*_ is increased further from *low* to *high*, the *CV* of each trophic level increased again. Hence, additional mechanisms governing the dynamics must also have a strong influence of the trophic level *CV* s. In Ceulemans et al. (2019), we observed a similar pattern in the trophic level *CV* s, which could be explained by the increased relevance of an additional dynamical timescale at high Δ_*T*_: the biomasses not only varied rapidly within predator-prey cycles, but also due to slower trait changes. As this slower timescale became more dominant, the *CV* increased again. Due to the similar model structure, this mechanism may be responsible for the increase in *CV* here as well. This result suggests that mechanisms for dampening community temporal variability established for simple but functionally diverse systems, such as compensatory dynamics arising from competition for a joint resource, may be counteracted by destabilizing effects in more complex—and thus more realistic—systems.

Examining how the functional composition at each trophic level and ecosystem functions are linked allows us to mechanistically understand why the biomass and biomass production on higher trophic levels is maximal when every trophic level is diverse, and why the diversity of the top level plays such a crucial role. This becomes particularly obvious when comparing the trends of the different metrics related to biomass production within the food web (see Results, Figure 5, and Appendix Figure S5.4).

A functionally diverse consumer community leads to an efficient exploitation of the production at the prey level due to their functional complementarity (Gamfeldt et al., 2005). In our model, this mechanism is present between both the top and intermediate, as well as between the intermediate and basal level: a diverse top community efficiently exploits the intermediate production, which in turn results in the basal production being efficiently exploited. In contrast, when the top community is not functionally diverse, potentially functionally diverse intermediate and basal communities adjust in species composition so that they escape efficient predation (Filip et al., 2014; Seiler et al., 2017). As a consequence, a higher proportion of the production is lost from the system by non-grazing mortality rather than transferred up to the level above. In this way, the effects of functional diversity of different trophic levels synergize to make the food web with diversity everywhere the most efficient configuration in transferring biomass from the basal to the top level (Figure 5, *P_T_/P_B_*). Importantly, even though the trade-off structure in our model is comparatively simple, analysis of the individual populations’ biomasses (Figure S7.1, Appendix S7) confirms that they function as intended and prevent any one species from dominating others on average, thus providing additional evidence that the patterns we observe are caused by changes in trait differences between the species on each trophic level.

The importance of considering multitrophic diversity has been emphasized before (Gamfeldt et al., 2005; Filip et al., 2014; Soliveres et al., 2016; Lefcheck et al., 2015; Barnes et al., 2018; Ceulemans et al., 2019). With these complex interactions between functional diversity of different trophic levels clearly exhibited by our model, it is not surprising that studies focusing on a single food web structure or a single parametrization sometimes find incommensurable results. For example, increased primary producer diversity had often been linked to an increased producer biomass and production (Tilman et al., 1997; Cardinale et al., 2011). Our results show that this relationship not only depends on the trait values of the consumer level (Seabloom et al., 2017; Wohlgemuth et al., 2017), but crucially also on the top level. Hence, we reveal considerable variation in the behavior of differently structured food webs with respect to the relationship between diversity and ecosystem functioning. Nevertheless, we are able to identify clear trends and uncover mechanisms governing the behavior of tritrophic systems, even when considering a large range of different parameter combinations.

## Concluding remarks

Understanding the link between functional diversity and the functioning of complex food webs is crucial to accurately predict how losses in functional diversity will affect the functions of natural food webs everywhere around us. Considerable detailed knowledge about this link has been gained in communities comprising of one or two trophic levels. Partly, the knowledge gained from bitrophic systems helps to understand tritrophic ones, such as the enhanced exploitation of resources by a more diverse consumer community. However, accounting for the third trophic level clearly shows that a restriction to two trophic levels may yield misleading results for complex natural food webs. The present comparison of several different food webs consisting of three trophic levels (see Figure 1) reveals simultaneously operating bottom-up and top-down cascading effects over three trophic levels. At high functional diversity throughout the whole food web, functional shifts within the individual trophic levels result in a high food web efficiency and biomass on higher trophic levels, and a high degree of nutrient exploitation. Additionally, we show that the functional diversity on the top level is a strongly regulating factor for the biomass, temporal variability, and biomass production efficiency of any trophic level. Therefore, to prevent drastic reduction of important functions, as well as potentially irreversible transitions, it is of crucial importance to increase our efforts in conserving diversity of higher trophic levels, despite the often large operational problems involved.

## Acknowledgements

We thank Mridul Thomas for introducing us to using Random Forest Models on complex datasets, and Toni Klauschies and Markus Stark for interesting discussions of our results. We would also like to thank Stefanie Moorthi, Sabine Flöder, Peter de Ruiter, Laurie Wojcik, and two anonymous referees, for giving valuable feedback on an earlier version of the manuscript.

