## Supplementary material for "Top predators govern multitrophic diversity effects in tritrophic food webs": S1

### S1 Meaning and effect of the cross link scaling parameter

When functional diversity is present on adjacent trophic levels, as is the case for the BI, IT, and BIT food webs, there exist cross links between the species on the different sides of the web. In our model, the relative strength of these cross links can be varied independently of the amount of functional diversity through changing the relevant attack rates by multiplying them with $a_{\text{scale}}$. For higher $a_{\text{scale}}$, the cross links get relatively weaker.

The interaction between two species via a Holling-Type-III functional response is determined by both handling time ($h$ or $\eta$) and attack rate ($a$ or $\alpha$). In our model, trait differences affect the attack rates such that increasing trait differences on adjacent trophic levels may lead to increased specialization of the consumer species on their prey through reduction of the cross link strengths. Explicitly:

$$a = a_0 \begin{pmatrix} 1 & a_x \\ a_x & 1 \end{pmatrix}, \qquad \alpha = \alpha_0 \begin{pmatrix} 1 & \alpha_x \\ \alpha_x & 1 \end{pmatrix}, \tag{S1.1}$$

with

$$a_x = a_{\text{scale}}{}^{-\min(\Delta_B, \Delta_I)}, \qquad \alpha_x = a_{\text{scale}}{}^{-\min(\Delta_I, \Delta_T)}, \tag{S1.2}$$

such that this reduction depends on the minimal trait difference of the relevant trophic levels, and the feeding links are unaffected when one of the trait differences is equal to zero (Figure S1.1).

In a way, the cross link scaling parameter $a_{\text{scale}}$ affects the structure of the different food webs we have compared (see also Figure 1, main text). As the ratio between the parallel and the cross links in those food webs where diversity is present on adjacent trophic levels (i.e. BI, IT, and BIT) is varied, the food web structure changes from either a strongly linked ($a_{\text{scale}} \approx 1$), to two weakly connected chains ($a_{\text{scale}} \gg 1$).

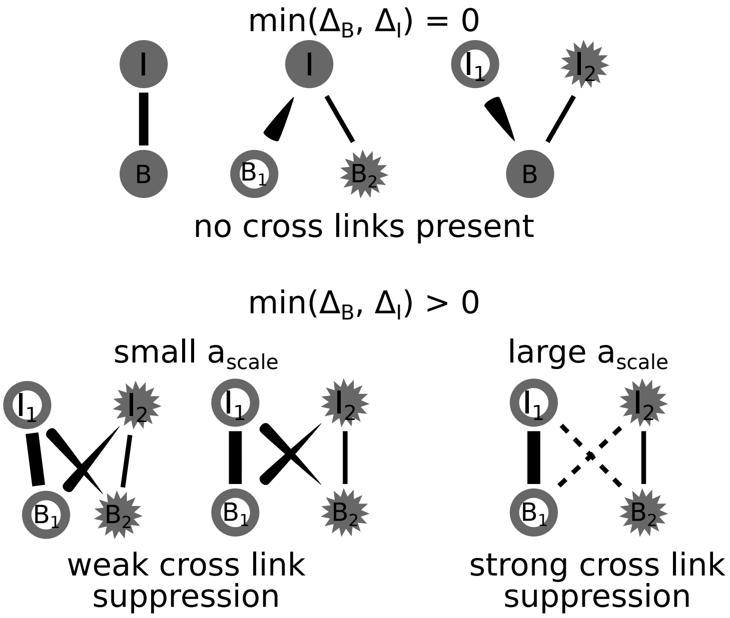


Figure S1.1: Pictorial representation of the effect of $a_{\text{scale}}$ on the food web structure. Only effects on the basal and intermediate level are shown for simplicity, but the interactions between the intermediate and top level is parametrised equivalently. When either $\Delta_B = 0$ or $\Delta_I = 0$ (top row), there are no cross links present and $a_{\text{scale}}$ has no effect. It is only when both $\Delta_B$ and $\Delta_I$ are positive that cross links appear between the basal and intermediate level in the food web. In this case, when $a_{\text{scale}}$ is small (close to 1), the cross links $B_1 - I_2$ and $B_2 - I_1$ are not significantly suppressed. Contrarily, for large $a_{\text{scale}}$ the trophic links between $B_1, I_2$ and $B_2, I_1$ are strongly suppressed, as indicated by the dashed lines.
