## Supplementary material for "Top predators govern multitrophic diversity effects in tritrophic food webs": S2

### S2 Details of the allometric properties

Allometric scaling entails that species' growth rates are linked to their body mass. In particular:

$$\frac{\text{maximal growth rate of trophic level } k+1}{\text{maximal growth rate of trophic level } k} = \left[\frac{m_{k+1}}{m_k}\right]^{\lambda} \tag{S2.1}$$

where $k \in \{1,2\}$, $m_k$ is the body mass of the species on level $k$, and $\lambda$ the allometric scaling exponent. As mentioned in the main text, the centers of the intervals for which the maximal growth and grazing rated are selected corresponds to a food chain with biomass ratios between adjacent trophic levels on the order of $10^3$, and $\lambda = -0.15$. However, assuming the same scaling exponent $\lambda$, the body mass ratio can vary from approximately 1 to approximately $10,000,000$ at the extreme cases for either maximal prey and minimal predator growth rates, or minimal prey and maximal predator growth rates, respectively.
