## Supplementary material for "Top predators govern multitrophic diversity effects in tritrophic food webs": S3

### S3 Detailed description of data production and analysis procedure

The goal of our study is to gain understanding in the general behaviour and ecological patterns that can be expected in tritrophic food webs, as a function of their diversity. Any serious attempt at studying these general properties cannot be limited to a certain small subset of the parameter space. On the other hand, there is a practical limit in the extent to which the whole ecologically parameter space can be explored.

In addition, fully capturing the behaviour of complex dynamical systems, such as ours, requires going beyond linear stability analysis. Due to modelling the interactions between species as non-linear generalised Holling-type-III functional responses, the resulting population dynamics will often relax to complex limit cycles or chaotic attractors.

As described in the main text, we balanced these limitations by adopting a two-step process. First, we numerically integrated the ordinary differential equations (ODEs) describing our food webs (cf. Eq. (1), main text) for $128,000$ parameter combinations, and saved aggregate properties of the resulting dynamics (the population and trophic level biomasses and Coefficients of Variation (CV)). This dataset was subsequently used for training a Random Forest model, in order to be able to predict these quantities for many more parameter combinations for which the ODEs were not solved.

In this Appendix, we will describe in detail what kind of data we have used for our results, and how it was produced. Our whole procedure can be split up into three main parts: producing the training dataset, training the Random Forest, and producing the Partial Dependence plots (Figs. 3, 4 and 5 in the main text, and others in the Supplementary Materials).

#### S3.1 Producing the training dataset

As described above, the starting point for obtaining our results is the set of ODEs (cf. Eq. (1), main text). These equations can be used to describe any of the food webs shown in Figure 1 (main text), through their explicit dependence on the different trait difference parameters $\Delta_B, \Delta_I \,\&\, \Delta_T$.

The only way to gain some insight in the behaviour of the solutions describing the population dynamics is to solve these equations. Other techniques such as calculating the fixed points and analysing their stability through the eigenvalues of the Jacobian matrix are not suitable, because of the high degree of non-linearity in the terms that define the interactions between the different populations in the food web (the generalised Holling Type-III functional responses). Due to these non-linear interaction terms the population dynamics frequently do not relax to a simple stable point, but rather to more complex attractors, such as limit cycles or chaotic attractors.

Figure S3.1 provides a schematic overview of the different steps that were taken to produce the data on which the Random Forest model will later be trained.

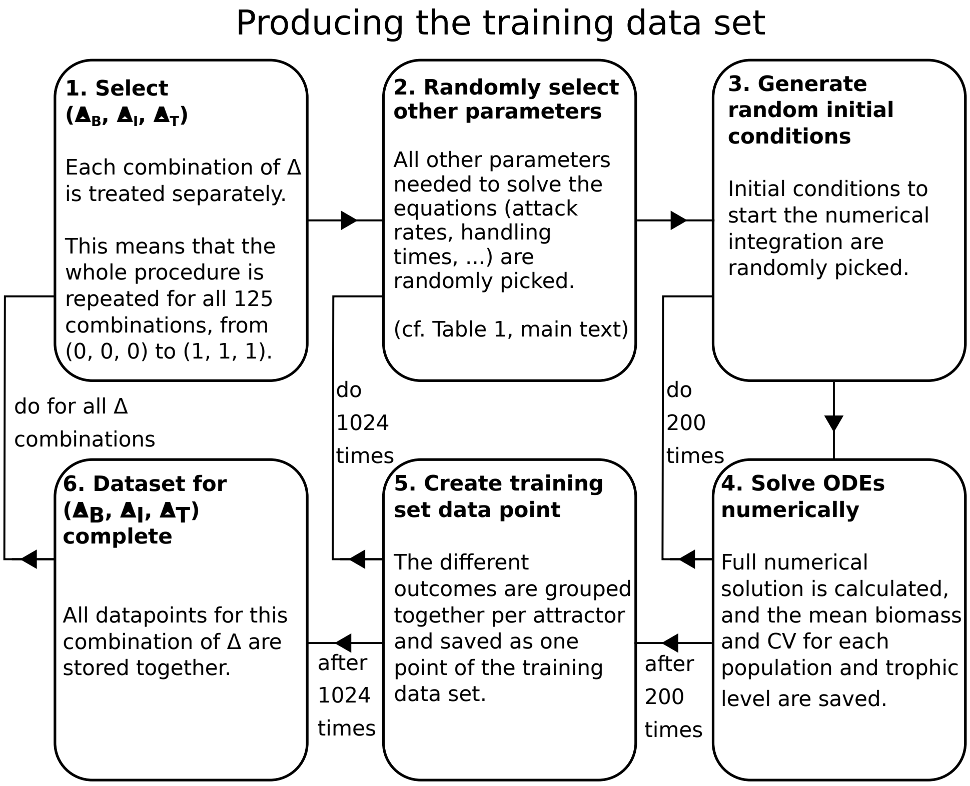


Figure S3.1: Schematic overview of the steps required to produce the training data set. See the text for more details on each step.

1. **Select** $(\Delta_B, \Delta_I, \Delta_T)$: While Figure 1 (main text) shows the different food webs ordered by the presence or absence of functional diversity on each trophic level, it can also be helpful to think about a different partitioning of all possibilities. Particularly, one could imagine ordering the food webs simply by their values of $\Delta_B$, $\Delta_I$, and $\Delta_T$. In that case there are not eight, but 125 different possibilities, since each $\Delta_i \in \{0, 0.25, 0.5, 0.75, 1\}$.

   It is then for each of these 125 cases that the dynamics of 1024 randomly sampled parameter combinations will be investigated.

2. **Randomly select other parameters**: The other eleven parameters required to solve the ODE model are randomly sampled from well defined intervals (see Table 1, main text).

3. **Generate random initial conditions**: Because of the increased complexity of the food webs we describe, it is possible that there is a strong dependence of the outcome of the ODE model on the initial conditions. Several different scenarios are possible: any particular initial condition may not lead to coexistence of all species in the food

web, or, there may be multiple attractors for which all species do coexist. It would thus be highly inaccurate to assume that one single initial condition is representative for all possible outcomes of this particular parametrisation. Therefore, for each parameter combination, it is additionally necessary to record the outcomes of several different initial conditions. To obtain the datasets used to train the RF model in the main text, we recorded the outcomes of 200 randomly sampled initial conditions per parameter combination. As described in the main text, these initial conditions were sampled in a way that the total biomass of each trophic level does not exceed twice its theoretical maximum, in order to remain ecologically reasonable and prevent very strong oscillations when the system is relaxing to its attractor.

4. **Solve ODEs numerically**: Now that all parameters have been assigned a value, and an initial value has been set, numerical integration of the ODE model is performed. To prevent any influence from the potential presence of very slow transient dynamical patterns, the system is allowed to relax to its equilibrium state for $100,000$ time units. After this time, the mean biomass of each population, as well as for each trophic level as a whole, is calculated over $30,000$ time units. Similarly, the standard deviation of the dynamics is calculated for each individual population and each trophic level, from which the coefficient of variation can be computed ($CV = \sigma/\mu$, where $\sigma$ is the standard deviation and $\mu$ the mean). For schematic details of this procedure, see Figure S3.2. A real example timeseries from our model is shown in Figure S3.3.

5. **Create training set data point**: After the mean biomasses and CVs of 200 unique randomly sampled initial conditions have been calculated, all the numerical simulations for one parameter combination have been completed. However, typically, many initial conditions will relax to the same attractor. It is therefore very useful to group the different outcomes together per attractor: when *all* of the different species' means and $CV$s are sufficiently close to each other (absolute tolerance of 5 for the means, 0.01 for the $CV$s, and relative tolerance of 0.1), the initial conditions are assumed to relax to the same attractor.

   The structure of a data point belonging to one parameter combination is the following:

```
dict; # Python dictionary containing all parameter values
num_attr; # number of different attractors
ext; # whether any species are extinct or not on each attractor
TL_B; # total biomass on each trophic level
odd_B; # biomass of B1, I1, T1
ind_CV; # CV of each population
TL_CV; # CV of each trophic level
attr_prop # proportion of initial conditions relaxing to each attractor
```

   A typical data point:

```
{'B_diff': 0.75, 'I_diff': 0.75, 'T_diff': 0.25, 'N0': 666.210112394016,
'h_N': 18.91630737427768, 'r0': 1.8743678849645211,
'a0': 0.0015176450564662867, 'h0': 0.79237825772891,
'alpha0': 0.0003298639600209292, 'eta0': 3.209067776129767,
```

```
'delta': 0.05710530459267592, 'n': 1.907283906197544,
'nu': 1.3633537312107176, 'a_scale': 1.2903979398824104, 'id': 2824000};
2;
[0,1];
[[100.677904  ,168.3763992 ,198.2327592 ,233.3333859 ],
 [535.659272  , 19.29489409,268.8575037 ,  1.64734166]];
[[105.72043   ,170.586387 ,168.072246 ],
 [ 14.6636983, 59.3558927,  0.        ]];
[[0.61712753,1.32296074,0.47741629,0.04292298],
 [0.        ,0.        ,0.        ,0.        ]];
[[1.42259412,0.51964668,0.04594913],
  [0.        ,0.        ,0.        ]];
[[1.19062106,0.21908273,0.03512972],
  [0.        ,0.        ,0.        ]];
[0.83,0.17]
```

This data point has $\Delta_B = 0.75, \Delta_I = 0.75, \Delta_T = 0.25$, and thus belongs to the BIT food web and there are two different attractors found. 83% of the initial conditions relaxed to the first one listed, for which all species coexist, whereas 17% relaxed to the second one listed, which has only one species on the top level (as the biomass of $T_1$ equals zero). Additionally, one can see that the second attractor listed must be a stable fixed point (due to the $CVs$ being zero), whereas the first attractor cannot be a stable fixed point and must therefore be a limit cycle or a more complex type of attractor. Investigation of the actual timeseries confirms that the attractor is a limit cycle in this case (Figure S3.3).

6. **Dataset for** $(\Delta_B, \Delta_I, \Delta_T)$ **complete**: After 1024 randomly sampled parameter combinations have been investigated (each for 200 initial conditions), and thus 1024 data points have been added to the training data set, all the $ODE$ simulations for this combination of $\Delta_B$, $\Delta_I$, and $\Delta_T$ have been completed.

This whole process is now repeated for the next combination of trait differences until there are 1024 data points for each combination. This means that the ODE model described in Equation (1) (main text) will have been solved $125 \cdot 1024 \cdot 200 = 25,600,000$ times. If we assume that each individual integration takes just one second, this requires just under 300 *days* of total integration time. Luckily, this whole process is easily parallelizable, which can substantially reduce the required time, depending on the amount of available computing cores.

### S3.2 Training the Random Forest

The next step required to produce our results is to train a Random Forest (RF) on the training dataset, produced by solving the ODE model millions of times and recording the mean biomasses and coefficients of variation (CVs), as described above. There are two main benefits of using a RF to analyze and present our data. Firstly, assuming that the RF model predictions are accurate, the ability to predict the output value of parameter combinations that are not in the training dataset increases the confidence level of any trends that can be identified (see Figure S3.5). Secondly, the RF model provides us with a measure of the importance of each of the input parameters (see Table 1, main text) in determining the output value (see Figure 6, main text).

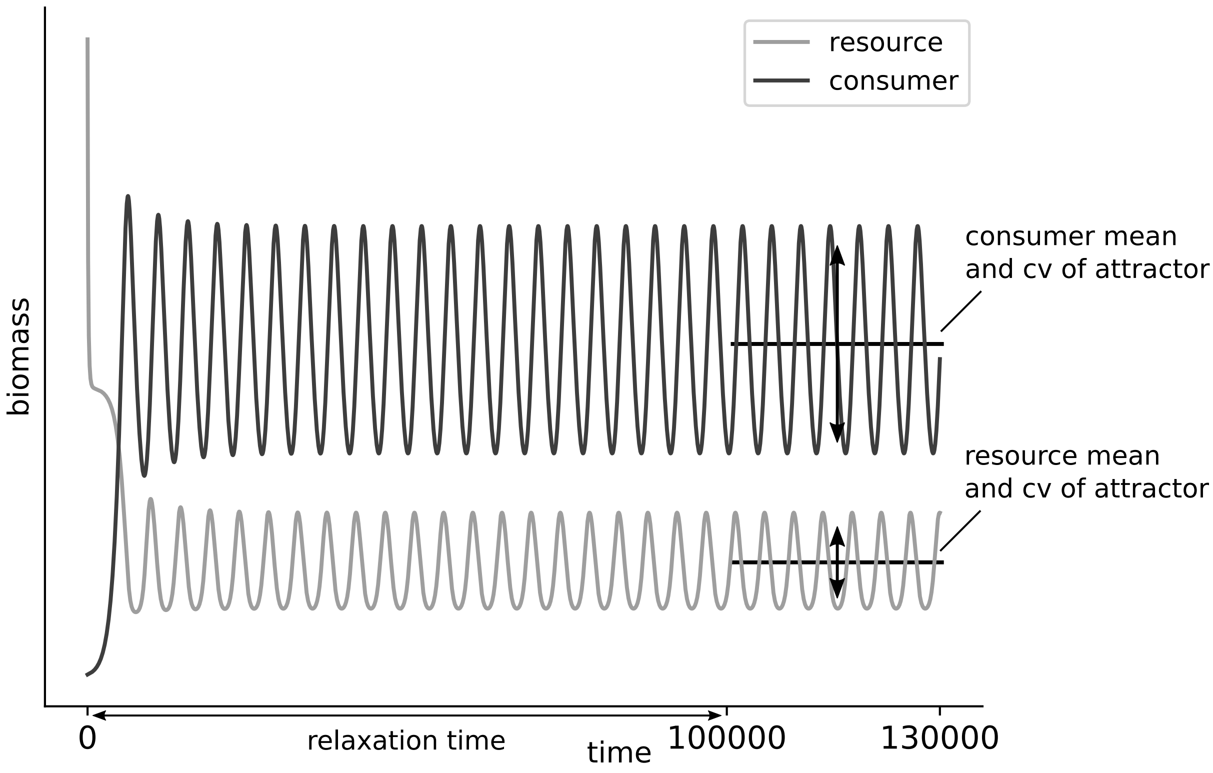


Figure S3.2: Schematic example of a trajectory starting from a random initial condition, for a simple consumer-resource system. To estimate the mean biomasses and biomass $CV$s on the attractor, they are calculated over a period of $30,000$ time units, after the system has been allowed to relax to the attractor for $100,000$ time units.

#### S3.2.1 Short overview of how a Random Forest works

The goal of any model is to estimate each output value $y_i$ ($i \in \{1, \dots, n\}$, where $n$ is the number of data points) as well as possible by constructing a function $f$, for which

$$\hat{y}_i = f(\boldsymbol{x}_i) \tag{S3.1}$$

where $\boldsymbol{x}_i = (x_{i,1}, x_{i,2}, \cdots, x_{i,p})$ are the input variables (or features), with $p$ the amount of features, such that $\hat{y}_i$ is the model estimate of $y_i$.

Random Forest Models are a class of machine learning models, which are popularly used due to their relatively simple structure and high versatility. In a random forest, each $\hat{y}_i$ is calculated by averaging the predictions of many different regression trees. Each individual regression tree has a certain degree of randomness: only a random subset of the data (~ 63%) is used to calculate the tree, and at each node only a randomly selected subset of the data features is taken into account. By averaging the predictions of a large amount of such quasi-randomly generated regression trees, a random forest model can make more accurate predictions than a single optimised regression tree (Breiman, 2001).

Because each individual tree in the forest has only been trained on a subset of the data, the remaining data points can be used to estimate the accuracy of that tree. This process is called Out-Of-Bag (OOB) error estimation, and gives a measure for the accuracy of the model:

$$\mathrm{OOB}_{\mathrm{score}} = 1 - \frac{\mathrm{MSE}_{\mathrm{OOB}}}{\hat{\sigma}_y^2}, \tag{S3.2}$$

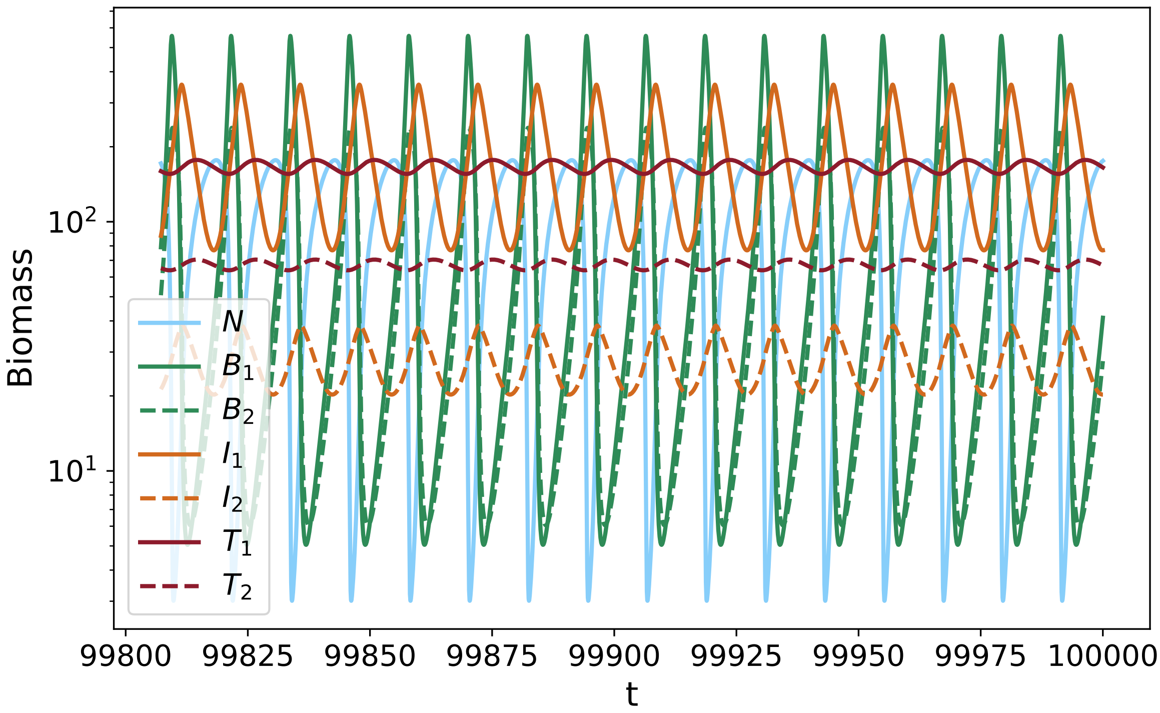


Figure S3.3: Example timeseries of the dynamics possible in our model. This timeseries was created using the parameters given in the example datapoint above. It can clearly be seen that the attractor is a limit cycle, which means that the dynamics will oscillate in this way perpetually.

where $\hat{\sigma}_y^2$ is the variance estimator of the outcome variable $y$, and

$$\mathrm{MSE_{OOB}} = \frac{1}{n}\sum_{i=1}^{n}\left(y_i - \bar{y}_i^{\mathrm{OOB}}\right)^2, \tag{S3.3}$$

with $\bar{y}_i^{\mathrm{OOB}}$ being the mean OOB predicted value of $y_i$. In this way, an OOB score of 1 signifies that the model is able to predict the outcome perfectly. Using the OOB error estimation, it is also possible to evaluate the importance of each of the input parameters in predicting the output value. This is done by quantifying the change in the random forest's accuracy when the input parameter's values are permuted. A large change in accuracy indicates a high importance in predicting the correct value, and vice-versa.

For each quantity of interest (see Results, main text), an Extremely Random Forest consisting of 2000 trees was trained using the Scikit-learn (Pedregosa et al., 2011) package in Python. Importantly, each forest was only trained on the subset of parameter combinations for which coexistence was found to be possible (see also Appendix S4). In case multiple attractors were found for which all species were in coexistence, the weighted average of the different outcomes was used.

#### S3.2.2 How Random Forests were used to analyse our data

For extra clarity, let us apply the general equation (S3.1) to a specific example: the biomass produced by the top trophic level ($P_T$). The procedure for using the RF model to predict any of the other quantities presented in the main text and appendices is similar.

As described above, the goal of the RF algorithm is to be able to accurately predict the output values $y_i$. In this example, let $y_i = P_{Ti}$, such that $P_{Ti}$ denotes the biomass production on the top level for the $i^{\text{th}}$ data point. This quantity can easily be calculated using the biomasses on the top trophic level and the relevant input parameters (see Appendix S5 for a more elaborate explanation). The input information that the RF algorithm will use to make these predictions is the collection of parameters we have varied to produce the data. Explicitly, the RF algorithms will thus estimate the function $F$ for which:

$$y = F(\boldsymbol{x}) \Leftrightarrow \forall i : y_i = F(\boldsymbol{x}_i), \tag{S3.4}$$

with

$$y_i = P_{Ti} \quad \text{and} \quad \boldsymbol{x}_i = [\Delta_{Bi}, \Delta_{Ii}, \Delta_{Ti}, N_{0i}, h_{Ni}, r'_{0i}, a_{0i}, h_{0i}, \alpha_{0i}, \eta_{0i}, \delta_i, a_{\text{scale},i}, n_i, \nu_i] \tag{S3.5}$$

describing the predicted value and all the relevant parameters that belong to it. These two arrays contain all the information the RF algorithm needs to be able to estimate the function $F$, and calculate the *OOB* score and parameter importances.

### S3.3 Application of the Random Forest model to produce partial dependence plots

Now that the Random Forest (RF) has been trained on the training data, it can finally be used to gain an increased understanding of the system we set out to study. The parameter importances (see Figure 6, main text) are calculated during the training phase of the RF model, and thus are readily available for further analysis. In contrast, to construct the partial dependence plots (Figs. 3, 4 and 5 in the main text, and others in the Supplementary Materials), additional calculations using the RF model are necessary.

As explained above, by training the RF we have constructed a function with which we can predict the desired outcome quantity, using the input parameters used to train the model (cf. Eq. S3.1). Importantly, we can now use our model to make predictions for parameter combinations that are not in the training dataset, with which we can calculate the partial dependence (PD) of the quantities predicted by the RF on the different input parameters. In other words, the partial dependence of the RF prediction $f(\boldsymbol{x})$ on the input parameter $x_j$ ($j \in \{0, ..., p\}$ with $p$ the number of input parameters) is typically defined as $f_j(x_j)$ (Hastie et al., 2009):

$$f_j(x_j) = \frac{1}{n} \sum_{i=1}^{n} f(x_{i,1}, ..., x_j, ..., x_{i,p}), \tag{S3.6}$$

where $x_{i,k}$ is the value of the $k^{\text{th}}$ input parameter in the $i^{\text{th}}$ data point.[1]

A graphical explanation of calculating the partial effects in this way is shown in Figure S3.4, for $f_1$. For each value of $x_1$, the RF makes $n$ (number of data points) predictions, of which the median value is taken. These $n$ predictions are the predicted output values of all data points, where the actual value of $x_1$ in the data point has been substituted by the currently evaluated one. The procedure is then repeated for the next value of $x_1$. Using this definition, the degree of extrapolation necessary to calculate the partial effects is kept minimal.

[1] Note that, by this definition, $f_j$ does *not* denote the $j$-th component of $f$.

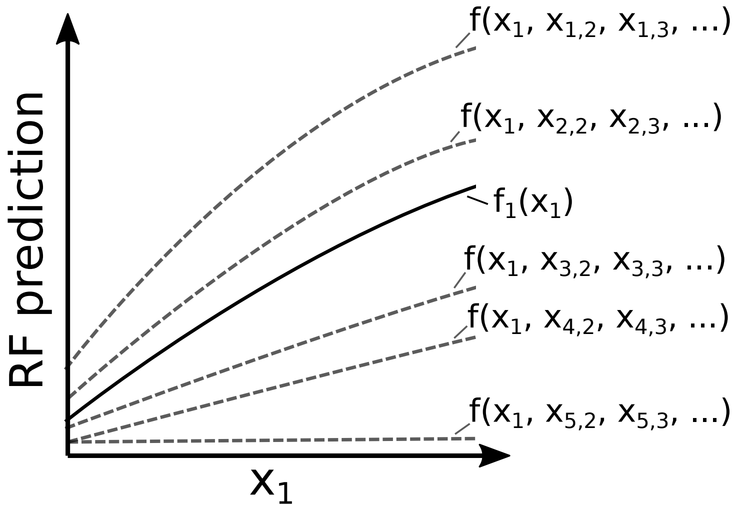


Figure S3.4: Schematic example of the construction of the partial dependence (PD) of the RF prediction on the input parameter $x_1$: $f_1$. See the text for a detailed explanation.

#### S3.3.1 Increased accuracy of estimate due to the Random Forest

The Partial Dependence Plots (PDPs) additionally allow for a clear visualisation of the benefit of using the RF trained on our data, as compared to simply using the data itself (Figure S3.5). In this graph, the 99% confidence intervals on the mean are estimated by $3\sigma/\sqrt{n_s}$, where $\sigma$ and $n_s$ denote the standard deviation and size of the sample, respectively.

The most important reason for the difference in size between in the error bars lies in the potentially huge difference between $n_s$ for the data itself as compared to what is predicted by the PDP. When using the data itself to estimate the mean top biomass for a given value of $\Delta_T$, $n_s$ is the number of data points in the training data set that actually have that value of $\Delta_T$ as an input. In general, this number will be only a fraction of the total dataset as there will be many other data points with different values of $\Delta_T$. However, for the PDP estimate, $n_s$ is always the total number of data points in the whole training data set. This is because, for a certain value of $\Delta_T$, the PDP predicts the top level biomass by substituting that value of $\Delta_T$ into every single data point in the training set. The error on the estimate of the mean top level biomass will therefore generally be much smaller. In Figure S3.5, the error on the estimate of the data itself is also rather small, because there are still many data points for each value of $\Delta_T$ in this one-dimensional division of the data. However, one can easily imagine partitioning the data into a much higher number of categories such that the number of data points in each category is much lower compared to the total number of data points in the training set (such as in the pseudo-three-dimensional PDPs shown in Figures 3-5 (main text) and similar ones in Appendices S5 and S7).

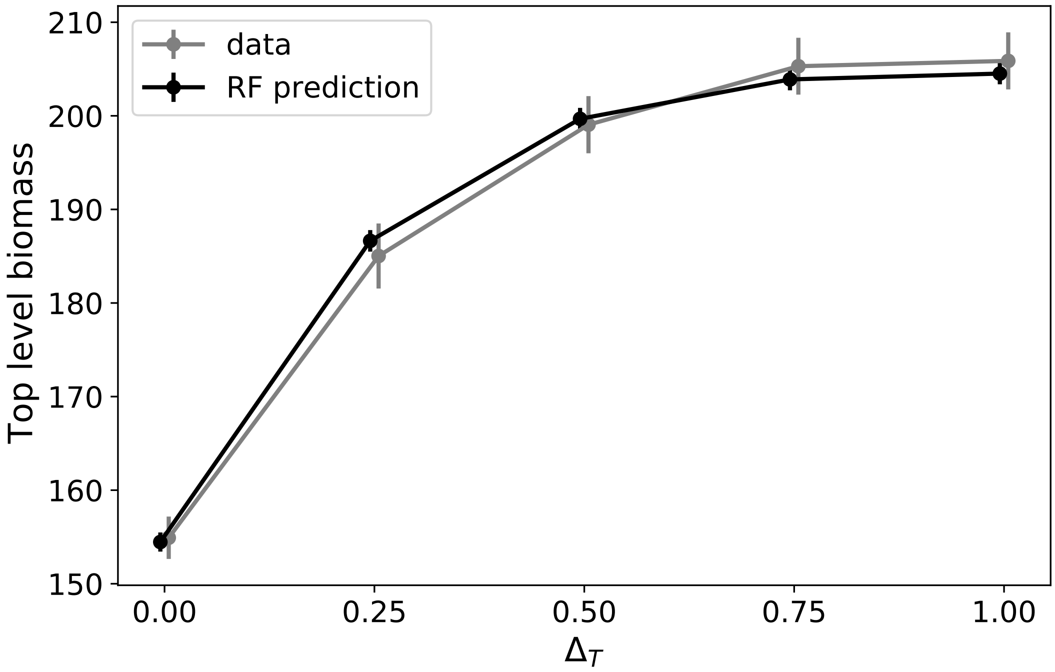


Figure S3.5: Partial Dependence Plot (PDP) of the total biomass on the top level on the trait difference on the top level $\Delta_T$ (black line), compared to what can be extracted from the data itself (gray line). The circles indicate the estimated mean biomass for each value of $\Delta_T$, and the error bars an estimate for for the 99% confidence intervals. One can see that the mean biomass on the top level is more accurately estimated by using the PDP provided by the RF, rather than simply estimating this quantity from the training itself.
