## Supplementary material for "Top predators govern multitrophic diversity effects in tritrophic food webs": S4

### S4 Proportion of coexistence of all species

All the results presented only apply to those parameter combinations that actually lead to coexistence of all species in the food web. In most cases, coexistence of all species was probable to very probable. However, for the T and IT food webs, coexistence of all species was very rare (only in resp. 1 and 8 cases, out of resp. 4096 and 16384). See the corresponding paragraph in the Discussion (main text) for an explanation of these results.

| Web | Coexistence proportion |
|---|---|
| chain | 0.945 |
| B | 0.931 |
| I | 0.936 |
| BI | 0.938 |
| T | < 0.001 |
| BT | < 0.001 |
| IT | 0.637 |
| BIT | 0.647 |
