## Supplementary material for "Top predators govern multitrophic diversity effects in tritrophic food webs": S5


### S5 On calculating the average biomass production

We calculated the mean biomass production and $P/B$ ratios from the mean biomasses of the the individual species and of the trophic levels per parameter combination and initial condition (Figure S5.1). See also Appendix S3 for a details description of how these quantities were obtained.

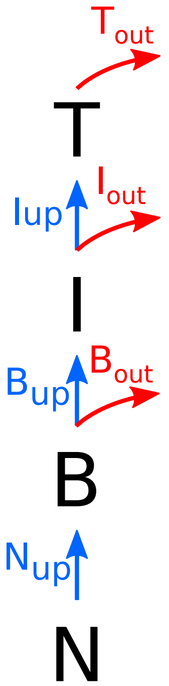


$$T_{\text{out}} = \sum d_{T_i} T_i$$

$$I_{\text{up}} \equiv \frac{T_{\text{out}}}{e} = \frac{\sum d_{T_i} T_i}{e}$$

$$I_{\text{out}} = \sum d_{I_i} I_i$$

$$B_{\text{up}} \equiv \frac{I_{\text{up}} + I_{\text{out}}}{e} = \frac{\sum d_{T_i} T_i}{e^2} + \frac{\sum d_{I_i} I_i}{e}$$

$$B_{\text{out}} = \sum d_{B_i} B_i$$

$$N_{\text{up}} \equiv B_{\text{up}} + B_{\text{out}} = \frac{\sum d_{T_i} T_i}{e^2} + \frac{\sum d_{I_i} I_i}{e} + \sum d_{B_i} B_i$$

Figure S5.1: Step by step illustration on how to calculate the biomass flows between trophic levels in our system. These equations hold for all of the different food webs we compared. When there are two species on a trophic level, the sums run over $i \in \{1, 2\}$; when there is only one, no summation is required. $N_{\text{up}}$ denotes the nutrient uptake by the basal species, and $B_{\text{up}}$ & $I_{\text{up}}$ the total biomass flowing upwards from the basal and intermediate level respectively.

The time-averaged mean biomass production $\bar{P}$ of each trophic level is then given by:

$$\begin{aligned}\bar{P}_T &= \bar{T}_{\text{out}} = \sum d_{T_i}\bar{T}_i \\ \bar{P}_I &= \bar{I}_{\text{up}} + \bar{I}_{\text{out}} = \frac{\sum d_{T_i}\bar{T}_i}{e} + \sum d_{I_i}\bar{I}_i \\ \bar{P}_B &= \bar{B}_{\text{up}} + \bar{B}_{\text{out}} = \frac{\sum d_{T_i}\bar{T}_i}{e^2} + \frac{\sum d_{I_i}\bar{I}_i}{e} + \sum d_{B_i}\bar{B}_i,\end{aligned} \tag{S5.1}$$

since all the loss rates $d$ are density-independent (cf. Eqs. (1) and (8), main text). In these equations, the bar is simply present to emphasise that our data consists of long-term temporal averages. The quantities $\bar{B}_i, \bar{I}_i$, and $\bar{T}_i$ are exactly the mean biomasses per population calculated by solving the ODE system (Equation (8)) and stored in the dataset (see also Appendix S3).

Shown below are the partial dependence graphs for some of these quantities, to support the mechanisms which yield increased biomass on the top level when diversity on all trophic levels is high. Importantly, these figures also support our claim that the random forests trained on our data are actually accurate: the equalities $P_T = e \cdot I_{\text{up}}$ and $P_I = e \cdot B_{\text{up}}$ hold throughout the whole range of $\Delta_B$, $\Delta_T$, and $\Delta_T$.

The quantities $P_T$, $B_{\text{up}}$, $(P/B)_B$, and $P_T/P_B$ are shown in Figure 5 (main text).

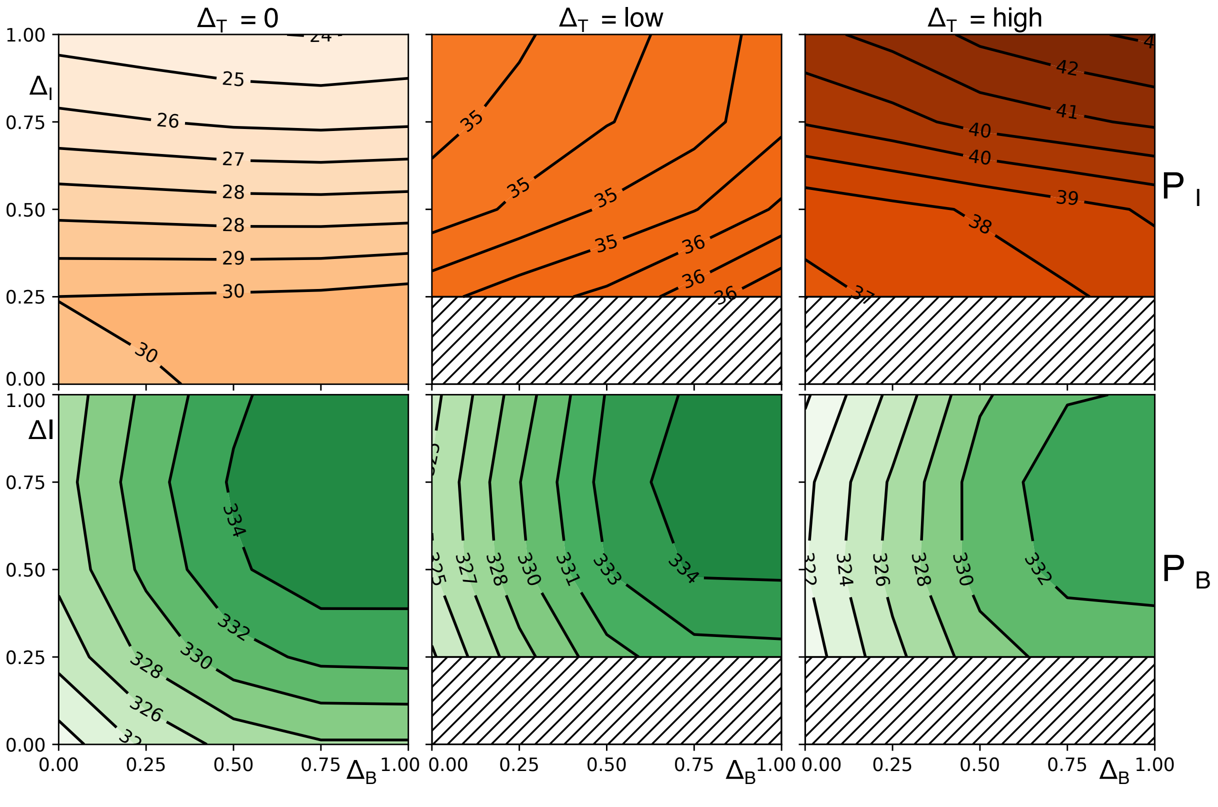


Figure S5.2: Partial dependence graphs of the total biomass production on the basal ($P_B$) and intermediate ($P_I$) level, in the same style as the figures in the main text. See the next section for an explanation for why $P_B$ is approximately constant.

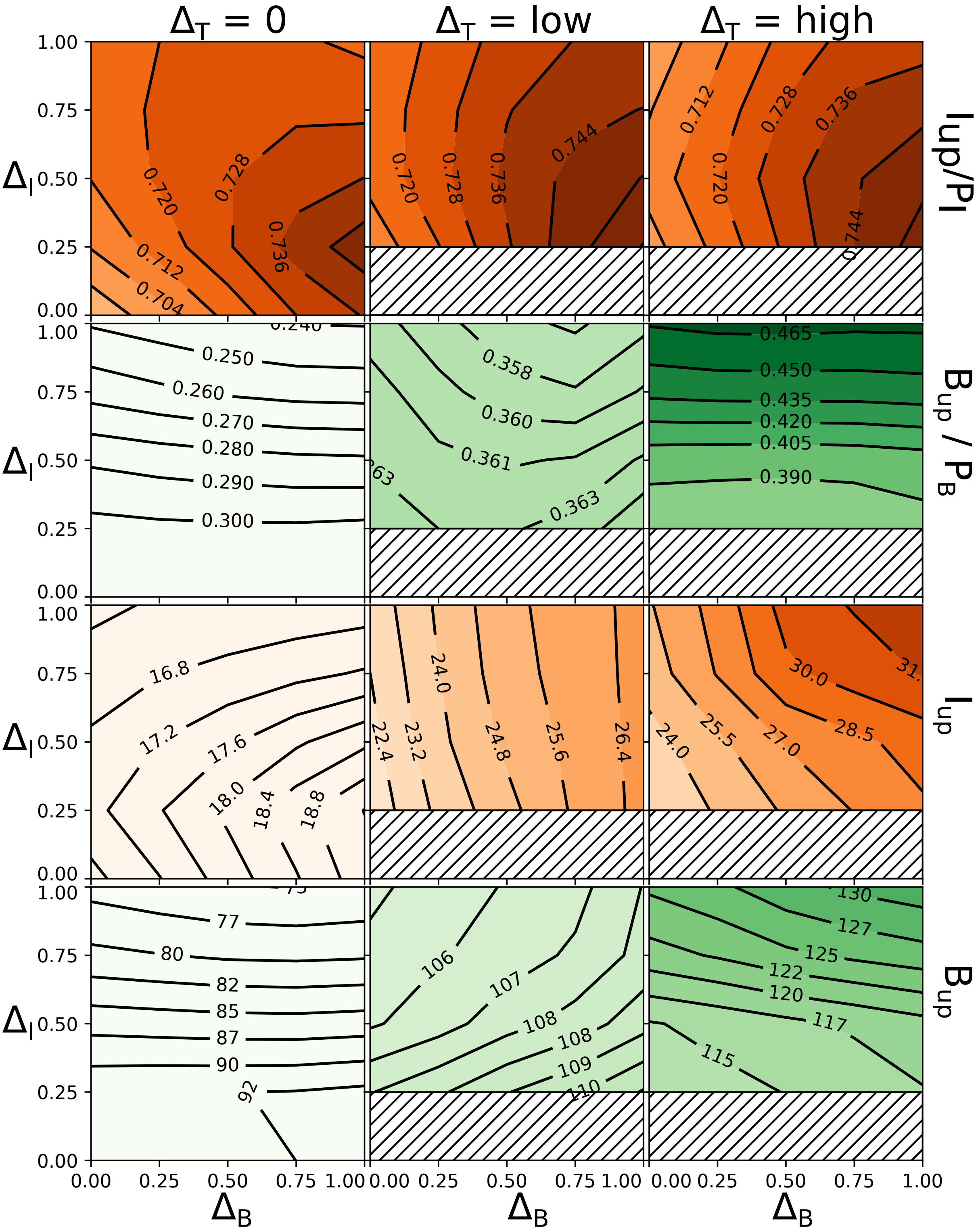


Figure S5.3: Partial dependence graphs of the total biomass flowing to the top and intermediate level ($I_{\text{up}}$ and $B_{\text{up}}$, respectively), and as a fraction of the total biomass production of the intermediate and top level ($I_{\text{up}}/P_I$ and $B_{\text{up}}/P_B$).

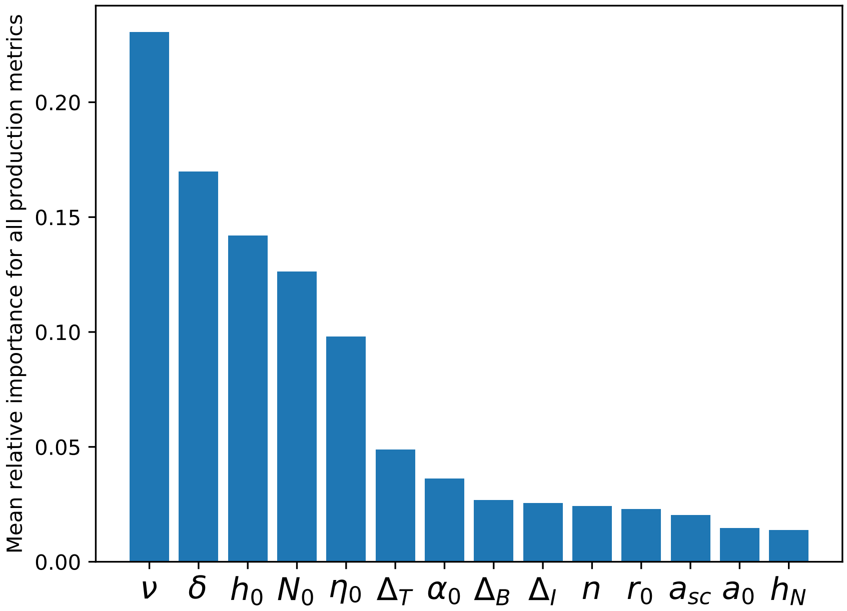


Figure S5.4: Mean relative importance of the different input parameters over *all* production metrics shown in the main text and this appendix: $P_B$, $P_I$, $P_T$, $B_{\mathrm{up}}$, $I_{\mathrm{up}}$, $B_{\mathrm{up}}/P_B$, $I_{\mathrm{up}}/P_I$, $(P/B)_B$, $(P/B)_I$, $(P/B)_T$, $P_T/P_B$. Like for the biomasses and CVs (Fig. 6, main text), parameters regulating the interaction between the top and intermediate level tend to be of higher importance than those of the intermediate-basal interaction. In particular, the trait difference between the top species $\Delta_T$ is of higher importance than that of $\Delta_I$ and $\Delta_B$.
