## Supplementary material for "Top predators govern multitrophic diversity effects in tritrophic food webs": S6

### S6 Link between mean free nutrient level and biomass production on the basal level

Here, we motivate why the biomass production of the basal level ($P_B$) is approximately constant in our model. Equation (1) (main text) reads:

$$\dot{N} = \delta(N_0 - N) - \frac{c_N}{c_C} \sum_i r_i B_i. \tag{S6.1}$$

Since $\sum_i r_i B_i = P_B$ we get:

$$\dot{N} = \delta(N_0 - N) - \frac{c_N}{c_C} P_B. \tag{S6.2}$$

When averaging over time, $\bar{\dot{N}} = 0$ must hold, hence:

$$\bar{P}_B = \frac{c_C}{c_N} \delta(N_0 - \bar{N}). \tag{S6.3}$$

This means that the time-averaged basal production $\bar{P}_B$ is fully determined by $\delta, N_0$, and $\bar{N}$. This relationship can clearly be observed when comparing the partial dependence plots of $N$ (Figure 3, main text) and $P_B$ (Figure S5.2 ). Moreover, the fact that $P_B$ barely varies is explained by the relatively low average values of $N$ (between approx. 45 and 190), as compared to the average $N_0$ of approximately 1200 (cf. Table 1).

This property is nicely captured by the random forest trained on the data for $P_B$ (cf. Figure S6.1).

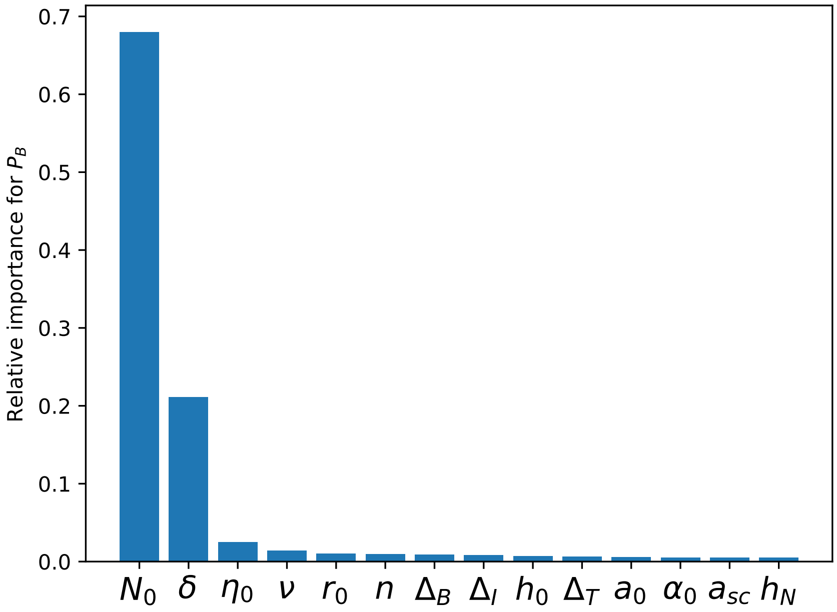


Figure S6.1: Relative importance of the different parameters for predicting the total biomass production on the basal level, $P_B$. Even though the random forest has no knowledge of the dynamical equations of our food webs, it accurately predicts that only $N_0$ and $\delta$ are of high importance to determine $P_B$.
