## Supplementary material for "Top predators govern multitrophic diversity effects in tritrophic food webs": S7

### S7 Relative biomasses per trophic level

Information on the contribution of individual species' biomasses to their respective trophic level helps with understanding the patterns of the biomass production efficiencies and food web efficiency (see Figure 5, main text). For example, a higher proportion of the fast growing species $B_1$ leads to an increased basal biomass production of the trophic level as a whole. In Figure S7.1, the proportion of $X_1$ is defined as $\frac{X_1}{X_1+X_2}$, with $X \in \{B, I, T\}$. To fairly asses the relative biomass of the fast growing species, two species have to be present on that trophic level. For that reason, all regions where $\Delta_X < 0.25$ have been excluded when calculating the relative biomass of $X_1$.

While the OOB-scores (measure for the goodness-of-fit of the random forest model, see Appendix S3.2) for $I_1$ and $T_1$ are good, they are negative for $B_1$ (cf. Table S7.1). This means that the random forest's prediction is worse than a constant prediction of the mean relative biomass $B_1$ every time. However, splitting the dataset by the amount of top diversity in the same way as for the partial dependence plots shows that the OOB score in the case of $\Delta_T = 0$ is still relatively accurate.

| Outcome variable | Overall OOB score | $\Delta_T = 0$ | $\Delta_T$ = low | $\Delta_T$ = high |
|---|---|---|---|---|
| rel. $T_1$ | 0.54 | n.a. | 0.53 | 0.46 |
| rel. $I_1$ | 0.88 | 0.74 | 0.84 | 0.88 |
| rel. $B_1$ | −0.05 | 0.52 | −0.39 | −0.19 |

Table S7.1: OOB scores estimating the accuracy of the random forest model, for the relative biomasses of $B_1, I_1$, and $T_1$. The second column shows the overall OOB score, i.e., of all the data, whereas the next three show the OOB scores separated by $\Delta_T$, as in Figure S7.1. An OOB score of 1 represents a perfect model prediction, whereas an OOB score of 0 means that the model is as accurate as simply predicting the mean outcome value every time. Therefore, we do not rely on the predictions for $B_1$ when $\Delta_T > 0$.

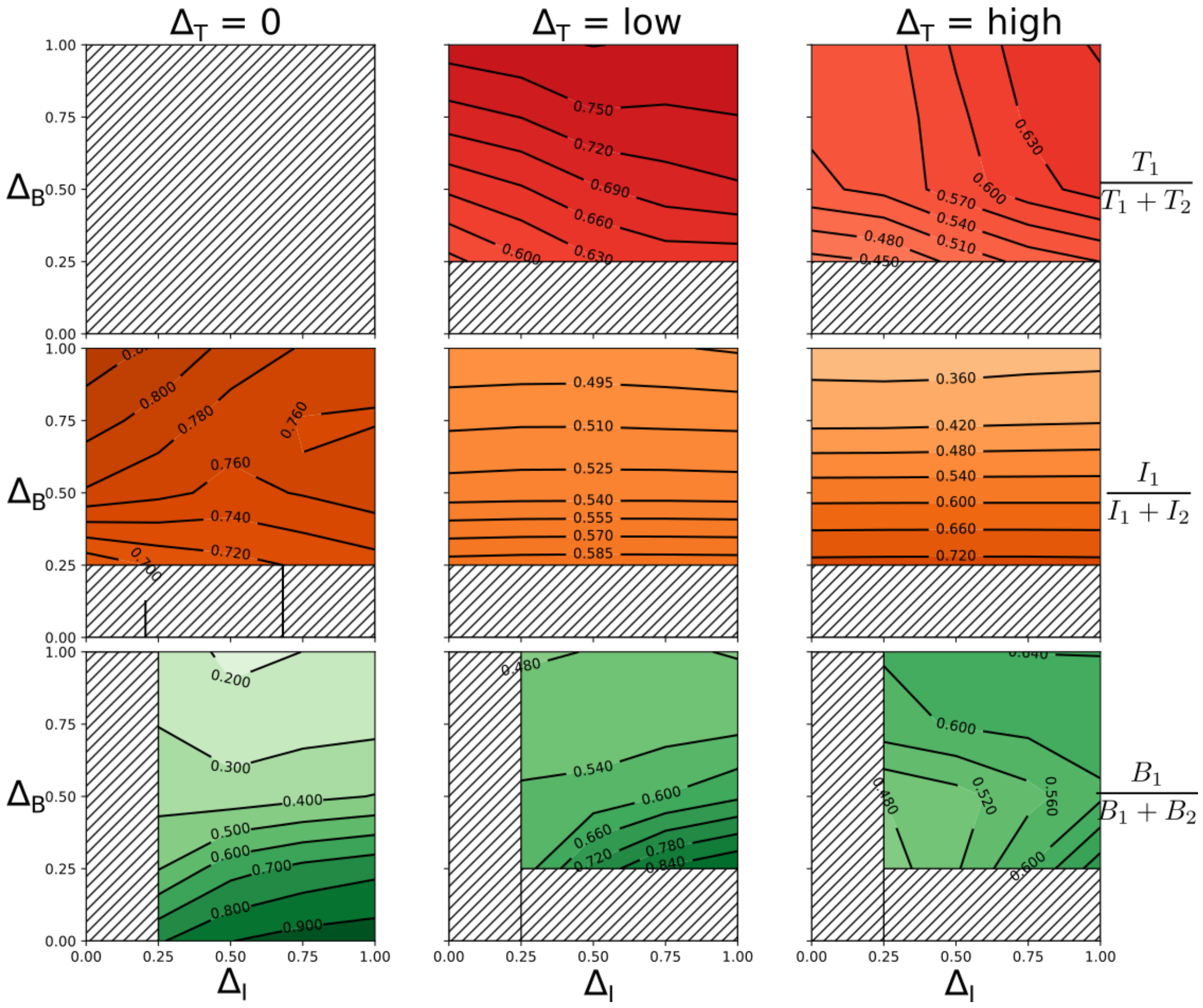

Figure S7.1: Partial dependence graphs of the trait differences $\Delta_B$ and $\Delta_I$, for $\Delta_T$ = 0, low $\Delta_T$, and high $\Delta_T$ = 1 (for more information see Methods), on the relative biomass of the fast growing $B_1$ (green), $I_1$ (orange), and $T_1$ (red), in the same style as Figures 3-5. See text for details on the additionally excluded areas.
