## Supplementary material for "Top predators govern multitrophic diversity effects in tritrophic food webs": Code and data: README.txt

*************** how to run our code ***************
It is important that the folder structure stays conserved. 
Producing and analyzing the data may only be possible on a UNIX system.
We can confirm it is possible to do everything on Ubuntu, Linux Mint, and macOS.

1) to produce the data 
	1.1 install Python 3, and the SUNDIALS suite (2.7.0) https://computing.llnl.gov/projects/sundials/sundials-software 
	1.2 Compile sys_solve.c (in folder /to_produce_data/c) into executable sys_solve
		* e.g. gcc -o sys_solve sys_solve.c  -lm -lgsl -lgslcblas -lsundials_cvode -lsundials_nvecserial
	1.3 Run analyze_web.py (in folder /to_produce_data/python) with desired parameters
		* e.g. to check 1024 parameter combinations, each for 50 initital conditions, for the BIT food web:
				webid = 7 
				total = 1024  
				duplicates = 50

		* this code can be run in parallel, so make sure to put the available amount of cores into num_cores
		* in our paper, we ran the python code with the following, for duplicates = 200:
				chain: total = 1024
				B, I, T: total = 4096
				BI, BT, IT: total = 16384
				BIT: total = 65536
		* to try different parameter intervals to sample from, simply change the corresponding variables
			e.g. to change to sampling Hill exponent n from [1, 1.5] set
				n_a = np.array([1, 1.5]) (line 59)
				
	1.4 C output files are stored in /to_produce_data/c_output_files/ (they are not needed for further analysis and can be removed if so desired)
	1.5 Python output files are stored in /to_produce_data/python_output_files
		These should be removed if starting a new simulation with the same webid, total, and duplicate variables, otherwise new simulation outcomes will be appended. 
	1.6 To analyze the Python output files, move them into /to_analyze_data/food_web_data


2) to analyze the data
	2.1 For this step, only Python (with various common packages such as NumPy, pandas, and scikit-learn) is needed when using the data files we provided. It may be possible to do this on a Windows system, but we cannot confirm this. 
	2.2 For the partial dependency plots PDPbox is required: https://github.com/SauceCat/PDPbox
		(can be installed in Anaconda through conda install -c conda-forge pdpbox)
	2.2 To generate Table 2, Figure 3, 4, 5, 6, or the various Appendix Figures, simply run the corresponding python file. 
		* for Figure S3.3 (in /to_produce_data/code/python) the C executable is required as well (see step 1.2)
		* when trying to plot data other than the data we provided make sure to update the file paths in /to_analyze_data/import_raw_data.py

*******************************************************
These files were created and tested to work with 
the following software versions (although particularly
for Python and its packages other versions are expected
to work as well): 
	* SUNDIALS 2.7.0
	* gcc 7.5.0 and 4.2.1
	* Python 3.7.6
	* NumPy 1.18.1
	* scikit-learn 0.22.1
	* matplotlib 3.1.3
