## Supplementary material for "Top predators govern multitrophic diversity effects in tritrophic food webs": Code and data: calc_pdps.py

```
import numpy as np
import pandas as pd

# calculate the median of all outcomes for the PDP
def calc_pdps(model, data_pd):
    med_vals = np.zeros((5, 5, 3))
    for tdiff in [0, 1, 2]: #[0, low, high]
        for bdiff in [0, 0.25, 0.5, 0.75, 1]:
            for idiff in [0, 0.25, 0.5, 0.75, 1]:
                if tdiff == 1:
                    tdiff1 = 0.25
                    p_data1 = data_pd.copy()
                    p_data1.loc[:, 'B_diff'] = bdiff
                    p_data1.loc[:, 'I_diff'] = idiff
                    p_data1.loc[:, 'T_diff'] = tdiff1
                    tdiff2 = 0.5
                    p_data2 = data_pd.copy()
                    p_data2.loc[:, 'B_diff'] = bdiff
                    p_data2.loc[:, 'I_diff'] = idiff
                    p_data2.loc[:, 'T_diff'] = tdiff2
                    drop_indices = []
                    for i in range(len(p_data1)):
                        if (p_data1.loc[i, 'T_diff'] > 0 
                            and p_data1.loc[i, 'I_diff'] == 0):
                            drop_indices.append(i)
                            
                    p_data1 = p_data1.drop(drop_indices)
                    p_data2 = p_data2.drop(drop_indices)
                    try:
                        med = np.median(model.predict(pd.concat([p_data1, p_data2])))
                    except ValueError:
                        med = 0
                    med_vals[int(4*bdiff), int(4*idiff), tdiff] = med
                elif tdiff == 2:
                    tdiff1 = 0.75
                    p_data1 = data_pd.copy()
                    p_data1.loc[:, 'B_diff'] = bdiff
                    p_data1.loc[:, 'I_diff'] = idiff
                    p_data1.loc[:, 'T_diff'] = tdiff1
                    tdiff2 = 1
                    p_data2 = data_pd.copy()
                    p_data2.loc[:, 'B_diff'] = bdiff
                    p_data2.loc[:, 'I_diff'] = idiff
                    p_data2.loc[:, 'T_diff'] = tdiff2
                    drop_indices = []
                    for i in range(len(p_data1)):
                        if (p_data1.loc[i, 'T_diff'] > 0 
                            and p_data1.loc[i, 'I_diff'] == 0):
                            drop_indices.append(i)
                            
                    p_data1 = p_data1.drop(drop_indices)
                    p_data2 = p_data2.drop(drop_indices)
                    try:
                        med = np.median(model.predict(pd.concat([p_data1, p_data2])))
                    except ValueError:
                        med = 0
                    med_vals[int(4*bdiff), int(4*idiff), tdiff] = med
                else:
                    p_data = data_pd.copy()
                    p_data.loc[:, 'B_diff'] = bdiff
                    p_data.loc[:, 'I_diff'] = idiff
                    p_data.loc[:, 'T_diff'] = tdiff
                    med = np.median(model.predict(p_data))
                    med_vals[int(4*bdiff), int(4*idiff), tdiff] = med
    return med_vals
```
