## Supplementary material for "Top predators govern multitrophic diversity effects in tritrophic food webs": Code and data: calc_prods.py

```
#!/usr/bin/env python3
# -*- coding: utf-8 -*-
"""
Created on Fri Jun 5 2020

@author: ruben
"""

import numpy as np

def calc_prods(unformatted_biomass, params, webid):
    
    Bdiff = 0
    Idiff = 0
    Tdiff = 0
    
    
    Biomass = np.zeros(7)
    Biomass[0] = unformatted_biomass[0] # N
    Biomass[1] = unformatted_biomass[1] # B1
    if webid == 0: # chain
        Biomass[3] = unformatted_biomass[2] # I1
        Biomass[5] = unformatted_biomass[3] # T1
    elif webid == 1: # B
        Biomass[2] = unformatted_biomass[2] # B2
        Biomass[3] = unformatted_biomass[3] # I1
        Biomass[5] = unformatted_biomass[4] # T1
        Bdiff = params['B_diff']
    
    elif webid == 2: # I
        Biomass[3] = unformatted_biomass[2] # I1
        Biomass[4] = unformatted_biomass[3] # I2
        Biomass[5] = unformatted_biomass[4] # T1
        Idiff = params['I_diff']
    
    elif webid == 3: # BI
        Biomass[2] = unformatted_biomass[2] # B2
        Biomass[3] = unformatted_biomass[3] # I1
        Biomass[4] = unformatted_biomass[4] # I2
        Biomass[5] = unformatted_biomass[5] # T1
        Bdiff = params['B_diff']
        Idiff = params['I_diff']
    
    elif webid == 4: # T
        Biomass[3] = unformatted_biomass[2] # I1
        Biomass[5] = unformatted_biomass[3] # T1
        Biomass[6] = unformatted_biomass[4] # T2
        Tdiff = params['T_diff']
    
    elif webid == 5: # BT
        Biomass[2] = unformatted_biomass[2] # B2
        Biomass[3] = unformatted_biomass[3] # I1
        Biomass[5] = unformatted_biomass[4] # T1
        Biomass[6] = unformatted_biomass[5] # T2
        Bdiff = params['B_diff']
        Tdiff = params['T_diff']
    
    elif webid == 6: # IT
        Biomass[3] = unformatted_biomass[2] # I1
        Biomass[4] = unformatted_biomass[3] # I2
        Biomass[5] = unformatted_biomass[4] # T1
        Biomass[6] = unformatted_biomass[5] # T2
        Idiff = params['I_diff']
        Tdiff = params['T_diff']
    
    elif webid == 7: # BIT
        Biomass = unformatted_biomass
        Bdiff = params['B_diff']
        Idiff = params['I_diff']
        Tdiff = params['T_diff']
        
    delta_ch = params['delta']
    
    N = np.array(Biomass[0])
    B = np.array(Biomass[1:3])
    I = np.array(Biomass[3:5])
    T = np.array(Biomass[5:7])
    
    r_inc = 0.5
    h_dec = 0.5
    eta_dec = 0.5
    
    e = 0.33
    eps = 0.33

   
    B_inc = 1 + Bdiff*r_inc
    I_inc = 1 + Idiff*h_dec
    T_inc = 1 + Tdiff*eta_dec
    
    delta = delta_ch*np.array([1, 
                               B_inc, 1/B_inc,  # 1 2
                               I_inc, 1/I_inc,  # 3 4
                               T_inc, 1/T_inc]) # 5 6

    Tprod = np.sum(delta[5:7]*T)
    Iup = Tprod/eps
    
    Iout = np.sum(delta[3:5]*I)
    Iprod = Iup + Iout
    Bup = Iprod/e
    
    Bout = np.sum(delta[1:3]*B)
    Bprod = Bup + Bout
    
    if Bprod > 0:
        PtopPbasal = Tprod/Bprod
    else:
        PtopPbasal = 0
        
    if np.sum(B) > 0:
        PBbasal = Bprod/np.sum(B)
    else:
        PBbasal = 0
    
    if np.sum(T) > 0:
        PBtop = Tprod/np.sum(T)
    else:
        PBtop = 0
   
    prods = np.array([PtopPbasal, Bup, PBbasal, Tprod])
    
    return prods

def all_prods(unformatted_biomass, params, webid):
    
    Bdiff = 0
    Idiff = 0
    Tdiff = 0
    
    
    Biomass = np.zeros(7)
    Biomass[0] = unformatted_biomass[0] # N
    Biomass[1] = unformatted_biomass[1] # B1
    if webid == 0: # chain
        Biomass[3] = unformatted_biomass[2] # I1
        Biomass[5] = unformatted_biomass[3] # T1
    elif webid == 1: # B
        Biomass[2] = unformatted_biomass[2] # B2
        Biomass[3] = unformatted_biomass[3] # I1
        Biomass[5] = unformatted_biomass[4] # T1
        Bdiff = params['B_diff']
    
    elif webid == 2: # I
        Biomass[3] = unformatted_biomass[2] # I1
        Biomass[4] = unformatted_biomass[3] # I2
        Biomass[5] = unformatted_biomass[4] # T1
        Idiff = params['I_diff']
    
    elif webid == 3: # BI
        Biomass[2] = unformatted_biomass[2] # B2
        Biomass[3] = unformatted_biomass[3] # I1
        Biomass[4] = unformatted_biomass[4] # I2
        Biomass[5] = unformatted_biomass[5] # T1
        Bdiff = params['B_diff']
        Idiff = params['I_diff']
    
    elif webid == 4: # T
        Biomass[3] = unformatted_biomass[2] # I1
        Biomass[5] = unformatted_biomass[3] # T1
        Biomass[6] = unformatted_biomass[4] # T2
        Tdiff = params['T_diff']
    
    elif webid == 5: # BT
        Biomass[2] = unformatted_biomass[2] # B2
        Biomass[3] = unformatted_biomass[3] # I1
        Biomass[5] = unformatted_biomass[4] # T1
        Biomass[6] = unformatted_biomass[5] # T2
        Bdiff = params['B_diff']
        Tdiff = params['T_diff']
    
    elif webid == 6: # IT
        Biomass[3] = unformatted_biomass[2] # I1
        Biomass[4] = unformatted_biomass[3] # I2
        Biomass[5] = unformatted_biomass[4] # T1
        Biomass[6] = unformatted_biomass[5] # T2
        Idiff = params['I_diff']
        Tdiff = params['T_diff']
    
    elif webid == 7: # BIT
        Biomass = unformatted_biomass
        Bdiff = params['B_diff']
        Idiff = params['I_diff']
        Tdiff = params['T_diff']
        
    delta_ch = params['delta']
    
    N = np.array(Biomass[0])
    B = np.array(Biomass[1:3])
    I = np.array(Biomass[3:5])
    T = np.array(Biomass[5:7])
    
    r_inc = 0.5
    h_dec = 0.5
    eta_dec = 0.5
    
    e = 0.33
    eps = 0.33

   
    B_inc = 1 + Bdiff*r_inc
    I_inc = 1 + Idiff*h_dec
    T_inc = 1 + Tdiff*eta_dec
    
    delta = delta_ch*np.array([1, 
                               B_inc, 1/B_inc,  # 1 2
                               I_inc, 1/I_inc,  # 3 4
                               T_inc, 1/T_inc]) # 5 6

    Tprod = np.sum(delta[5:7]*T)
    Iup = Tprod/eps
    
    Iout = np.sum(delta[3:5]*I)
    Iprod = Iup + Iout
    Bup = Iprod/e
    
    Bout = np.sum(delta[1:3]*B)
    Bprod = Bup + Bout
    
    if Bprod > 0:
        PtopPbasal = Tprod/Bprod
        Bup_prop = Bup/Bprod
    else:
        PtopPbasal = 0
        Bup_prop = 0
        
    if np.sum(B) > 0:
        PBbasal = Bprod/np.sum(B)
    else:
        PBbasal = 0
        
    if np.sum(I) > 0:
        PBI = Iprod/np.sum(I)
    else: 
        PBI = 0
    if Iprod > 0:
        Iup_prop = Iup/Iprod
    else: 
        Iup_prop = 0
        
    if np.sum(T) > 0:
        PBtop = Tprod/np.sum(T)
    else:
        PBtop = 0
        
    allprod = np.array([Bprod, Iprod, 
                       Bup, Iup, Bup_prop, Iup_prop, 
                       Tprod, PBbasal, PBI, PBtop, PtopPbasal])    
        
    return allprod
```
