## Supplementary material for "Top predators govern multitrophic diversity effects in tritrophic food webs": Code and data: fig5.py

```
#!/usr/bin/env python3
# -*- coding: utf-8 -*-
"""
Created on Fri Jun 5 2020

@author: ruben
"""

# this script will generate Figure 5 (main text)
import pandas as pd
import numpy as np
import ast
import matplotlib.pyplot as plt

import time

from sklearn.ensemble import ExtraTreesRegressor

from import_raw_data import import_data
from calc_prods import calc_prods
from calc_pdps import calc_pdps

# %%===========================================================================
# import data
# =============================================================================
p_d_list = [[], [], [], [], [], [], [], []]
ext = [[], [], [], [], [], [], [], []]
TLmeans = [[], [], [], [], [], [], [], []]
cvTLs = [[], [], [], [], [], [], [], []]
means = [[], [], [], [], [], [], [], []]

levelstr = ['PT PB', 'Bup', 'PB B', 'PT']

#range for contour lines
vmin_arr = [0.017, 77, 0.06, 5.4]
vmax_arr = [0.034, 130, 0.12, 10.5]

fmt_arr = ['%.4f', '%i', '%.3f', '%.1f'] # for formatting


# make sure the axes can be plotted correctly
contourBax = [0, 0.25, 0.5, 0.75, 1]
contourIax = [[0, 0.25, 0.5, 0.75, 1],[0.25, 0.5, 0.75, 1],
              [0.25, 0.5, 0.75, 1]]
I_startindex = [0, 1, 1]
# colormap and contour line colors             
cmapstr = ['Blues', 'Greens', 'Greens', 'Reds']

[p_d_list[0], p_d_list[1], p_d_list[2], p_d_list[3], p_d_list[4],
p_d_list[5], p_d_list[6], p_d_list[7], ext[0], ext[1], ext[2], ext[3],
ext[4], ext[5], ext[6], ext[7], TLmeans[0], TLmeans[1], TLmeans[2], 
TLmeans[3], TLmeans[4], TLmeans[5], TLmeans[6], TLmeans[7], cvTLs[0], 
cvTLs[1], cvTLs[2], cvTLs[3], cvTLs[4], cvTLs[5], cvTLs[6], cvTLs[7], 
means[0], means[1], means[2], means[3], means[4], means[5], means[6], 
means[7]] = import_data()

# put production metrics (figure 5) in same structure as the others
prods = []
for web in range(8):
    web_prods = []
    for combination in range(len(p_d_list[web])):
        prods_attr = []
        for attr in range(len(means[web][combination])):
            prod = calc_prods(means[web][combination][attr], 
                               p_d_list[web][combination], web)
            prods_attr.append(prod)
        web_prods.append(prods_attr)
    prods.append(web_prods)

# make sure that all trait differences are added to every data point          
def add_all_tdiffs_everywhere(web):
    if web == 0:
        add_dict = {'B_diff': 0, 'I_diff': 0, 'T_diff': 0}
    if web == 1:
        add_dict = {'I_diff': 0, 'T_diff': 0}
    if web == 2:
        add_dict = {'B_diff': 0, 'T_diff': 0}
    if web == 3:
        add_dict = {'T_diff': 0}
    if web == 4:
        add_dict = {'B_diff': 0, 'I_diff': 0}
    if web == 5:
        add_dict = {'I_diff': 0}
    if web == 6:
        add_dict = {'B_diff': 0}
    if web == 7:
        return
    for i in p_d_list[web]:
            i.update(add_dict)
    return

def check_tdiff(tdiff, value):
    if value == '0':
        if tdiff == 0:
            return 1
        else:
            return 0
    if value == 'low':
        if tdiff == 0.25:
            return 1
        elif tdiff == 0.5:
            return 1
        else:
            return 0
    if value == 'high':
        if tdiff == 0.75:
            return 1
        elif tdiff == 1:
            return 1
        else:
            return 0
    
feature_labels = ['B_diff', 'I_diff', 'T_diff', 'N0', 'h_N', 'r0', 'a0', 'h0', 
                'alpha0', 'eta0', 'delta', 'n', 'nu', 'a_scale']
for web in range(8):
    add_all_tdiffs_everywhere(web)
# %%===========================================================================
# "main"
# =============================================================================

num_cores = 4
n_trees = 2000
    
fig, axes = plt.subplots(4, 3, sharex = 'all', sharey = 'all',
                         figsize = (12, 12))
for level in [0, 1, 2, 3]:
    # first prepare data for random forest algorithm
    function = []
    data_samples = []
    for web in [0, 1, 2, 3, 6, 7]: # do not treat webs 4 and 5
        for i in range(len(p_d_list[web])):
            if 0 in ext[web][i]:
                tdiff = p_d_list[web][i]['T_diff']
                indices = np.where(ext[web][i] == 0)
                for j in indices[0]:
                    function.append(prods[web][i][j][level])
                    values = []
                    for key in feature_labels:
                        values.append(p_d_list[web][i][key])
                    data_samples.append(values)
    
    function = np.asarray(function)
    data_samples = np.asarray(data_samples)
    ts = time.time()
    clf = ExtraTreesRegressor(n_estimators = n_trees, bootstrap = 1, 
                              oob_score = 1, n_jobs = num_cores)
    model = clf.fit(data_samples, function)

    # generate 2d partial dependence plot
    data_pd = pd.DataFrame(data = data_samples, 
                              index = range(len(data_samples)),
                              columns = feature_labels)
    
    P = calc_pdps(model, data_pd)
    for tdiff in [0, 1, 2]:
        CSf = axes[3 - level, tdiff].contourf(contourBax, contourIax[tdiff], 
                           P[:, I_startindex[tdiff]:, tdiff].T, 
                           cmap = cmapstr[level],
                           vmin = vmin_arr[level], vmax = vmax_arr[level])

        CS = axes[3 - level, tdiff].contour(contourBax, contourIax[tdiff], 
                         P[:, I_startindex[tdiff]:, tdiff].T, 
                         colors = 'k',
                         vmin = vmin_arr[level], vmax = vmax_arr[level])


        axes[3 - level, tdiff].set_aspect('equal')
        axes[3 - level, tdiff].clabel(CS, inline = 1, fontsize = 10, fmt = fmt_arr[level])
        
axes[0, 0].set_yticks(contourBax)
axes[0, 0].set_xticks(contourBax)
axes[0, 0].set_ylim([0, 1])
for i in [0, 1, 2, 3]:
    for j in [1, 2]:
        axes[i, j].fill_between([0, 1], 0.25, hatch = 3*'/', edgecolor = 'k', facecolor = 'none')
plt.tight_layout()
fig.subplots_adjust(wspace=-0.52, hspace=0.03)
plt.savefig('fig_5.pdf')
```
