## Supplementary material for "Top predators govern multitrophic diversity effects in tritrophic food webs": Code and data: fig6.py

```
#!/usr/bin/env python3
# -*- coding: utf-8 -*-
"""
Created on Fri Jun 5 2020

@author: ruben
"""

# this script will generate Figure 6 (main text)

import pandas as pd
import numpy as np
import ast
import matplotlib.pyplot as plt

import time

from sklearn.ensemble import ExtraTreesRegressor

from import_raw_data import import_data
 # %%===========================================================================
# import data
# =============================================================================
p_d_list = [[], [], [], [], [], [], [], []]
ext = [[], [], [], [], [], [], [], []]
TLmeans = [[], [], [], [], [], [], [], []]
cvTLs = [[], [], [], [], [], [], [], []]
means = [[], [], [], [], [], [], [], []]

# levelstr = ['PB','PI','PT','B_up','I_up', 'Bupprop', 'Iupprop', 'PBbasal',
#             'PBI', 'PBtop', 'PtopPbasal']
levelstr = ['N', 'B', 'I', 'T']

[p_d_list[0], p_d_list[1], p_d_list[2], p_d_list[3], p_d_list[4],
p_d_list[5], p_d_list[6], p_d_list[7], ext[0], ext[1], ext[2], ext[3],
ext[4], ext[5], ext[6], ext[7], TLmeans[0], TLmeans[1], TLmeans[2], 
TLmeans[3], TLmeans[4], TLmeans[5], TLmeans[6], TLmeans[7], cvTLs[0], 
cvTLs[1], cvTLs[2], cvTLs[3], cvTLs[4], cvTLs[5], cvTLs[6], cvTLs[7], 
means[0], means[1], means[2], means[3], means[4], means[5], means[6], 
means[7]] = import_data()

# make sure that all trait differences are added to every data point
def add_all_tdiffs_everywhere(web):
    if web == 0:
        add_dict = {'B_diff': 0, 'I_diff': 0, 'T_diff': 0}
    if web == 1:
        add_dict = {'I_diff': 0, 'T_diff': 0}
    if web == 2:
        add_dict = {'B_diff': 0, 'T_diff': 0}
    if web == 3:
        add_dict = {'T_diff': 0}
    if web == 4:
        add_dict = {'B_diff': 0, 'I_diff': 0}
    if web == 5:
        add_dict = {'I_diff': 0}
    if web == 6:
        add_dict = {'B_diff': 0}
    if web == 7:
        return
    for i in p_d_list[web]:
            i.update(add_dict)
    return

feature_labels = np.array(['B_diff', 'I_diff', 'T_diff', 'N0', 'h_N', 'r0', 'a0', 'h0', 
                'alpha0', 'eta0', 'delta', 'n', 'nu', 'a_scale'])
for web in range(8):
    add_all_tdiffs_everywhere(web)
    
    
# %%===========================================================================
# "main"
# =============================================================================

num_cores = 4
n_trees = 2000


params = np.array(['nu', 'N0', 'eta0', 'delta', 'h0', 'T_diff', 'I_diff', 'n', 
          'alpha0', 'r0', 'B_diff', 'a_scale', 'a0', 'h_N'])

labels_dict = {'nu': r'$\nu$',
               'N0': r'$N_0$',
               'eta0': r'$\eta_0$',
               'delta': r'$\delta$',
               'h0': r'$h_0$', 
               'T_diff': r'$\Delta_T$',
               'I_diff': r'$\Delta_I$', 
               'n': r'$n$', 
               'alpha0': r'$\alpha_0$',
               'r0': r'$r_0$', 
               'B_diff': r'$\Delta_B$',
               'a_scale': r'$a_{sc}$',
               'a0': r'$a_0$',
               'h_N': r'$h_N$'}

# biomasses
imp_dict_B = {}
label_dict_B = {}


for level in range(4):
    # first prepare data for random forest algorithm
    function = []
    data_samples = []
    for web in [0, 1, 2, 3, 6, 7]: # do not treat webs 4 and 5
        for i in range(len(p_d_list[web])):
            if 0 in ext[web][i]:
                indices = np.where(ext[web][i] == 0)
                for j in indices[0]:
                    function.append(TLmeans[web][i][j][level])
                    values = []
                    for key in feature_labels:
                        values.append(p_d_list[web][i][key])
                    data_samples.append(values)
                        
    function = np.asarray(function)
    data_samples = np.asarray(data_samples)
    ts = time.time()
    clf = ExtraTreesRegressor(n_estimators = n_trees, bootstrap = 1, oob_score = 1, n_jobs = num_cores)
    model = clf.fit(data_samples, function)
    importances = model.feature_importances_
    indices = np.argsort(importances)[::-1]
            
    labels = [feature_labels[i] for i in indices]
    
    imp_dict_B[levelstr[level]] = importances[indices]
    label_dict_B[levelstr[level]] = labels

# make plot

#calculate mean importance per parameter
mean_imps_B= []

for param in labels_dict:
    mean_val = 0
    for metric in levelstr:
        param_index = label_dict_B[metric].index(param)
        mean_val += imp_dict_B[metric][param_index]
    mean_imps_B.append(mean_val/4)
mean_imps_B = np.asarray(mean_imps_B)  
order = np.argsort(-mean_imps_B)

width = 0.2
colors = ['C0', 'C2', 'C1', 'C3']
xlabels = [labels_dict[param] for param in params[order]]

fig, ax = plt.subplots(2, 1, figsize = (7, 10))

for i in range(len(feature_labels)):
    paramstr = params[order][i]
    for metric in levelstr:
        metric_num = levelstr.index(metric)
        ax[0].bar(i - 3*width/2 + width*metric_num, 
                imp_dict_B[metric][label_dict_B[metric].index(params[order][i])],
                width = width, color = colors[metric_num],
                label = levelstr[metric_num] if paramstr == 'nu' else '')
x = np.arange(len(labels_dict)) # the label locations
ax[0].set_ylabel('Mean relative importance for biomasses')
ax[0].set_xticks(x)
ax[0].set_xticklabels(xlabels, fontsize = 15)
ax[0].legend()

# CVs
imp_dict_CV = {}
label_dict_CV = {}


for level in range(4):
    # first prepare data for random forest algorithm
    function = []
    data_samples = []
    for web in [0, 1, 2, 3, 6, 7]: # do not treat webs 4 and 5
        for i in range(len(p_d_list[web])):
            if 0 in ext[web][i]:
                indices = np.where(ext[web][i] == 0)
                for j in indices[0]:
                    function.append(cvTLs[web][i][j][level])
                    values = []
                    for key in feature_labels:
                        values.append(p_d_list[web][i][key])
                    data_samples.append(values)
                        
    function = np.asarray(function)
    data_samples = np.asarray(data_samples)
    ts = time.time()
    clf = ExtraTreesRegressor(n_estimators = n_trees, bootstrap = 1, oob_score = 1, n_jobs = num_cores)
    model = clf.fit(data_samples, function)
    importances = model.feature_importances_
    indices = np.argsort(importances)[::-1]
            
    labels = [feature_labels[i] for i in indices]
    
    imp_dict_CV[levelstr[level]] = importances[indices]
    label_dict_CV[levelstr[level]] = labels

# make plot

#calculate mean importance per parameter
mean_imps_CV = []

for param in labels_dict:
    mean_val = 0
    for metric in levelstr:
        param_index = label_dict_CV[metric].index(param)
        mean_val += imp_dict_CV[metric][param_index]
    mean_imps_CV.append(mean_val/4)
mean_imps_CV = np.asarray(mean_imps_CV)  
order = np.argsort(-mean_imps_CV)

width = 0.2
colors = ['C0', 'C2', 'C1', 'C3']
xlabels = [labels_dict[param] for param in params[order]]

for i in range(len(feature_labels)):
    paramstr = params[order][i]
    for metric in levelstr:
        metric_num = levelstr.index(metric)
        ax[1].bar(i - 3*width/2 + width*metric_num, 
                imp_dict_CV[metric][label_dict_CV[metric].index(params[order][i])],
                width = width, color = colors[metric_num],
                label = levelstr[metric_num] if paramstr == 'nu' else '')
x = np.arange(len(labels_dict)) # the label locations
ax[1].set_ylabel('Mean relative importance for CVs')
ax[1].set_xticks(x)
ax[1].set_xticklabels(xlabels, fontsize = 15)
ax[1].legend()

plt.savefig('fig_6.pdf')
```
