## Supplementary material for "Top predators govern multitrophic diversity effects in tritrophic food webs": Code and data: figS3-5.py

```
#!/usr/bin/env python3
# -*- coding: utf-8 -*-
"""
Created on Fri Oct  2 15:54:05 2020

@author: ruben
"""
# this script will generate Figure S3.5 (Appendix S3)

import pandas as pd
import numpy as np
import matplotlib.pyplot as plt

import time

from sklearn.ensemble import ExtraTreesRegressor

from import_raw_data import import_data

# %%===========================================================================
# import data
# =============================================================================
p_d_list = [[], [], [], [], [], [], [], []]
ext = [[], [], [], [], [], [], [], []]
TLmeans = [[], [], [], [], [], [], [], []]
cvTLs = [[], [], [], [], [], [], [], []]
means = [[], [], [], [], [], [], [], []]

[p_d_list[0], p_d_list[1], p_d_list[2], p_d_list[3], p_d_list[4],
p_d_list[5], p_d_list[6], p_d_list[7], ext[0], ext[1], ext[2], ext[3],
ext[4], ext[5], ext[6], ext[7], TLmeans[0], TLmeans[1], TLmeans[2], 
TLmeans[3], TLmeans[4], TLmeans[5], TLmeans[6], TLmeans[7], cvTLs[0], 
cvTLs[1], cvTLs[2], cvTLs[3], cvTLs[4], cvTLs[5], cvTLs[6], cvTLs[7], 
means[0], means[1], means[2], means[3], means[4], means[5], means[6], 
means[7]] = import_data()

# make sure that all trait differences are added to every data point          
def add_all_tdiffs_everywhere(web):
    if web == 0:
        add_dict = {'B_diff': 0, 'I_diff': 0, 'T_diff': 0}
    if web == 1:
        add_dict = {'I_diff': 0, 'T_diff': 0}
    if web == 2:
        add_dict = {'B_diff': 0, 'T_diff': 0}
    if web == 3:
        add_dict = {'T_diff': 0}
    if web == 4:
        add_dict = {'B_diff': 0, 'I_diff': 0}
    if web == 5:
        add_dict = {'I_diff': 0}
    if web == 6:
        add_dict = {'B_diff': 0}
    if web == 7:
        return
    for i in p_d_list[web]:
            i.update(add_dict)
    return

def check_tdiff(tdiff, value):
    if value == '0':
        if tdiff == 0:
            return 1
        else:
            return 0
    if value == 'low':
        if tdiff == 0.25:
            return 1
        elif tdiff == 0.5:
            return 1
        else:
            return 0
    if value == 'high':
        if tdiff == 0.75:
            return 1
        elif tdiff == 1:
            return 1
        else:
            return 0
    
feature_labels = ['B_diff', 'I_diff', 'T_diff', 'N0', 'h_N', 'r0', 'a0', 'h0', 
                'alpha0', 'eta0', 'delta', 'n', 'nu', 'a_scale']

for web in range(8):
    add_all_tdiffs_everywhere(web)
    
def calc_pdp_1d(data_pd):
    mean_vals = []
    err_vals = []
    for tdiff in [0, 0.25, 0.5, 0.75, 1]:
        p_data = data_pd.copy()
        p_data.loc[:, 'T_diff'] = tdiff
        preds = model.predict(p_data)
        mean_vals.append(np.mean(preds))
        err_vals.append(3*np.std(preds)/(len(p_data)**(1/2)))
    return np.asarray(mean_vals), np.asarray(err_vals)

# %%===========================================================================
# "main"
# =============================================================================

num_cores = 4
n_trees = 2000
    
function = []
data_samples = []
for web in [0, 1, 2, 3, 6, 7]: # do not treat webs 4 and 5
    for i in range(len(p_d_list[web])):
        if 0 in ext[web][i]:
            tdiff = p_d_list[web][i]['T_diff']
            indices = np.where(ext[web][i] == 0)
            for j in indices[0]:
                function.append(TLmeans[web][i][j][3])
                values = []
                for key in feature_labels:
                    values.append(p_d_list[web][i][key])
                data_samples.append(values)

function = np.asarray(function)
data_samples = np.asarray(data_samples)
ts = time.time()
clf = ExtraTreesRegressor(n_estimators = n_trees, bootstrap = 1, oob_score = 1, n_jobs = num_cores)
model = clf.fit(data_samples, function)

data_pd = pd.DataFrame(data = data_samples, 
                          index = range(len(data_samples)),
                          columns = feature_labels)

mean_pred, err_pred = calc_pdp_1d(data_pd)

mean_data = []
err_data = []

for tdiff in [0, 0.25, 0.5, 0.75, 1]:
    cur_data = function[data_pd['T_diff'] == tdiff]
    mean_data.append(np.mean(cur_data))
    err_data.append(3*np.std(cur_data)/len(cur_data)**(1/2))
    

fig, ax = plt.subplots(figsize = (8, 5))
tdiff = np.asarray([0, 0.25, 0.5, 0.75, 1])
ax.errorbar(tdiff+0.005, mean_data, yerr = err_data, linewidth = 2, color = 'gray',
            marker = 'o', label = 'data')
ax.errorbar(tdiff-0.005, mean_pred, yerr = err_pred, linewidth = 2, color = 'k',
            marker = 'o', label = 'RF prediction')
ax.set_xticks(tdiff)
ax.legend(fontsize = 14)
ax.tick_params(axis='both', which='major', labelsize=14)
ax.set_xlabel(r'$\Delta_T$', fontsize = 16)
ax.set_ylabel(r'Top level biomass', fontsize = 16)

plt.tight_layout
plt.savefig('S3-5.pdf')
```
