## Supplementary material for "Top predators govern multitrophic diversity effects in tritrophic food webs": Code and data: figS7-1.py

```
#!/usr/bin/env python3
# -*- coding: utf-8 -*-
"""
Created on Fri Jun 5 2020

@author: ruben
"""
# this script will generate Appendix S7 Table S7.1 and  Figure S7.1

import pandas as pd
import numpy as np
import matplotlib.pyplot as plt

import time

from sklearn.ensemble import ExtraTreesRegressor

from import_raw_data import import_data
from calc_prods import calc_prods

# %%===========================================================================
# import data
# =============================================================================
p_d_list = [[], [], [], [], [], [], [], []]
ext = [[], [], [], [], [], [], [], []]
TLmeans = [[], [], [], [], [], [], [], []]
cvTLs = [[], [], [], [], [], [], [], []]
means = [[], [], [], [], [], [], [], []]

levelstr = ['N', 'B', 'I', 'T']

#range for contour lines
vmin_arr = [0, 0, 0, 0]
vmax_arr = [1, 1, 1, 1]

# make sure the axes can be plotted correctly

contourBax = [0, 0.25, 0.5, 0.75, 1]
contourIax = [[0, 0.25, 0.5, 0.75, 1],[0.25, 0.5, 0.75, 1],
              [0.25, 0.5, 0.75, 1]]
I_startindex = [0, 1, 1]
tdiff_arr = ['0', 'low', 'high']
cmapstr = ['Blues', 'Greens', 'Oranges', 'Reds']


[p_d_list[0], p_d_list[1], p_d_list[2], p_d_list[3], p_d_list[4],
p_d_list[5], p_d_list[6], p_d_list[7], ext[0], ext[1], ext[2], ext[3],
ext[4], ext[5], ext[6], ext[7], TLmeans[0], TLmeans[1], TLmeans[2], 
TLmeans[3], TLmeans[4], TLmeans[5], TLmeans[6], TLmeans[7], cvTLs[0], 
cvTLs[1], cvTLs[2], cvTLs[3], cvTLs[4], cvTLs[5], cvTLs[6], cvTLs[7], 
means[0], means[1], means[2], means[3], means[4], means[5], means[6], 
means[7]] = import_data()

# make sure that all trait differences are added to every data point                      
def add_all_tdiffs_everywhere(web):
    if web == 0:
        add_dict = {'B_diff': 0, 'I_diff': 0, 'T_diff': 0}
    if web == 1:
        add_dict = {'I_diff': 0, 'T_diff': 0}
    if web == 2:
        add_dict = {'B_diff': 0, 'T_diff': 0}
    if web == 3:
        add_dict = {'T_diff': 0}
    if web == 4:
        add_dict = {'B_diff': 0, 'I_diff': 0}
    if web == 5:
        add_dict = {'I_diff': 0}
    if web == 6:
        add_dict = {'B_diff': 0}
    if web == 7:
        return
    for i in p_d_list[web]:
            i.update(add_dict)
    return

def check_tdiff(tdiff, value):
    if value == '0':
        if tdiff == 0:
            return 1
        else:
            return 0
    if value == 'low':
        if tdiff == 0.25:
            return 1
        elif tdiff == 0.5:
            return 1
        else:
            return 0
    if value == 'high':
        if tdiff == 0.75:
            return 1
        elif tdiff == 1:
            return 1
        else:
            return 0

def get_relative_biomass(means, web, level):
    if level == 0:
        pass
    
    if web == 0:
        return means[level]
    if web == 1:
        if level == 1:
            return means[1]/(means[1] + means[2])
        else: 
            return means[level + 1]
    if web == 2:
        if level == 1:
            return means[1]
        if level == 2:
            return means[2]/(means[2] + means[3])
        if level == 3:
            return means[4]
    if web == 3:
        if level == 1:
            return means[1]/(means[1] + means[2])
        if level == 2:
            return means[3]/(means[3] + means[4])
        if level == 3:
            return means[5]
    if web == 4:
        if level < 3:
            return means[level]
        if level == 3:
            return means[3]/(means[3] + means[4])
    if web == 5:
        if level == 1:
            return means[1]/(means[1] + means[2])
        if level == 2:
            return means[3]
        if level == 3:
            return means[4]/(means[4] + means[5])
    if web == 6:
        if level == 1:
            return means[1]
        if level == 2:
            return means[2]/(means[2] + means[3])
        if level == 3:
            return means[4]/(means[4] + means[5])
    if web == 7:
        return means[2*level - 1]/(means[2*level - 1] + means[2*level])
        
    
feature_labels = ['B_diff', 'I_diff', 'T_diff', 'N0', 'h_N', 'r0', 'a0', 'h0', 
                'alpha0', 'eta0', 'delta', 'n', 'nu', 'a_scale']
for web in range(8):
    add_all_tdiffs_everywhere(web)
    
def calc_pdps_2d(data_pd):
    med_vals = np.zeros((5, 5))
    for bdiff in [0, 0.25, 0.5, 0.75, 1]:
        for idiff in [0, 0.25, 0.5, 0.75, 1]:
                p_data = data_pd.copy()
                p_data.loc[:, 'B_diff'] = bdiff
                p_data.loc[:, 'I_diff'] = idiff
                drop_indices = []
                for i in range(len(p_data)):
                        if (p_data.loc[i, 'T_diff'] > 0 
                            and p_data.loc[i, 'I_diff'] == 0):
                            drop_indices.append(i)
                p_data = p_data.drop(drop_indices)
                try:
                    med = np.median(model.predict(p_data))
                except ValueError:
                    med = 0
                med_vals[int(4*bdiff), int(4*idiff)] = med
    return med_vals

# %%===========================================================================
# "main"
# =============================================================================
n_trees = 2000
num_cores = 4
# FIGURE S7.1
fig, axes = plt.subplots(3, 3, sharex = 'all', sharey = 'all',
                          figsize = (12, 12))
col = 0
for i_tdiff in [0, 1, 2]:
    for level in [1, 2, 3]:
        if (tdiff_arr[i_tdiff] == '0' and level == 3):
            pass # not valid
        else:
            # first prepare data for random forest algorithm
            function = []
            data_samples = []
            for web in [0, 1, 2, 3, 6, 7]: # do not treat webs 4 and 5
                for i in range(len(p_d_list[web])):
                    include_p = 0
                    
                    if level == 1:
                        if p_d_list[web][i]['B_diff'] >= 0.25:
                            include_p = 1
                    if level == 2:
                        if p_d_list[web][i]['I_diff'] >= 0.25:
                            include_p = 1
                    if level == 3:
                        if p_d_list[web][i]['T_diff'] >= 0.25:
                            include_p = 1
                            
                    if include_p:
                        if 0 in ext[web][i]:
                            tdiff = p_d_list[web][i]['T_diff']
                            include = check_tdiff(tdiff, tdiff_arr[i_tdiff])
                            if include:
                                indices = np.where(ext[web][i] == 0)
                                for j in indices[0]:
                                    rel_B = get_relative_biomass(means[web][i][j],
                                                                    web, level)
                                    function.append(rel_B)
                                    values = []
                                    for key in feature_labels:
                                        values.append(p_d_list[web][i][key])
                                    data_samples.append(values)
            
            function = np.asarray(function)
            data_samples = np.asarray(data_samples)
            ts = time.time()
            clf = ExtraTreesRegressor(n_estimators = n_trees, bootstrap = 1, oob_score = 1, n_jobs = num_cores)
            model = clf.fit(data_samples, function)

            #for Table S7.1
            print("OOB " + levelstr[level] + tdiff_arr[i_tdiff] + ': ' + str(model.oob_score_))

            # generate 2d partial dependence plot
            data_pd = pd.DataFrame(data = data_samples, 
                                      index = range(len(data_samples)),
                                      columns = feature_labels)
            data = calc_pdps_2d(data_pd)
           
            if (i_tdiff == 0 and level == 3):
                axes[3 - level, i_tdiff].set_aspect('equal')
            elif (level == 1):
                CSf = axes[3 - level, i_tdiff].contourf([0.25, 0.5, 0.75, 1],
                                                  contourIax[i_tdiff], 
                                                  data[1:, I_startindex[i_tdiff]:].T, 
                                                  cmap = cmapstr[level],
                                                  vmin = vmin_arr[level], 
                                                  vmax = vmax_arr[level])
    
                CS = axes[3 - level, i_tdiff].contour([0.25, 0.5, 0.75, 1],
                                                contourIax[i_tdiff], 
                             data[1:, I_startindex[i_tdiff]:].T, 
                             colors = 'k',
                             vmin = vmin_arr[level], vmax = vmax_arr[level])
                axes[3 - level, i_tdiff].set_aspect('equal')
                axes[3 - level, i_tdiff].clabel(CS, inline = 1, fontsize = 10, 
                                      fmt = '%.3f')
            else:            
                CSf = axes[3 - level, i_tdiff].contourf(contourBax, contourIax[i_tdiff], 
                               data[:, I_startindex[i_tdiff]:].T, 
                               cmap = cmapstr[level],
                               vmin = vmin_arr[level], vmax = vmax_arr[level])
    
                CS = axes[3 - level, i_tdiff].contour(contourBax, contourIax[i_tdiff], 
                             data[:, I_startindex[i_tdiff]:].T, 
                             colors = 'k',
                             vmin = vmin_arr[level], vmax = vmax_arr[level])
            
                axes[3 - level, i_tdiff].set_aspect('equal')
                axes[3 - level, i_tdiff].clabel(CS, inline = 1, fontsize = 10, 
                                      fmt = '%.3f')
            
axes[0, 0].set_yticks([0, 0.25, 0.5, 0.75, 1])
axes[0, 0].set_xticks([0, 0.25, 0.5, 0.75, 1])
axes[0, 0].set_ylim([0, 1])
for i in [0, 1, 2]:
    for j in [1, 2]:
        axes[i, j].fill_between([0, 1], 0.25, hatch = 3*'/', edgecolor = 'k', facecolor = 'none')
        if i == 2:
            axes[i, j].fill_betweenx([0, 1], 0.25, hatch = 3*'/', edgecolor = 'k', facecolor = 'white')
axes[1, 0].fill_between([0, 1], 0.25, hatch = 3*'/', edgecolor = 'k', facecolor = 'white')
axes[2, 0].fill_betweenx([0, 1], 0.25, hatch = 3*'/', edgecolor = 'k', facecolor = 'white')
axes[0, 0].fill_between([0, 1], 1, hatch = 3*'/', edgecolor = 'k', facecolor = 'white')
plt.tight_layout()
fig.subplots_adjust(wspace=-0.2, hspace=0.05)
plt.savefig('fig_S7-1.pdf')
```
