## Supplementary material for "Top predators govern multitrophic diversity effects in tritrophic food webs": Code and data: figsS5_and_S6.py

```
#!/usr/bin/env python3
# -*- coding: utf-8 -*-
"""
Created on Fri Jun 5 2020

@author: ruben
"""

# this script will generate Appendix S5 Figures S5.2, S5.3, S5.4 and S6 Figure S6.1

import pandas as pd
import numpy as np
import matplotlib.pyplot as plt

import time

from sklearn.ensemble import ExtraTreesRegressor

from import_raw_data import import_data
from calc_prods import all_prods
from calc_pdps import calc_pdps


num_cores = 4
n_trees = 2000

# %%===========================================================================
# import data
# =============================================================================
p_d_list = [[], [], [], [], [], [], [], []]
ext = [[], [], [], [], [], [], [], []]
TLmeans = [[], [], [], [], [], [], [], []]
cvTLs = [[], [], [], [], [], [], [], []]
means = [[], [], [], [], [], [], [], []]

levelstr_1 = ['PB', 'PI']
levelstr_2 = ['Bup', 'Iup', 'Bup PB', 'Iup PI']
levelstr_3 = ['PB', 'PI', 'Bup', 'Iup', 'Bup PB', 'Iup PI',
                'PT', 'PBbasal', 'PBI', 'PBtop', 'PtopPbasal']

[p_d_list[0], p_d_list[1], p_d_list[2], p_d_list[3], p_d_list[4],
p_d_list[5], p_d_list[6], p_d_list[7], ext[0], ext[1], ext[2], ext[3],
ext[4], ext[5], ext[6], ext[7], TLmeans[0], TLmeans[1], TLmeans[2], 
TLmeans[3], TLmeans[4], TLmeans[5], TLmeans[6], TLmeans[7], cvTLs[0], 
cvTLs[1], cvTLs[2], cvTLs[3], cvTLs[4], cvTLs[5], cvTLs[6], cvTLs[7], 
means[0], means[1], means[2], means[3], means[4], means[5], means[6], 
means[7]] = import_data()

# put production metrics in same structure as the others
prods_1 = []
prods_2 = []
prods_3 = []
for web in range(8):
    web_prods_1 = []
    web_prods_2 = []
    web_prods_3 = []
    for combination in range(len(p_d_list[web])):
        prods_1_attr = []
        prods_2_attr = []
        prods_3_attr = []
        for attr in range(len(means[web][combination])):
            prod_1 = all_prods(means[web][combination][attr], 
                               p_d_list[web][combination], web)[0:2]
            prod_2 = all_prods(means[web][combination][attr], 
                               p_d_list[web][combination], web)[2:6]
            prod_3 = all_prods(means[web][combination][attr], 
                               p_d_list[web][combination], web)
            prods_1_attr.append(prod_1)
            prods_2_attr.append(prod_2)
            prods_3_attr.append(prod_3)
        web_prods_1.append(prods_1_attr)
        web_prods_2.append(prods_2_attr)
        web_prods_3.append(prods_3_attr)
    prods_1.append(web_prods_1)
    prods_2.append(web_prods_2)
    prods_3.append(web_prods_3)

# make sure that all trait differences are added to every data point          
def add_all_tdiffs_everywhere(web):
    if web == 0:
        add_dict = {'B_diff': 0, 'I_diff': 0, 'T_diff': 0}
    if web == 1:
        add_dict = {'I_diff': 0, 'T_diff': 0}
    if web == 2:
        add_dict = {'B_diff': 0, 'T_diff': 0}
    if web == 3:
        add_dict = {'T_diff': 0}
    if web == 4:
        add_dict = {'B_diff': 0, 'I_diff': 0}
    if web == 5:
        add_dict = {'I_diff': 0}
    if web == 6:
        add_dict = {'B_diff': 0}
    if web == 7:
        return
    for i in p_d_list[web]:
            i.update(add_dict)
    return

def check_tdiff(tdiff, value):
    if value == '0':
        if tdiff == 0:
            return 1
        else:
            return 0
    if value == 'low':
        if tdiff == 0.25:
            return 1
        elif tdiff == 0.5:
            return 1
        else:
            return 0
    if value == 'high':
        if tdiff == 0.75:
            return 1
        elif tdiff == 1:
            return 1
        else:
            return 0
    
feature_labels = ['B_diff', 'I_diff', 'T_diff', 'N0', 'h_N', 'r0', 'a0', 'h0', 
                'alpha0', 'eta0', 'delta', 'n', 'nu', 'a_scale']

for web in range(8):
    add_all_tdiffs_everywhere(web)

# %%===========================================================================
# "main"
# =============================================================================
# FIGURE S5.2
#range for contour lines
vmin_arr = [320, 24]
vmax_arr = [340, 43]
contourBax = [0, 0.25, 0.5, 0.75, 1]
contourIax = [[0, 0.25, 0.5, 0.75, 1],[0.25, 0.5, 0.75, 1],
              [0.25, 0.5, 0.75, 1]]
I_startindex = [0, 1, 1]
cmapstr = ['Greens', 'Oranges']

fmt_arr = ['%i', '%i'] # for formatting


fig, axes = plt.subplots(2, 3, sharex = 'all', sharey = 'all',
                         figsize = (12, 6))
for level in [0, 1]:
    # first prepare data for random forest algorithm
    function = []
    data_samples = []
    for web in [0, 1, 2, 3, 6, 7]: # do not treat webs 4 and 5
        for i in range(len(p_d_list[web])):
            if 0 in ext[web][i]:
                tdiff = p_d_list[web][i]['T_diff']
                indices = np.where(ext[web][i] == 0)
                for j in indices[0]:
                    function.append(prods_1[web][i][j][level])
                    values = []
                    for key in feature_labels:
                        values.append(p_d_list[web][i][key])
                    data_samples.append(values)
    
    function = np.asarray(function)
    data_samples = np.asarray(data_samples)
    ts = time.time()
    clf = ExtraTreesRegressor(n_estimators = n_trees, bootstrap = 1, 
                              oob_score = 1, n_jobs = num_cores)
    model = clf.fit(data_samples, function)

    # generate 2d partial dependence plot
    data_pd = pd.DataFrame(data = data_samples, 
                              index = range(len(data_samples)),
                              columns = feature_labels)
    
    P1 = calc_pdps(model, data_pd)
    for tdiff in [0, 1, 2]:
        CSf = axes[1 - level, tdiff].contourf(contourBax, contourIax[tdiff], 
                           P1[:, I_startindex[tdiff]:, tdiff].T, 
                           cmap = cmapstr[level],
                           vmin = vmin_arr[level], vmax = vmax_arr[level])

        CS = axes[1 - level, tdiff].contour(contourBax, contourIax[tdiff], 
                         P1[:, I_startindex[tdiff]:, tdiff].T, 
                         colors = 'k',
                         vmin = vmin_arr[level], vmax = vmax_arr[level])


        axes[1 - level, tdiff].set_aspect('equal')
        axes[1 - level, tdiff].clabel(CS, inline = 1, fontsize = 10, fmt = fmt_arr[level])
        
axes[0, 0].set_yticks(contourBax)
axes[0, 0].set_xticks(contourBax)
axes[0, 0].set_ylim([0, 1])
for i in [0, 1]:
    for j in [1, 2]:
        axes[i, j].fill_between([0, 1], 0.25, hatch = 3*'/', edgecolor = 'k', facecolor = 'none')
plt.tight_layout()
fig.subplots_adjust(wspace=-0.52, hspace=0.03)
plt.savefig('fig_S5-2.pdf')

#%% FIGURE S5.3
#range for contour lines
vmin_arr = [95, 20, 0.306, 0.66]
vmax_arr = [155, 35, 0.48, 0.75]
contourBax = [0, 0.25, 0.5, 0.75, 1]
contourIax = [[0, 0.25, 0.5, 0.75, 1],[0.25, 0.5, 0.75, 1],
              [0.25, 0.5, 0.75, 1]]
I_startindex = [0, 1, 1]
cmapstr = ['Greens', 'Oranges', 'Greens', 'Oranges']

fmt_arr = ['%i', '%.1f', '%.3f', '%.3f']

fig, axes = plt.subplots(4, 3, sharex = 'all', sharey = 'all',
                         figsize = (12, 12))
for level in [0, 1, 2, 3]:
    # first prepare data for random forest algorithm
    function = []
    data_samples = []
    for web in [0, 1, 2, 3, 6, 7]: # do not treat webs 4 and 5
        for i in range(len(p_d_list[web])):
            if 0 in ext[web][i]:
                tdiff = p_d_list[web][i]['T_diff']
                indices = np.where(ext[web][i] == 0)
                for j in indices[0]:
                    function.append(prods_2[web][i][j][level])
                    values = []
                    for key in feature_labels:
                        values.append(p_d_list[web][i][key])
                    data_samples.append(values)
    
    function = np.asarray(function)
    data_samples = np.asarray(data_samples)
    ts = time.time()
    clf = ExtraTreesRegressor(n_estimators = n_trees, bootstrap = 1, 
                              oob_score = 1, n_jobs = num_cores)
    model = clf.fit(data_samples, function)

    # generate 2d partial dependence plot
    data_pd = pd.DataFrame(data = data_samples, 
                              index = range(len(data_samples)),
                              columns = feature_labels)
    
    P1 = calc_pdps(model, data_pd)
    for tdiff in [0, 1, 2]:
        CSf = axes[3 - level, tdiff].contourf(contourBax, contourIax[tdiff], 
                           P1[:, I_startindex[tdiff]:, tdiff].T, 
                           cmap = cmapstr[level],
                           vmin = vmin_arr[level], vmax = vmax_arr[level])

        CS = axes[3 - level, tdiff].contour(contourBax, contourIax[tdiff], 
                         P1[:, I_startindex[tdiff]:, tdiff].T, 
                         colors = 'k',
                         vmin = vmin_arr[level], vmax = vmax_arr[level])


        axes[3 - level, tdiff].set_aspect('equal')
        axes[3 - level, tdiff].clabel(CS, inline = 1, fontsize = 10, fmt = fmt_arr[level])
        
axes[0, 0].set_yticks(contourBax)
axes[0, 0].set_xticks(contourBax)
axes[0, 0].set_ylim([0, 1])
for i in [0, 1, 2, 3]:
    for j in [1, 2]:
        axes[i, j].fill_between([0, 1], 0.25, hatch = 3*'/', edgecolor = 'k', facecolor = 'none')
plt.tight_layout()
fig.subplots_adjust(wspace=-0.52, hspace=0.03)
plt.savefig('fig_S5-3.pdf')
#%% figure S5.4

params = np.array(['nu', 'N0', 'eta0', 'delta', 'h0', 'T_diff', 'I_diff', 'n', 
          'alpha0', 'r0', 'B_diff', 'a_scale', 'a0', 'h_N'])

labels_dict = {'nu': r'$\nu$',
                'N0': r'$N_0$',
                'eta0': r'$\eta_0$',
                'delta': r'$\delta$',
                'h0': r'$h_0$', 
                'T_diff': r'$\Delta_T$',
                'I_diff': r'$\Delta_I$', 
                'n': r'$n$', 
                'alpha0': r'$\alpha_0$',
                'r0': r'$r_0$', 
                'B_diff': r'$\Delta_B$',
                'a_scale': r'$a_{sc}$',
                'a0': r'$a_0$',
                'h_N': r'$h_N$'}


imp_dict_prod = {}
label_dict_prod = {}

for level in range(len(levelstr_3)):
    # first prepare data for random forest algorithm
    function = []
    data_samples = []
    for web in [0, 1, 2, 3, 6, 7]: # do not treat webs 4 and 5
        for i in range(len(p_d_list[web])):
            if 0 in ext[web][i]:
                indices = np.where(ext[web][i] == 0)
                for j in indices[0]:
                    function.append(prods_3[web][i][j][level])
                    values = []
                    for key in feature_labels:
                        values.append(p_d_list[web][i][key])
                    data_samples.append(values)
                        
    function = np.asarray(function)
    data_samples = np.asarray(data_samples)
    ts = time.time()
    clf = ExtraTreesRegressor(n_estimators = n_trees, bootstrap = 1, oob_score = 1, n_jobs = num_cores)
    model = clf.fit(data_samples, function)
    importances = model.feature_importances_
    indices = np.argsort(importances)[::-1]
            
    labels = [feature_labels[i] for i in indices]
    
    imp_dict_prod[levelstr_3[level]] = importances[indices]
    label_dict_prod[levelstr_3[level]] = labels

mean_imps_prod = []

for param in labels_dict:
    mean_val = 0
    for metric in levelstr_3:
        param_index = label_dict_prod[metric].index(param)
        mean_val += imp_dict_prod[metric][param_index]
    mean_imps_prod.append(mean_val/11)
mean_imps_prod = np.asarray(mean_imps_prod)  
order = np.argsort(-mean_imps_prod)

xlabels = [labels_dict[param] for param in params[order]]

fig, ax = plt.subplots()
for i in range(len(params)):
    paramstr = params[order][i]
    ax.bar(i, mean_imps_prod[order[i]], color = 'C0')
x = np.arange(len(labels_dict)) # the label locations
ax.set_ylabel('Mean relative importance for all production metrics')
ax.set_xticks(x)
ax.set_xticklabels(xlabels, fontsize = 15)
plt.savefig('fig_S5-4.pdf')


#%% figure S6.1
xlabels = [labels_dict[param] for param in label_dict_prod['PB']]

fig, ax = plt.subplots()
for i in range(len(params)):
    ax.bar(i, imp_dict_prod['PB'][i], color = 'C0')
x = np.arange(len(labels_dict)) # the label locations
ax.set_ylabel(r'Relative importance for $P_B$')
ax.set_xticks(x)
ax.set_xticklabels(xlabels, fontsize = 15)

plt.savefig('fig_S6-1.pdf')
```
