## Supplementary material for "Top predators govern multitrophic diversity effects in tritrophic food webs": Code and data: import_raw_data.py

```
#!/usr/bin/env python3
# -*- coding: utf-8 -*-
"""
Created on Fri Jun 5 2020

@author: ruben
"""
import ast
import numpy as np
import pandas as pd
from pathlib import Path

# =============================================================================
# import raw data
# this function will read the Python data files and put everything in numpy
# arrays
# =============================================================================
def import_data():
    # change the names accordingly to analyze files other than the ones provided
    path_0 = Path("food_web_data/0-data_1024-200.out")
    path_1 = Path("food_web_data/1-data_4096-200.out")
    path_2 = Path("food_web_data/2-data_4096-200.out")
    path_3 = Path("food_web_data/3-data_16384-200.out")
    path_4 = Path("food_web_data/4-data_4096-200.out")
    path_5 = Path("food_web_data/5-data_16384-200.out")
    path_6 = Path("food_web_data/6-data_16384-200.out")
    path_7 = Path("food_web_data/7-data_65536-200.out")

    ### 0 chain
    
    output_arr_0 = pd.read_csv(path_0, sep = ';',
                     header = None)
    # parameter value used
    p_d_list_0 = []
    
    for i in range(len(output_arr_0)):
        p_d_list_0.append(ast.literal_eval(output_arr_0[0][i]))
    
    # # of attractors
    attr_num_0  = output_arr_0[1][:]
    
    # any extinctions?
    ext_0 = []
    for i in output_arr_0[2][:]:
        ext_0.append(np.fromstring(i.replace('[', '').replace(']', '').replace(' ', ''), 
                          sep = ','))
        
    # means at different attractors
    unfor_TLmeans_0 = []
    for i in range(len(output_arr_0)):
        temp_array = np.array(output_arr_0[3][i].replace('[', '').replace(']', '')
                        .replace(' ', '').split(',')).astype(float)
        unfor_TLmeans_0.append(temp_array)
    unfor_TLmeans_0 = np.asarray(unfor_TLmeans_0)
    
    TLmeans_0 = []
    
    for i in range(len(output_arr_0)):
        k = len(unfor_TLmeans_0[i])
        new_list = []
        for j in range(k//4):
            new_list.append(unfor_TLmeans_0[i][4*j:4*(j + 1)])
        TLmeans_0.append(np.asarray(new_list))
    TLmeans_0 = np.asarray(TLmeans_0)
    
    means_0 = TLmeans_0
            
    
    # cvTL at different attractors
    unfor_cvTL_0 = []
    for i in range(len(output_arr_0)):
        temp_array = np.array(output_arr_0[5][i].replace('[', '').replace(']', '')
                        .replace(' ', '').split(',')).astype(float)
        unfor_cvTL_0.append(temp_array)
    unfor_cvTL_0 = np.asarray(unfor_cvTL_0)
    
    cvTLs_0 = []
    
    for i in range(len(output_arr_0)):
        k = len(unfor_cvTL_0[i])
        new_list = []
        for j in range(k//4):
            new_list.append(unfor_cvTL_0[i][4*j:4*(j + 1)])
        cvTLs_0.append(np.asarray(new_list))
    cvTLs_0 = np.asarray(cvTLs_0)
        
    # relative frequencies of attractors
    freqs_0 = []
    for i in range(len(output_arr_0)):
        freqs_0.append(np.array(output_arr_0[8][i].replace('[', '').replace(']', '')
                                .replace(' ', '').split(',')).astype(float))
    freqs_0 = np.asarray(freqs_0)
    
    ### 1 B
    
    output_arr_1 = pd.read_csv(path_1, sep = ';',
                    header = None)
    # parameter value used
    p_d_list_1 = []
    
    for i in range(len(output_arr_1)):
        p_d_list_1.append(ast.literal_eval(output_arr_1[0][i]))
    
    # # of attractors
    attr_num_1  = output_arr_1[1][:]
    
    # coexistence of all species?
    ext_1 = []
    for i in output_arr_1[2][:]:
        ext_1.append(np.fromstring(i.replace('[', '').replace(']', '').replace(' ', ''), 
                          sep = ','))
        
    # means at different attractors
    unfor_TLmeans_1 = []
    for i in range(len(output_arr_1)):
        temp_array = np.array(output_arr_1[3][i].replace('[', '').replace(']', '')
                        .replace(' ', '').split(',')).astype(float)
        unfor_TLmeans_1.append(temp_array)
    unfor_TLmeans_1 = np.asarray(unfor_TLmeans_1)
    
    TLmeans_1 = []
    
    for i in range(len(output_arr_1)):
        k = len(unfor_TLmeans_1[i])
        new_list = []
        for j in range(k//4):
            new_list.append(unfor_TLmeans_1[i][4*j:4*(j + 1)])
        TLmeans_1.append(np.asarray(new_list))
    TLmeans_1 = np.asarray(TLmeans_1)
    
    unfor_leftmeans_1 = []
    for i in range(len(output_arr_1)):
        temp_array = np.array(output_arr_1[4][i].replace('[', '').replace(']', '')
                        .replace(' ', '').split(',')).astype(float)
        unfor_leftmeans_1.append(temp_array)
    unfor_leftmeans_1 = np.asarray(unfor_leftmeans_1)
    
    leftmeans_1 = []
    
    for i in range(len(output_arr_1)):
        k = len(unfor_leftmeans_1[i])
        new_list = []
        for j in range(k//3):
            new_list.append(unfor_leftmeans_1[i][3*j:3*(j + 1)])
        leftmeans_1.append(np.asarray(new_list))
    leftmeans_1 = np.asarray(leftmeans_1)
    
    means_1 = []
    for i in range(len(output_arr_1)):
        attr_arr = []
        for j in range(len(TLmeans_1[i])):
            meansarr = []
            meansarr.append(TLmeans_1[i][j][0])
            meansarr.append(leftmeans_1[i][j][0])
            meansarr.append(TLmeans_1[i][j][1] - leftmeans_1[i][j][0])
            meansarr.append(TLmeans_1[i][j][2])
            meansarr.append(TLmeans_1[i][j][3])
            attr_arr.append(np.asarray(meansarr))
        means_1.append(np.asarray(attr_arr))
    means_1 = np.asarray(means_1)
    
    # cvTL at different attractors
    unfor_cvTL_1 = []
    for i in range(len(output_arr_1)):
        temp_array = np.array(output_arr_1[5][i].replace('[', '').replace(']', '')
                        .replace(' ', '').split(',')).astype(float)
        unfor_cvTL_1.append(temp_array)
    unfor_cvTL_1 = np.asarray(unfor_cvTL_1)
    
    cvTLs_1 = []
    
    for i in range(len(output_arr_1)):
        k = len(unfor_cvTL_1[i])
        new_list = []
        for j in range(k//4):
            new_list.append(unfor_cvTL_1[i][4*j:4*(j + 1)])
        cvTLs_1.append(np.asarray(new_list))
    cvTLs_1 = np.asarray(cvTLs_1)
    
    unfor_leftCVs_1 = []
    for i in range(len(output_arr_1)):
        temp_array = np.array(output_arr_1[6][i].replace('[', '').replace(']', '')
                        .replace(' ', '').split(',')).astype(float)
        unfor_leftCVs_1.append(temp_array)
    unfor_leftCVs_1 = np.asarray(unfor_leftCVs_1)
    
    leftCVs_1 = []
    
    for i in range(len(output_arr_1)):
        k = len(unfor_leftCVs_1[i])
        new_list = []
        for j in range(k//3):
            new_list.append(unfor_leftCVs_1[i][3*j:3*(j + 1)])
        leftCVs_1.append(np.asarray(new_list))
    leftCVs_1 = np.asarray(leftCVs_1)
    
    unfor_rightCVs_1 = []
    for i in range(len(output_arr_1)):
        temp_array = np.array(output_arr_1[7][i].replace('[', '').replace(']', '')
                        .replace(' ', '').split(',')).astype(float)
        unfor_rightCVs_1.append(temp_array)
    unfor_rightCVs_1 = np.asarray(unfor_rightCVs_1)
    
    rightCVs_1 = []
    
    for i in range(len(output_arr_1)):
        k = len(unfor_rightCVs_1[i])
        new_list = []
        for j in range(k//3):
            new_list.append(unfor_rightCVs_1[i][3*j:3*(j + 1)])
        rightCVs_1.append(np.asarray(new_list))
    rightCVs_1 = np.asarray(rightCVs_1)
    
    CVs_1 = []
    for i in range(len(output_arr_1)):
        attr_arr = []
        for j in range(len(cvTLs_1[i])):
            CVarr = []
            CVarr.append(cvTLs_1[i][j][0])
            CVarr.append(leftCVs_1[i][j][0])
            CVarr.append(rightCVs_1[i][j][0])
            CVarr.append(cvTLs_1[i][j][2])
            CVarr.append(cvTLs_1[i][j][3])
            attr_arr.append(np.asarray(CVarr))
        CVs_1.append(np.asarray(attr_arr))
    CVs_1 = np.asarray(CVs_1)
    
    # relative frequencies of attractors
    freqs_1 = []
    for i in range(len(output_arr_1)):
        freqs_1.append(np.array(output_arr_1[8][i].replace('[', '').replace(']', '')
                                .replace(' ', '').split(',')).astype(float))
    freqs_1 = np.asarray(freqs_1)
    
    ### 2 I
    
    output_arr_2 = pd.read_csv(path_2, sep = ';',
                    header = None)
    # parameter value used
    p_d_list_2 = []
    
    for i in range(len(output_arr_2)):
        p_d_list_2.append(ast.literal_eval(output_arr_2[0][i]))
    
    # # of attractors
    attr_num_2  = output_arr_2[1][:]
    
    # coexistence of all species?
    ext_2 = []
    for i in output_arr_2[2][:]:
        ext_2.append(np.fromstring(i.replace('[', '').replace(']', '').replace(' ', ''), 
                          sep = ','))
        
    # means at different attractors
    unfor_TLmeans_2 = []
    for i in range(len(output_arr_2)):
        temp_array = np.array(output_arr_2[3][i].replace('[', '').replace(']', '')
                        .replace(' ', '').split(',')).astype(float)
        unfor_TLmeans_2.append(temp_array)
    unfor_TLmeans_2 = np.asarray(unfor_TLmeans_2)
    
    TLmeans_2 = []
    
    for i in range(len(output_arr_2)):
        k = len(unfor_TLmeans_2[i])
        new_list = []
        for j in range(k//4):
            new_list.append(unfor_TLmeans_2[i][4*j:4*(j + 1)])
        TLmeans_2.append(np.asarray(new_list))
    TLmeans_2 = np.asarray(TLmeans_2)
    
    unfor_leftmeans_2 = []
    for i in range(len(output_arr_2)):
        temp_array = np.array(output_arr_2[4][i].replace('[', '').replace(']', '')
                        .replace(' ', '').split(',')).astype(float)
        unfor_leftmeans_2.append(temp_array)
    unfor_leftmeans_2 = np.asarray(unfor_leftmeans_2)
    
    leftmeans_2 = []
    
    for i in range(len(output_arr_2)):
        k = len(unfor_leftmeans_2[i])
        new_list = []
        for j in range(k//3):
            new_list.append(unfor_leftmeans_2[i][3*j:3*(j + 1)])
        leftmeans_2.append(np.asarray(new_list))
    leftmeans_2 = np.asarray(leftmeans_2)
    
    means_2 = []
    for i in range(len(output_arr_2)):
        attr_arr = []
        for j in range(len(TLmeans_2[i])):
            meansarr = []
            meansarr.append(TLmeans_2[i][j][0])
            meansarr.append(TLmeans_2[i][j][1])
            meansarr.append(leftmeans_2[i][j][1])
            meansarr.append(TLmeans_2[i][j][2] - leftmeans_2[i][j][1])
            meansarr.append(TLmeans_2[i][j][3])
            attr_arr.append(np.asarray(meansarr))
        means_2.append(np.asarray(attr_arr))
    means_2 = np.asarray(means_2)
    
    # cvTL at different attractors
    unfor_cvTL_2 = []
    for i in range(len(output_arr_2)):
        temp_array = np.array(output_arr_2[5][i].replace('[', '').replace(']', '')
                        .replace(' ', '').split(',')).astype(float)
        unfor_cvTL_2.append(temp_array)
    unfor_cvTL_2 = np.asarray(unfor_cvTL_2)
    
    cvTLs_2 = []
    
    for i in range(len(output_arr_2)):
        k = len(unfor_cvTL_2[i])
        new_list = []
        for j in range(k//4):
            new_list.append(unfor_cvTL_2[i][4*j:4*(j + 1)])
        cvTLs_2.append(np.asarray(new_list))
    cvTLs_2 = np.asarray(cvTLs_2)
    
    unfor_leftCVs_2 = []
    for i in range(len(output_arr_2)):
        temp_array = np.array(output_arr_2[6][i].replace('[', '').replace(']', '')
                        .replace(' ', '').split(',')).astype(float)
        unfor_leftCVs_2.append(temp_array)
    unfor_leftCVs_2 = np.asarray(unfor_leftCVs_2)
    
    leftCVs_2 = []
    
    for i in range(len(output_arr_2)):
        k = len(unfor_leftCVs_2[i])
        new_list = []
        for j in range(k//3):
            new_list.append(unfor_leftCVs_2[i][3*j:3*(j + 1)])
        leftCVs_2.append(np.asarray(new_list))
    leftCVs_2 = np.asarray(leftCVs_2)
    
    unfor_rightCVs_2 = []
    for i in range(len(output_arr_2)):
        temp_array = np.array(output_arr_2[7][i].replace('[', '').replace(']', '')
                        .replace(' ', '').split(',')).astype(float)
        unfor_rightCVs_2.append(temp_array)
    unfor_rightCVs_2 = np.asarray(unfor_rightCVs_2)
    
    rightCVs_2 = []
    
    for i in range(len(output_arr_2)):
        k = len(unfor_rightCVs_2[i])
        new_list = []
        for j in range(k//3):
            new_list.append(unfor_rightCVs_2[i][3*j:3*(j + 1)])
        rightCVs_2.append(np.asarray(new_list))
    rightCVs_2 = np.asarray(rightCVs_2)
    
    CVs_2 = []
    for i in range(len(output_arr_2)):
        attr_arr = []
        for j in range(len(cvTLs_2[i])):
            CVarr = []
            CVarr.append(cvTLs_2[i][j][0])
            CVarr.append(cvTLs_2[i][j][0])
            CVarr.append(leftCVs_2[i][j][1])
            CVarr.append(rightCVs_2[i][j][1])
            CVarr.append(cvTLs_2[i][j][3])
            attr_arr.append(np.asarray(CVarr))
        CVs_2.append(np.asarray(attr_arr))
    CVs_2 = np.asarray(CVs_2)
    
    # relative frequencies of attractors
    freqs_2 = []
    for i in range(len(output_arr_2)):
        freqs_2.append(np.array(output_arr_2[8][i].replace('[', '').replace(']', '')
                                .replace(' ', '').split(',')).astype(float))
    freqs_2 = np.asarray(freqs_2)
    
    
    ### 3 BI
    
    output_arr_3 = pd.read_csv(path_3, sep = ';',
                    header = None)
    # parameter value used
    p_d_list_3 = []
    
    for i in range(len(output_arr_3)):
        p_d_list_3.append(ast.literal_eval(output_arr_3[0][i]))
    
    # # of attractors
    attr_num_3  = output_arr_3[1][:]
    
    # coexistence of all species?
    ext_3 = []
    for i in output_arr_3[2][:]:
        ext_3.append(np.fromstring(i.replace('[', '').replace(']', '').replace(' ', ''), 
                          sep = ','))
        
    # means at different attractors
    unfor_TLmeans_3 = []
    for i in range(len(output_arr_3)):
        temp_array = np.array(output_arr_3[3][i].replace('[', '').replace(']', '')
                        .replace(' ', '').split(',')).astype(float)
        unfor_TLmeans_3.append(temp_array)
    unfor_TLmeans_3 = np.asarray(unfor_TLmeans_3)
    
    TLmeans_3 = []
    
    for i in range(len(output_arr_3)):
        k = len(unfor_TLmeans_3[i])
        new_list = []
        for j in range(k//4):
            new_list.append(unfor_TLmeans_3[i][4*j:4*(j + 1)])
        TLmeans_3.append(np.asarray(new_list))
    TLmeans_3 = np.asarray(TLmeans_3)
    
    unfor_leftmeans_3 = []
    for i in range(len(output_arr_3)):
        temp_array = np.array(output_arr_3[4][i].replace('[', '').replace(']', '')
                        .replace(' ', '').split(',')).astype(float)
        unfor_leftmeans_3.append(temp_array)
    unfor_leftmeans_3 = np.asarray(unfor_leftmeans_3)
    
    leftmeans_3 = []
    
    for i in range(len(output_arr_3)):
        k = len(unfor_leftmeans_3[i])
        new_list = []
        for j in range(k//3):
            new_list.append(unfor_leftmeans_3[i][3*j:3*(j + 1)])
        leftmeans_3.append(np.asarray(new_list))
    leftmeans_3 = np.asarray(leftmeans_3)
    
    means_3 = []
    for i in range(len(output_arr_3)):
        attr_arr = []
        for j in range(len(TLmeans_3[i])):
            meansarr = []
            meansarr.append(TLmeans_3[i][j][0])
            meansarr.append(leftmeans_3[i][j][0])
            meansarr.append(TLmeans_3[i][j][1] - leftmeans_3[i][j][0])
            meansarr.append(leftmeans_3[i][j][1])
            meansarr.append(TLmeans_3[i][j][2] - leftmeans_3[i][j][1])
            meansarr.append(TLmeans_3[i][j][3])
            attr_arr.append(np.asarray(meansarr))
        means_3.append(np.asarray(attr_arr))
    means_3 = np.asarray(means_3)
    
    # cvTL at different attractors
    unfor_cvTL_3 = []
    for i in range(len(output_arr_3)):
        temp_array = np.array(output_arr_3[5][i].replace('[', '').replace(']', '')
                        .replace(' ', '').split(',')).astype(float)
        unfor_cvTL_3.append(temp_array)
    unfor_cvTL_3 = np.asarray(unfor_cvTL_3)
    
    cvTLs_3 = []
    
    for i in range(len(output_arr_3)):
        k = len(unfor_cvTL_3[i])
        new_list = []
        for j in range(k//4):
            new_list.append(unfor_cvTL_3[i][4*j:4*(j + 1)])
        cvTLs_3.append(np.asarray(new_list))
    cvTLs_3 = np.asarray(cvTLs_3)
    
    unfor_leftCVs_3 = []
    for i in range(len(output_arr_3)):
        temp_array = np.array(output_arr_3[6][i].replace('[', '').replace(']', '')
                        .replace(' ', '').split(',')).astype(float)
        unfor_leftCVs_3.append(temp_array)
    unfor_leftCVs_3 = np.asarray(unfor_leftCVs_3)
    
    leftCVs_3 = []
    
    for i in range(len(output_arr_3)):
        k = len(unfor_leftCVs_3[i])
        new_list = []
        for j in range(k//3):
            new_list.append(unfor_leftCVs_3[i][3*j:3*(j + 1)])
        leftCVs_3.append(np.asarray(new_list))
    leftCVs_3 = np.asarray(leftCVs_3)
    
    unfor_rightCVs_3 = []
    for i in range(len(output_arr_3)):
        temp_array = np.array(output_arr_3[7][i].replace('[', '').replace(']', '')
                        .replace(' ', '').split(',')).astype(float)
        unfor_rightCVs_3.append(temp_array)
    unfor_rightCVs_3 = np.asarray(unfor_rightCVs_3)
    
    rightCVs_3 = []
    
    for i in range(len(output_arr_3)):
        k = len(unfor_rightCVs_3[i])
        new_list = []
        for j in range(k//3):
            new_list.append(unfor_rightCVs_3[i][3*j:3*(j + 1)])
        rightCVs_3.append(np.asarray(new_list))
    rightCVs_3 = np.asarray(rightCVs_3)
    
    CVs_3 = []
    for i in range(len(output_arr_3)):
        attr_arr = []
        for j in range(len(cvTLs_3[i])):
            CVarr = []
            CVarr.append(cvTLs_3[i][j][0])
            CVarr.append(leftCVs_3[i][j][0])
            CVarr.append(rightCVs_3[i][j][0])
            CVarr.append(leftCVs_3[i][j][1])
            CVarr.append(rightCVs_3[i][j][1])
            CVarr.append(cvTLs_3[i][j][3])
            attr_arr.append(np.asarray(CVarr))
        CVs_3.append(np.asarray(attr_arr))
    CVs_3 = np.asarray(CVs_3)
    
    # relative frequencies of attractors
    freqs_3 = []
    for i in range(len(output_arr_3)):
        freqs_3.append(np.array(output_arr_3[8][i].replace('[', '').replace(']', '')
                                .replace(' ', '').split(',')).astype(float))
    freqs_3 = np.asarray(freqs_3)
    
                            
    ### 4 T
    
    output_arr_4 = pd.read_csv(path_4, sep = ';',
                    header = None)
    # parameter value used
    p_d_list_4 = []
    
    for i in range(len(output_arr_4)):
        p_d_list_4.append(ast.literal_eval(output_arr_4[0][i]))
    
    # # of attractors
    attr_num_4  = output_arr_4[1][:]
    
    # coexistence of all species?
    ext_4 = []
    for i in output_arr_4[2][:]:
        ext_4.append(np.fromstring(i.replace('[', '').replace(']', '').replace(' ', ''), 
                          sep = ','))
        
    # means at different attractors
    unfor_TLmeans_4 = []
    for i in range(len(output_arr_4)):
        temp_array = np.array(output_arr_4[3][i].replace('[', '').replace(']', '')
                        .replace(' ', '').split(',')).astype(float)
        unfor_TLmeans_4.append(temp_array)
    unfor_TLmeans_4 = np.asarray(unfor_TLmeans_4)
    
    TLmeans_4 = []
    
    for i in range(len(output_arr_4)):
        k = len(unfor_TLmeans_4[i])
        new_list = []
        for j in range(k//4):
            new_list.append(unfor_TLmeans_4[i][4*j:4*(j + 1)])
        TLmeans_4.append(np.asarray(new_list))
    TLmeans_4 = np.asarray(TLmeans_4)
    
    unfor_leftmeans_4 = []
    for i in range(len(output_arr_4)):
        temp_array = np.array(output_arr_4[4][i].replace('[', '').replace(']', '')
                        .replace(' ', '').split(',')).astype(float)
        unfor_leftmeans_4.append(temp_array)
    unfor_leftmeans_4 = np.asarray(unfor_leftmeans_4)
    
    leftmeans_4 = []
    
    for i in range(len(output_arr_4)):
        k = len(unfor_leftmeans_4[i])
        new_list = []
        for j in range(k//3):
            new_list.append(unfor_leftmeans_4[i][3*j:3*(j + 1)])
        leftmeans_4.append(np.asarray(new_list))
    leftmeans_4 = np.asarray(leftmeans_4)
    
    means_4 = []
    for i in range(len(output_arr_4)):
        attr_arr = []
        for j in range(len(TLmeans_4[i])):
            meansarr = []
            meansarr.append(TLmeans_4[i][j][0])
            meansarr.append(TLmeans_4[i][j][1])
            meansarr.append(TLmeans_4[i][j][2])
            meansarr.append(leftmeans_4[i][j][2])
            meansarr.append(TLmeans_4[i][j][3] - leftmeans_4[i][j][2])
            attr_arr.append(np.asarray(meansarr))
        means_4.append(np.asarray(attr_arr))
    means_4 = np.asarray(means_4)
    
    # cvTL at different attractors
    unfor_cvTL_4 = []
    for i in range(len(output_arr_4)):
        temp_array = np.array(output_arr_4[5][i].replace('[', '').replace(']', '')
                        .replace(' ', '').split(',')).astype(float)
        unfor_cvTL_4.append(temp_array)
    unfor_cvTL_4 = np.asarray(unfor_cvTL_4)
    
    cvTLs_4 = []
    
    for i in range(len(output_arr_4)):
        k = len(unfor_cvTL_4[i])
        new_list = []
        for j in range(k//4):
            new_list.append(unfor_cvTL_4[i][4*j:4*(j + 1)])
        cvTLs_4.append(np.asarray(new_list))
    cvTLs_4 = np.asarray(cvTLs_4)
    
    unfor_leftCVs_4 = []
    for i in range(len(output_arr_4)):
        temp_array = np.array(output_arr_4[6][i].replace('[', '').replace(']', '')
                        .replace(' ', '').split(',')).astype(float)
        unfor_leftCVs_4.append(temp_array)
    unfor_leftCVs_4 = np.asarray(unfor_leftCVs_4)
    
    leftCVs_4 = []
    
    for i in range(len(output_arr_4)):
        k = len(unfor_leftCVs_4[i])
        new_list = []
        for j in range(k//3):
            new_list.append(unfor_leftCVs_4[i][3*j:3*(j + 1)])
        leftCVs_4.append(np.asarray(new_list))
    leftCVs_4 = np.asarray(leftCVs_4)
    
    unfor_rightCVs_4 = []
    for i in range(len(output_arr_4)):
        temp_array = np.array(output_arr_4[7][i].replace('[', '').replace(']', '')
                        .replace(' ', '').split(',')).astype(float)
        unfor_rightCVs_4.append(temp_array)
    unfor_rightCVs_4 = np.asarray(unfor_rightCVs_4)
    
    rightCVs_4 = []
    
    for i in range(len(output_arr_4)):
        k = len(unfor_rightCVs_4[i])
        new_list = []
        for j in range(k//3):
            new_list.append(unfor_rightCVs_4[i][3*j:3*(j + 1)])
        rightCVs_4.append(np.asarray(new_list))
    rightCVs_4 = np.asarray(rightCVs_4)
    
    CVs_4 = []
    for i in range(len(output_arr_4)):
        attr_arr = []
        for j in range(len(cvTLs_4[i])):
            CVarr = []
            CVarr.append(cvTLs_4[i][j][0])
            CVarr.append(cvTLs_4[i][j][1])
            CVarr.append(cvTLs_4[i][j][2])
            CVarr.append(leftCVs_4[i][j][2])
            CVarr.append(rightCVs_4[i][j][2])
            attr_arr.append(np.asarray(CVarr))
        CVs_4.append(np.asarray(attr_arr))
    CVs_4 = np.asarray(CVs_4)
    
    # relative frequencies of attractors
    freqs_4 = []
    for i in range(len(output_arr_4)):
        freqs_4.append(np.array(output_arr_4[8][i].replace('[', '').replace(']', '')
                                .replace(' ', '').split(',')).astype(float))
    freqs_4 = np.asarray(freqs_4)
    
    
    ### 5 BT
    
    output_arr_5 = pd.read_csv(path_5, sep = ';',
                    header = None)
    # parameter value used
    p_d_list_5 = []
    
    for i in range(len(output_arr_5)):
        p_d_list_5.append(ast.literal_eval(output_arr_5[0][i]))
    
    # # of attractors
    attr_num_5  = output_arr_5[1][:]
    
    # coexistence of all species?
    ext_5 = []
    for i in output_arr_5[2][:]:
        ext_5.append(np.fromstring(i.replace('[', '').replace(']', '').replace(' ', ''), 
                          sep = ','))
        
    # means at different attractors
    unfor_TLmeans_5 = []
    for i in range(len(output_arr_5)):
        temp_array = np.array(output_arr_5[3][i].replace('[', '').replace(']', '')
                        .replace(' ', '').split(',')).astype(float)
        unfor_TLmeans_5.append(temp_array)
    unfor_TLmeans_5 = np.asarray(unfor_TLmeans_5)
    
    TLmeans_5 = []
    
    for i in range(len(output_arr_5)):
        k = len(unfor_TLmeans_5[i])
        new_list = []
        for j in range(k//4):
            new_list.append(unfor_TLmeans_5[i][4*j:4*(j + 1)])
        TLmeans_5.append(np.asarray(new_list))
    TLmeans_5 = np.asarray(TLmeans_5)
    
    unfor_leftmeans_5 = []
    for i in range(len(output_arr_5)):
        temp_array = np.array(output_arr_5[4][i].replace('[', '').replace(']', '')
                        .replace(' ', '').split(',')).astype(float)
        unfor_leftmeans_5.append(temp_array)
    unfor_leftmeans_5 = np.asarray(unfor_leftmeans_5)
    
    leftmeans_5 = []
    
    for i in range(len(output_arr_5)):
        k = len(unfor_leftmeans_5[i])
        new_list = []
        for j in range(k//3):
            new_list.append(unfor_leftmeans_5[i][3*j:3*(j + 1)])
        leftmeans_5.append(np.asarray(new_list))
    leftmeans_5 = np.asarray(leftmeans_5)
    
    means_5 = []
    for i in range(len(output_arr_5)):
        attr_arr = []
        for j in range(len(TLmeans_5[i])):
            meansarr = []
            meansarr.append(TLmeans_5[i][j][0])
            meansarr.append(leftmeans_5[i][j][0])
            meansarr.append(TLmeans_5[i][j][1] - leftmeans_5[i][j][0])
            meansarr.append(TLmeans_5[i][j][2])
            meansarr.append(leftmeans_5[i][j][2])
            meansarr.append(TLmeans_5[i][j][3] - leftmeans_5[i][j][2])
            attr_arr.append(np.asarray(meansarr))
        means_5.append(np.asarray(attr_arr))
    means_5 = np.asarray(means_5)
    
    # cvTL at different attractors
    unfor_cvTL_5 = []
    for i in range(len(output_arr_5)):
        temp_array = np.array(output_arr_5[5][i].replace('[', '').replace(']', '')
                        .replace(' ', '').split(',')).astype(float)
        unfor_cvTL_5.append(temp_array)
    unfor_cvTL_5 = np.asarray(unfor_cvTL_5)
    
    cvTLs_5 = []
    
    for i in range(len(output_arr_5)):
        k = len(unfor_cvTL_5[i])
        new_list = []
        for j in range(k//4):
            new_list.append(unfor_cvTL_5[i][4*j:4*(j + 1)])
        cvTLs_5.append(np.asarray(new_list))
    cvTLs_5 = np.asarray(cvTLs_5)
    
    unfor_leftCVs_5 = []
    for i in range(len(output_arr_5)):
        temp_array = np.array(output_arr_5[6][i].replace('[', '').replace(']', '')
                        .replace(' ', '').split(',')).astype(float)
        unfor_leftCVs_5.append(temp_array)
    unfor_leftCVs_5 = np.asarray(unfor_leftCVs_5)
    
    leftCVs_5 = []
    
    for i in range(len(output_arr_5)):
        k = len(unfor_leftCVs_5[i])
        new_list = []
        for j in range(k//3):
            new_list.append(unfor_leftCVs_5[i][3*j:3*(j + 1)])
        leftCVs_5.append(np.asarray(new_list))
    leftCVs_5 = np.asarray(leftCVs_5)
    
    unfor_rightCVs_5 = []
    for i in range(len(output_arr_5)):
        temp_array = np.array(output_arr_5[7][i].replace('[', '').replace(']', '')
                        .replace(' ', '').split(',')).astype(float)
        unfor_rightCVs_5.append(temp_array)
    unfor_rightCVs_5 = np.asarray(unfor_rightCVs_5)
    
    rightCVs_5 = []
    
    for i in range(len(output_arr_5)):
        k = len(unfor_rightCVs_5[i])
        new_list = []
        for j in range(k//3):
            new_list.append(unfor_rightCVs_5[i][3*j:3*(j + 1)])
        rightCVs_5.append(np.asarray(new_list))
    rightCVs_5 = np.asarray(rightCVs_5)
    
    CVs_5 = []
    for i in range(len(output_arr_5)):
        attr_arr = []
        for j in range(len(cvTLs_5[i])):
            CVarr = []
            CVarr.append(cvTLs_5[i][j][0])
            CVarr.append(leftCVs_5[i][j][0])
            CVarr.append(rightCVs_5[i][j][0])
            CVarr.append(cvTLs_5[i][j][2])
            CVarr.append(leftCVs_5[i][j][2])
            CVarr.append(rightCVs_5[i][j][2])
            attr_arr.append(np.asarray(CVarr))
        CVs_5.append(np.asarray(attr_arr))
    CVs_5 = np.asarray(CVs_5)
    
    # relative frequencies of attractors
    freqs_5 = []
    for i in range(len(output_arr_5)):
        freqs_5.append(np.array(output_arr_5[8][i].replace('[', '').replace(']', '')
                                .replace(' ', '').split(',')).astype(float))
    freqs_5 = np.asarray(freqs_5)
    
                            
                            
    ### 6 IT
    
    output_arr_6 = pd.read_csv(path_6, sep = ';',
                    header = None)
    # parameter value used
    p_d_list_6 = []
    
    for i in range(len(output_arr_6)):
        p_d_list_6.append(ast.literal_eval(output_arr_6[0][i]))
    
    # # of attractors
    attr_num_6  = output_arr_6[1][:]
    
    # coexistence of all species?
    ext_6 = []
    for i in output_arr_6[2][:]:
        ext_6.append(np.fromstring(i.replace('[', '').replace(']', '').replace(' ', ''), 
                          sep = ','))
        
    # means at different attractors
    unfor_TLmeans_6 = []
    for i in range(len(output_arr_6)):
        temp_array = np.array(output_arr_6[3][i].replace('[', '').replace(']', '')
                        .replace(' ', '').split(',')).astype(float)
        unfor_TLmeans_6.append(temp_array)
    unfor_TLmeans_6 = np.asarray(unfor_TLmeans_6)
    
    TLmeans_6 = []
    
    for i in range(len(output_arr_6)):
        k = len(unfor_TLmeans_6[i])
        new_list = []
        for j in range(k//4):
            new_list.append(unfor_TLmeans_6[i][4*j:4*(j + 1)])
        TLmeans_6.append(np.asarray(new_list))
    TLmeans_6 = np.asarray(TLmeans_6)
    
    unfor_leftmeans_6 = []
    for i in range(len(output_arr_6)):
        temp_array = np.array(output_arr_6[4][i].replace('[', '').replace(']', '')
                        .replace(' ', '').split(',')).astype(float)
        unfor_leftmeans_6.append(temp_array)
    unfor_leftmeans_6 = np.asarray(unfor_leftmeans_6)
    
    leftmeans_6 = []
    
    for i in range(len(output_arr_6)):
        k = len(unfor_leftmeans_6[i])
        new_list = []
        for j in range(k//3):
            new_list.append(unfor_leftmeans_6[i][3*j:3*(j + 1)])
        leftmeans_6.append(np.asarray(new_list))
    leftmeans_6 = np.asarray(leftmeans_6)
    
    means_6 = []
    for i in range(len(output_arr_6)):
        attr_arr = []
        for j in range(len(TLmeans_6[i])):
            meansarr = []
            meansarr.append(TLmeans_6[i][j][0])
            meansarr.append(TLmeans_6[i][j][1])
            meansarr.append(leftmeans_6[i][j][1])                
            meansarr.append(TLmeans_6[i][j][2] - leftmeans_6[i][j][1])
            meansarr.append(leftmeans_6[i][j][2])
            meansarr.append(TLmeans_6[i][j][3] - leftmeans_6[i][j][2])
            attr_arr.append(np.asarray(meansarr))
        means_6.append(np.asarray(attr_arr))
    means_6 = np.asarray(means_6)
    
    
    # cvTL at different attractors
    unfor_cvTL_6 = []
    for i in range(len(output_arr_6)):
        temp_array = np.array(output_arr_6[5][i].replace('[', '').replace(']', '')
                        .replace(' ', '').split(',')).astype(float)
        unfor_cvTL_6.append(temp_array)
    unfor_cvTL_6 = np.asarray(unfor_cvTL_6)
    
    cvTLs_6 = []
    
    for i in range(len(output_arr_6)):
        k = len(unfor_cvTL_6[i])
        new_list = []
        for j in range(k//4):
            new_list.append(unfor_cvTL_6[i][4*j:4*(j + 1)])
        cvTLs_6.append(np.asarray(new_list))
    cvTLs_6 = np.asarray(cvTLs_6)
    
    unfor_leftCVs_6 = []
    for i in range(len(output_arr_6)):
        temp_array = np.array(output_arr_6[6][i].replace('[', '').replace(']', '')
                        .replace(' ', '').split(',')).astype(float)
        unfor_leftCVs_6.append(temp_array)
    unfor_leftCVs_6 = np.asarray(unfor_leftCVs_6)
    
    leftCVs_6 = []
    
    for i in range(len(output_arr_6)):
        k = len(unfor_leftCVs_6[i])
        new_list = []
        for j in range(k//3):
            new_list.append(unfor_leftCVs_6[i][3*j:3*(j + 1)])
        leftCVs_6.append(np.asarray(new_list))
    leftCVs_6 = np.asarray(leftCVs_6)
    
    unfor_rightCVs_6 = []
    for i in range(len(output_arr_6)):
        temp_array = np.array(output_arr_6[7][i].replace('[', '').replace(']', '')
                        .replace(' ', '').split(',')).astype(float)
        unfor_rightCVs_6.append(temp_array)
    unfor_rightCVs_6 = np.asarray(unfor_rightCVs_6)
    
    rightCVs_6 = []
    
    for i in range(len(output_arr_6)):
        k = len(unfor_rightCVs_6[i])
        new_list = []
        for j in range(k//3):
            new_list.append(unfor_rightCVs_6[i][3*j:3*(j + 1)])
        rightCVs_6.append(np.asarray(new_list))
    rightCVs_6 = np.asarray(rightCVs_6)
    
    CVs_6 = []
    for i in range(len(output_arr_6)):
        attr_arr = []
        for j in range(len(cvTLs_6[i])):
            CVarr = []
            CVarr.append(cvTLs_6[i][j][0])
            CVarr.append(cvTLs_6[i][j][1])        
            CVarr.append(leftCVs_6[i][j][1])
            CVarr.append(rightCVs_6[i][j][1])
            CVarr.append(leftCVs_6[i][j][2])
            CVarr.append(rightCVs_6[i][j][2])
            attr_arr.append(np.asarray(CVarr))
        CVs_6.append(np.asarray(attr_arr))
    CVs_6 = np.asarray(CVs_6)
    
    # relative frequencies of attractors
    freqs_6 = []
    for i in range(len(output_arr_6)):
        freqs_6.append(np.array(output_arr_6[8][i].replace('[', '').replace(']', '')
                                .replace(' ', '').split(',')).astype(float))
    freqs_6 = np.asarray(freqs_6)
    
                                          
                            
                            
    ### 7 BIT
    
    output_arr_7 = pd.read_csv(path_7, sep = ';', header = None)
    # parameter value used
    p_d_list_7 = []
    
    for i in range(len(output_arr_7)):
        p_d_list_7.append(ast.literal_eval(output_arr_7[0][i]))
    
    # # of attractors
    attr_num_7  = output_arr_7[1][:]
    
    # coexistence of all species?
    ext_7 = []
    for i in output_arr_7[2][:]:
        ext_7.append(np.fromstring(i.replace('[', '').replace(']', '').replace(' ', ''), 
                          sep = ','))
        
    # means at different attractors
    unfor_TLmeans_7 = []
    for i in range(len(output_arr_7)):
        temp_array = np.array(output_arr_7[3][i].replace('[', '').replace(']', '')
                        .replace(' ', '').split(',')).astype(float)
        unfor_TLmeans_7.append(temp_array)
    unfor_TLmeans_7 = np.asarray(unfor_TLmeans_7)
    
    TLmeans_7 = []
    
    for i in range(len(output_arr_7)):
        k = len(unfor_TLmeans_7[i])
        new_list = []
        for j in range(k//4):
            new_list.append(unfor_TLmeans_7[i][4*j:4*(j + 1)])
        TLmeans_7.append(np.asarray(new_list))
    TLmeans_7 = np.asarray(TLmeans_7)
    
    unfor_leftmeans_7 = []
    for i in range(len(output_arr_7)):
        temp_array = np.array(output_arr_7[4][i].replace('[', '').replace(']', '')
                        .replace(' ', '').split(',')).astype(float)
        unfor_leftmeans_7.append(temp_array)
    unfor_leftmeans_7 = np.asarray(unfor_leftmeans_7)
    
    leftmeans_7 = []
    
    for i in range(len(output_arr_7)):
        k = len(unfor_leftmeans_7[i])
        new_list = []
        for j in range(k//3):
            new_list.append(unfor_leftmeans_7[i][3*j:3*(j + 1)])
        leftmeans_7.append(np.asarray(new_list))
    leftmeans_7 = np.asarray(leftmeans_7)
    
    means_7 = []
    for i in range(len(output_arr_7)):
        attr_arr = []
        for j in range(len(TLmeans_7[i])):
            meansarr = []
            meansarr.append(TLmeans_7[i][j][0])
            meansarr.append(leftmeans_7[i][j][0])
            meansarr.append(TLmeans_7[i][j][1] - leftmeans_7[i][j][0])
            meansarr.append(leftmeans_7[i][j][1])                        
            meansarr.append(TLmeans_7[i][j][2] - leftmeans_7[i][j][1])
            meansarr.append(leftmeans_7[i][j][2])
            meansarr.append(TLmeans_7[i][j][3] - leftmeans_7[i][j][2])
            attr_arr.append(np.asarray(meansarr))
        means_7.append(np.asarray(attr_arr))
    means_7 = np.asarray(means_7)
    
    
    # cvTL at different attractors
    unfor_cvTL_7 = []
    for i in range(len(output_arr_7)):
        temp_array = np.array(output_arr_7[5][i].replace('[', '').replace(']', '')
                        .replace(' ', '').split(',')).astype(float)
        unfor_cvTL_7.append(temp_array)
    unfor_cvTL_7 = np.asarray(unfor_cvTL_7)
    
    cvTLs_7 = []
    
    for i in range(len(output_arr_7)):
        k = len(unfor_cvTL_7[i])
        new_list = []
        for j in range(k//4):
            new_list.append(unfor_cvTL_7[i][4*j:4*(j + 1)])
        cvTLs_7.append(np.asarray(new_list))
    cvTLs_7 = np.asarray(cvTLs_7)
    
    unfor_leftCVs_7 = []
    for i in range(len(output_arr_7)):
        temp_array = np.array(output_arr_7[6][i].replace('[', '').replace(']', '')
                        .replace(' ', '').split(',')).astype(float)
        unfor_leftCVs_7.append(temp_array)
    unfor_leftCVs_7 = np.asarray(unfor_leftCVs_7)
    
    leftCVs_7 = []
    
    for i in range(len(output_arr_7)):
        k = len(unfor_leftCVs_7[i])
        new_list = []
        for j in range(k//3):
            new_list.append(unfor_leftCVs_7[i][3*j:3*(j + 1)])
        leftCVs_7.append(np.asarray(new_list))
    leftCVs_7 = np.asarray(leftCVs_7)
    
    unfor_rightCVs_7 = []
    for i in range(len(output_arr_7)):
        temp_array = np.array(output_arr_7[7][i].replace('[', '').replace(']', '')
                        .replace(' ', '').split(',')).astype(float)
        unfor_rightCVs_7.append(temp_array)
    unfor_rightCVs_7 = np.asarray(unfor_rightCVs_7)
    
    rightCVs_7 = []
    
    for i in range(len(output_arr_7)):
        k = len(unfor_rightCVs_7[i])
        new_list = []
        for j in range(k//3):
            new_list.append(unfor_rightCVs_7[i][3*j:3*(j + 1)])
        rightCVs_7.append(np.asarray(new_list))
    rightCVs_7 = np.asarray(rightCVs_7)
    
    CVs_7 = []
    for i in range(len(output_arr_7)):
        attr_arr = []
        for j in range(len(cvTLs_7[i])):
            CVarr = []
            CVarr.append(cvTLs_7[i][j][0])
            CVarr.append(leftCVs_7[i][j][0])
            CVarr.append(rightCVs_7[i][j][0])
            CVarr.append(leftCVs_7[i][j][1])
            CVarr.append(rightCVs_7[i][j][1])
            CVarr.append(leftCVs_7[i][j][2])
            CVarr.append(rightCVs_7[i][j][2])
            attr_arr.append(np.asarray(CVarr))
        CVs_7.append(np.asarray(attr_arr))
    CVs_7 = np.asarray(CVs_7)
    
    # relative frequencies of attractors
    freqs_7 = []
    for i in range(len(output_arr_7)):
        freqs_7.append(np.array(output_arr_7[8][i].replace('[', '').replace(']', '')
                                .replace(' ', '').split(',')).astype(float))
    freqs_7 = np.asarray(freqs_7)
    
    return [p_d_list_0, p_d_list_1, p_d_list_2, p_d_list_3, p_d_list_4,
            p_d_list_5, p_d_list_6, p_d_list_7, ext_0, ext_1, ext_2, ext_3,
            ext_4, ext_5, ext_6, ext_7, TLmeans_0, TLmeans_1, TLmeans_2, 
            TLmeans_3, TLmeans_4, TLmeans_5, TLmeans_6, TLmeans_7, cvTLs_0, 
            cvTLs_1, cvTLs_2, cvTLs_3, cvTLs_4, cvTLs_5, cvTLs_6, cvTLs_7, 
            means_0, means_1, means_2, means_3, means_4, means_5, means_6, 
            means_7]
```
