## Supplementary material for "Top predators govern multitrophic diversity effects in tritrophic food webs": Code and data: table_3.py

```
#!/usr/bin/env python3
# -*- coding: utf-8 -*-
"""
Created on Fri Jun 5 2020

@author: ruben
"""

# this script will print the OOB values of all quantities in Table 3 (main text)


import pandas as pd
import numpy as np
import ast
import matplotlib.pyplot as plt

import time

from sklearn.ensemble import ExtraTreesRegressor

from import_raw_data import import_data
from calc_prods import calc_prods

# %%===========================================================================
# import data
# =============================================================================
p_d_list = [[], [], [], [], [], [], [], []]
ext = [[], [], [], [], [], [], [], []]
TLmeans = [[], [], [], [], [], [], [], []]
cvTLs = [[], [], [], [], [], [], [], []]
means = [[], [], [], [], [], [], [], []]

quantity_names_B = ['N', 'B', 'I', 'T']
quantity_names_CV = ['Ncv', 'Bcv', 'Icv', 'Tcv']
quantity_names_prod = ['PT PB', 'Bup', 'PB B', 'PT']


[p_d_list[0], p_d_list[1], p_d_list[2], p_d_list[3], p_d_list[4],
p_d_list[5], p_d_list[6], p_d_list[7], ext[0], ext[1], ext[2], ext[3],
ext[4], ext[5], ext[6], ext[7], TLmeans[0], TLmeans[1], TLmeans[2], 
TLmeans[3], TLmeans[4], TLmeans[5], TLmeans[6], TLmeans[7], cvTLs[0], 
cvTLs[1], cvTLs[2], cvTLs[3], cvTLs[4], cvTLs[5], cvTLs[6], cvTLs[7], 
means[0], means[1], means[2], means[3], means[4], means[5], means[6], 
means[7]] = import_data()

# put production metrics (figure 5) in same structure as the others
prods = []
for web in range(8):
    web_prods = []
    for combination in range(len(p_d_list[web])):
        prods_attr = []
        for attr in range(len(means[web][combination])):
            prod = calc_prods(means[web][combination][attr], 
                               p_d_list[web][combination], web)
            prods_attr.append(prod)
        web_prods.append(prods_attr)
    prods.append(web_prods)

# make sure that all trait differences are added to every data point
def add_all_tdiffs_everywhere(web):
    if web == 0:
        add_dict = {'B_diff': 0, 'I_diff': 0, 'T_diff': 0}
    if web == 1:
        add_dict = {'I_diff': 0, 'T_diff': 0}
    if web == 2:
        add_dict = {'B_diff': 0, 'T_diff': 0}
    if web == 3:
        add_dict = {'T_diff': 0}
    if web == 4:
        add_dict = {'B_diff': 0, 'I_diff': 0}
    if web == 5:
        add_dict = {'I_diff': 0}
    if web == 6:
        add_dict = {'B_diff': 0}
    if web == 7:
        return
    for i in p_d_list[web]:
            i.update(add_dict)
    return
    
for web in range(8):
    add_all_tdiffs_everywhere(web)

feature_labels = ['B_diff', 'I_diff', 'T_diff', 'N0', 'h_N', 'r0', 'a0', 'h0', 
                'alpha0', 'eta0', 'delta', 'n', 'nu', 'a_scale']
    
# %%===========================================================================
# "main"
# =============================================================================

num_cores = 4
n_trees = 2000

biomasses
for level in [0, 1, 2, 3]:
    # first prepare data for random forest algorithm
    function = []
    data_samples = []
    for web in [0, 1, 2, 3, 6, 7]: # do not treat webs 4 and 5
        for i in range(len(p_d_list[web])):
            if 0 in ext[web][i]:
                indices = np.where(ext[web][i] == 0)
                for j in indices[0]:
                    function.append(TLmeans[web][i][j][level])
                    values = []
                    for key in feature_labels:
                        values.append(p_d_list[web][i][key])
                    data_samples.append(values)
                        
    function = np.asarray(function)
    data_samples = np.asarray(data_samples)

    # train random forest
    ts = time.time()
    clf = ExtraTreesRegressor(n_estimators = n_trees, bootstrap = 1, oob_score = 1, n_jobs = num_cores)
    model = clf.fit(data_samples, function)
    print("OOB " + quantity_names_B[level] + ': ' + str(model.oob_score_))

# CVs
for level in [0, 1, 2, 3]:
    # first prepare data for random forest algorithm
    function = []
    data_samples = []
    for web in range(8):
        for i in range(len(p_d_list[web])):
            if 0 in ext[web][i]:
                indices = np.where(ext[web][i] == 0)
                for j in indices[0]:
                    function.append(cvTLs[web][i][j][level])
                    values = []
                    for key in feature_labels:
                        values.append(p_d_list[web][i][key])
                    data_samples.append(values)
                        
    function = np.asarray(function)
    data_samples = np.asarray(data_samples)

    # train random forest
    ts = time.time()
    clf = ExtraTreesRegressor(n_estimators = n_trees, bootstrap = 1, oob_score = 1, n_jobs = num_cores)
    model = clf.fit(data_samples, function)
    print("OOB " + quantity_names_CV[level] + ': ' + str(model.oob_score_))

# production metrics
for level in [0, 1, 2, 3]:
    # first prepare data for random forest algorithm
    function = []
    data_samples = []
    for web in range(8):
        for i in range(len(p_d_list[web])):
            if 0 in ext[web][i]:
                indices = np.where(ext[web][i] == 0)
                for j in indices[0]:
                    function.append(prods[web][i][j][level])
                    values = []
                    for key in feature_labels:
                        values.append(p_d_list[web][i][key])
                    data_samples.append(values)
                        
    function = np.asarray(function)
    data_samples = np.asarray(data_samples)

    # train random forest
    ts = time.time()
    clf = ExtraTreesRegressor(n_estimators = n_trees, bootstrap = 1, oob_score = 1, n_jobs = num_cores)
    model = clf.fit(data_samples, function)
    print("OOB " + quantity_names_prod[level] + ': ' + str(model.oob_score_))
```
