## Supplementary material for "Top predators govern multitrophic diversity effects in tritrophic food webs": Code and data: sys_solve.c

```
#include "sys_solve.h"

//***********************************************************
// Holling Type III ** with bidirectional trait axis **
//***********************************************************

//  basic simulation variables:
int S = 6;                      // number of species
int N = 1;                      // number of nutrients
int SN = 7;                      // number of species and nutrients
int N_A = 2;					// number of Basal species
int N_I = 2;					// number of Int species
int N_T = 2;					// number of Top

const double eps_rel= 1e-10;       // absolute error tolerance for ODE solver
const double eps_abs = 1e-10;       // relative error tolerance for ODE solver
const double Delta_t = 0.01;             // step size in ODE integration (internal step size can be less or larger) 
const double EXTINCT = 1e-10;       // extinction threshold for biomass densities


char timeseries_file[99];    // name of output file for the time series
char mean_file[99];    // name of output file for mean and Cv of biomasses

// params from input
double tend;                 		// simulation time

double N0;				// Nutrient inflow concentration
double h_N;				// Nutrient uptake half saturation constant

double r0;				//growth rate for Delta_B = 0
double r_inc;			//increase in growth rate (proportion of r0) due to B TO

double h0;				//B-I handling time for Delta_B = Delta_I = 0
double h_dec;			//decrease in handling time rate (proportion of h0) due to I TO

double eta0;			//I-T handling time for Delta_I = Delta_T = 0
double eta_dec;			//decrease in handling time rate (proportion of eta0) due to T TO

double a_s;				//B-I attack rate for Delta_B = Delta_I = 0 (straight links)
double alpha_s;			//I-I attack rate for Delta_I = Delta_B = 0 (straight links)

double a_factor;		//cross link reduction factor

double delta;			// nutrient dilution rate

double e_i;				// fast chain efficiency
double e_t;				// slow chain efficiency	

double n;				// B-I Hill coeff.
double nu;				// I-T Hill coeff.
double ratio;			// N to C ratio 
double teval;           // time for evaluation of mean and coeff. of variation

int TIMESERIES;            // if true, for the last network a file with the time series will be created  
int POP_STATS;            // 1: mean biomass densities and coefficients of variance are calculated
int para_id;
int web_id;

double B_diff;
double I_diff;
double T_diff;

// params which are a function of Delta
	// 1 = fast chain, 2 = slow chain
double a_x;
double alpha_x;
double* r_arr;
double* h_arr;
double* eta_arr;

	// memory for temp vectors
double* Atemp;
double* Itemp;
double* Ttemp;
double* a_arr;
double* alpha_arr;

double* deltaA_arr;
double* deltaI_arr;
double* deltaT_arr;

double* Mtemp_arr;
double* g_arr;
double* gamma_arr;

//*******************************************************************
// Main Part
//*******************************************************************
int main(int argc, char *argv[]) {
	if(argc != 23 + 1 + S + N + 3){printf("Needs %d arguments\n", 23 + S + N + 3);return(0);}
	else{
		FILE *data;	
		
		//Write_header(data);                        // write header for file with processed output
		
		Allocate_global_array_memory();
		set_args(argv);   
		
		Initialise_model(argv);              // in this function all the magic happens!
		
		Free_global_array_memory();
		printf("succes");
		return(0);
	}
}

static void set_args(char *argv[]){
	
	tend = atof(argv[1]);
	N0 = atof(argv[2]);
	h_N = atof(argv[3]);
	r0 = atof(argv[4]);
	r_inc = atof(argv[5]);
	h0 = atof(argv[6]);
	h_dec = atof(argv[7]);
	eta0 = atof(argv[8]);
	eta_dec = atof(argv[9]);
	a_s = atof(argv[10]);
	alpha_s = atof(argv[11]);
	a_factor = atof(argv[12]);
	delta = atof(argv[13]);
	e_i = atof(argv[14]);
	e_t = atof(argv[15]);
	n = atof(argv[16]);
	nu = atof(argv[17]);
	ratio  = atof(argv[18]);
	teval = atof(argv[19]);
	TIMESERIES = atof(argv[20]);
	POP_STATS  = atof(argv[21]);
	para_id = atof(argv[22]);
	web_id = atoi(argv[23]);

	B_diff = atof(argv[23 + S + N + 1]);
	I_diff = atof(argv[23 + S + N + 2]);
	T_diff = atof(argv[23 + S + N + 3]);

} 

static void print_gsl_vector(gsl_vector *V, int length){
int i;
	for (i = 0; i < length; i++){
		printf("%d = %g \n", i, gsl_vector_get(V, i));
	}
}

static void print_gsl_matrix(gsl_matrix *M, int n_rows, int n_cols){
int i, j;
	for (i = 0; i < n_rows; i++){
		for (j = 0; j < n_cols; j++){
			if (j == 0){
				printf("|  ");
			}
			printf("%g  ", gsl_matrix_get(M, i, j));
		}
	}
	printf("|");
}

static void gsl_vector_pow(gsl_vector *V, int length, double exponent){
	
	int i;
	
	for (i = 0; i < length; i++){
		
		gsl_vector_set(V, i, pow(gsl_vector_get(V, i), exponent));
	}
}

double min(double a, double b){
	return (a < b ? a : b);
}

//***************************************************************************************
// call functions for network creation, parameters, calculation of dynamics, and output
//*************************************************************************************** 
static void Initialise_model(char *argv[]) {
  
  double *B = (double *) calloc(S + N, sizeof(double));                      // biomass of the species
  double *meanB = (double *) calloc(S + N, sizeof(double));                  // mean TL biomass during evaluation period
  double *CV = (double *) calloc(S + N, sizeof(double));                   // coefficient of variation of biomasses
  
  double* meanB_TL = (double *) calloc(3, sizeof(double));
  double* CV_TL = (double *) calloc(3, sizeof(double));

//  *********** calling the most important functions that define and solve the problem ***********
  Generate_network_structure();                             // produce network topology
  Initialise_biomass_densities(B, argv);                     // set the initial conditions (initial biomass densities)
  
  Solve_ODE(B, meanB, CV, meanB_TL, CV_TL);                  // integrate the system of ODEs
  Write_mean_output(meanB, CV, meanB_TL, CV_TL);             // write mean and CV of biomasses to file
//  *********** Free memories ***********
  free(B);
  free(meanB);
  free(CV);
  free(meanB_TL);
  free(CV_TL);
}

//***************************************************************************************
//  basic network topology of an allometric feeding kernel-food web
//***************************************************************************************
static void Generate_network_structure() {
	int i;

	double B_inc = 1 + B_diff*r_inc;
	double I_inc = 1 + I_diff*h_dec;
	double T_inc = 1 + T_diff*eta_dec;

	*(r_arr) = r0*B_inc;
	*(r_arr + 1) = r0/B_inc;
	
	*(h_arr) = h0/(I_inc*B_inc);
	*(h_arr + 1) = h0*B_inc/I_inc;
	*(h_arr + 2) = h0*I_inc/B_inc;
	*(h_arr + 3) = h0*B_inc*I_inc;

	*(eta_arr) = eta0/(I_inc*T_inc);
	*(eta_arr + 1) = eta0*I_inc/T_inc;
	*(eta_arr + 2) = eta0*T_inc/I_inc;
	*(eta_arr + 3) = eta0*I_inc*T_inc;

	double att_rat_arr[5];
	for(i = 0; i < 5; i++){
		double exp = (double) i;
		att_rat_arr[i] = pow(a_factor, -exp);
	} 
	int a_index = 4*min(B_diff, I_diff);
	int alpha_index = 4*min(I_diff, T_diff);

	a_x = att_rat_arr[a_index];
	alpha_x = att_rat_arr[alpha_index];

	*(a_arr) = a_s;
	*(a_arr + 1) = a_x*a_s;
	*(a_arr + 2) = a_x*a_s;
	*(a_arr + 3) = a_s;

	*(alpha_arr) = alpha_s;
	*(alpha_arr + 1) = alpha_x*alpha_s;
	*(alpha_arr + 2) = alpha_x*alpha_s;
	*(alpha_arr + 3) = alpha_s;

    *(deltaA_arr) = -delta*B_inc;
    *(deltaA_arr + 1) = -delta/B_inc;
    *(deltaI_arr) = -delta*I_inc;
    *(deltaI_arr + 1) = -delta/I_inc;
    *(deltaT_arr) = -delta*T_inc;
    *(deltaT_arr + 1) = -delta/T_inc;

	// to get the right file to output to

	char ch_id[10];
	sprintf(ch_id, "%d-%d", para_id, web_id);
	
	strcpy(timeseries_file, "../../c_output_data/timeseries/timeseries");    // name of output file for the time series
	strcpy(mean_file, "../../c_output_data/mean/mean");    // name of output file for mean and Cv of biomasses
	strcat(timeseries_file, ch_id);
	strcat(timeseries_file, ".out");
	
	strcat(mean_file, ch_id);
	strcat(mean_file, ".out");
}

	
//***************************************************************************************
//  set initial biomass densities
//***************************************************************************************
static void Initialise_biomass_densities(double B[], char *argv[]) {
  int i;
  
  for(i = 0; i < S + N; i++){
	B[i] = atof(argv[24 + i]);
  }
  return;
}


//***************************************************************************************
//  efficient environment for solving the ODE
//***************************************************************************************
static void Solve_ODE(double B[], double meanB[], double CV[], double meanB_TL[], double CV_TL[])
{
	int i, flag, EXT_FLAG;
	double t = 0, tout = 0.01, t_stop;                               // time; time for next output; time at which the solver stopped

	gsl_vector_view Bvec = gsl_vector_view_array(B, S + N);         // biomass densities (including resources)
	gsl_vector_view Mvec = gsl_vector_view_array(meanB, S + N);     // mean biomass denisties (including resources)

	FILE *timeseries;
	if(TIMESERIES)
		Prepare_timeseries_file(timeseries);
	N_Vector y = N_VNew_Serial(S+N);							// N_vector is a sundials vector object, 
																// N_VNew_Serial creates  and  allocates  memory  for  a  serial N_Vector.   
																// 	  Its  only  argument  is  the vector length.
	N_VSetArrayPointer(B, y);  								// overwrites data in N_Vector with a given array of realtype
  
	void *cvode_mem = CVodeCreate(CV_BDF, CV_NEWTON); 		// set up solver; BDF = backward diff formula
	CVodeInit(cvode_mem, Cvode_rates, t, y);					// initialize solver; Cvode_rates used by solver apparently
	CVodeSStolerances(cvode_mem, eps_rel, eps_abs);			// set tolerances
	CVodeSetMaxHnilWarns(cvode_mem, -1);					// no warnings
	CVDense(cvode_mem, S+N);									// LU factorization and backsolving with dense matrices
	
 
  
// ********* integrate ODE to reach an attractor; optional: output of time series ********
	while (t < tend) {
		flag = CVode(cvode_mem, tout, y, &t, CV_ONE_STEP);
	
		if (flag != CV_SUCCESS)
			break;
	
		if (TIMESERIES)
			Write_timeseries_to_file(B,t,timeseries);

		EXT_FLAG = 0;  
	    for (i = 0; i < SN; i++) {
	      if (B[i] < EXTINCT && B[i] != 0)                     // determine whether a species just fell below the extinction threshold
	        EXT_FLAG = 1;
	    }
	    if (EXT_FLAG){                                         // if an extinction occured:
		  Extinction(B, SN);                                     // set the biomass of the species to 0...
	      tout = t + 0.01;                                       // set the next output time to a reasonable value...
	      CVodeInit(cvode_mem, Cvode_rates, t, y);             // and re-initialise the solver
	      CVDense(cvode_mem, SN);
	    }
	}
  
// ********* integrate ODE to calculate mean biomass densities and coefficients of variation ********
	if(POP_STATS) {
    	t_stop = t;
    	tout = t + Delta_t;
    	while (t < t_stop + 2*teval) {
			flag = CVode(cvode_mem, tout, y, &t, CV_NORMAL);

			if(flag != CV_SUCCESS)
			break;
			else
				tout += Delta_t;

    		EXT_FLAG = 0;  
		    for (i = 0; i < SN; i++) {
		      if (B[i] < EXTINCT && B[i] != 0)                     // determine whether a species just fell below the extinction threshold
		        EXT_FLAG = 1;
		    }
			if (t < t_stop + teval)
				gsl_vector_add(&Mvec.vector, &Bvec.vector);

			if (fabs(t - (t_stop + teval)) < 0.1*Delta_t)
				gsl_vector_scale(&Mvec.vector, Delta_t/teval);

			if (t >= t_stop + teval) {
				for (i = 0; i < S+N; i++)
			  		CV[i] += (B[i] - meanB[i])*(B[i] - meanB[i]);


				for (i = 0; i < 3; i++){
			  		CV_TL[i] += (B[2*i + 1] + B[2*(i + 1)] - meanB[2*i + 1] - meanB[2*(i + 1)])
			  				  *(B[2*i + 1] + B[2*(i + 1)] - meanB[2*i + 1] - meanB[2*(i + 1)]);
				}
			}

			if (EXT_FLAG){                                         // if an extinction occured:
				Extinction(B, SN);                                     // set the biomass of the species to 0...
				tout = t + 0.01;                                       // set the next output time to a reasonable value...
				CVodeInit(cvode_mem, Cvode_rates, t, y);             // and re-initialise the solver
				CVDense(cvode_mem, SN);
		    }

    	}  
     // MULTIPLY CV by Delta_t/teval

    	for(i = 0; i < 3; i++) {
    		meanB_TL[i] = meanB[2*i + 1] + meanB[2*(i + 1)];
    	}

	    for (i=0; i < SN; i++) {
	      if (meanB[i] > EXTINCT)
	        CV[i] = sqrt(CV[i]*Delta_t/teval);                          // Coefficient of variation 
	      else
	        CV[i] = 0;
	    }


	    for(i = 0; i < 3; i++) {
			if (meanB_TL[i] > EXTINCT){
				CV_TL[i] = sqrt(CV_TL[i]*Delta_t/teval);  // standard deviation
			}
			else {CV_TL[i] = 0;}
	    }
	     
	}
  
	CVodeFree(&cvode_mem);    

	return;
}

//***************************************************************************************
//  set biomass densities below the extinction threshold to 0
//***************************************************************************************
static void Extinction(double B[], int length) {
  int i;
  for (i = 0; i < length; i++) {
    if (B[i] < EXTINCT)
      B[i] = 0;
  }
      
  return;
}

//***************************************************************************************
//  function that is called by the cvode-solver
//***************************************************************************************
static int Cvode_rates(double t, N_Vector y, N_Vector ydot, void *params) {  
  Calculate_total_rates(N_VGetArrayPointer(y), N_VGetArrayPointer(ydot));
  return(0);
}

//***************************************************************************************
//  define elements of Bdot[], i.e., the differential equations
//***************************************************************************************
static void Calculate_total_rates(double B[], double Bdot[]) {
	int i;

	gsl_vector_view B_vec = gsl_vector_view_array(B, S+N);
	gsl_vector *Bvec = &B_vec.vector;
  
	gsl_vector_view Bdot_vec = gsl_vector_view_array(Bdot,S+N);          // vector for time derivatives
	gsl_vector *Bdotvec = &Bdot_vec.vector;
	Calculate_net_rates(Bvec,Bdotvec);                      // calculate dilution (Dvec) and transfer (Tvec) rates
  
	for (i=0; i<S+N; i++){
		Bdot[i] = gsl_vector_get(Bdotvec, i);
	}
   
	return;
}  

//***************************************************************************************
//  calculate net per capita rates of biomass production and mortality 
//***************************************************************************************
static void Calculate_net_rates(gsl_vector *Bvec, gsl_vector *Bdotvec) {
	int i;
	// make matrix/vector views

	gsl_vector_view r_v = gsl_vector_view_array(r_arr, N_A);
	gsl_vector* r = &r_v.vector;
	
	gsl_matrix_view h_v = gsl_matrix_view_array(h_arr, N_I, N_A); // gsl_matrix_view_arr(arr, Nrows, Ncols):
	gsl_matrix *h = &h_v.matrix;				      // matrix[i, j] = arr[i * Ncols + j]
	gsl_matrix_view eta_v = gsl_matrix_view_array(eta_arr, N_T, N_I);
	gsl_matrix *eta = &eta_v.matrix;

	// make vectors for N, B, I, T
	double N = gsl_vector_get(Bvec, 0);

	gsl_vector_view A_v = gsl_vector_subvector(Bvec, 1, N_A);
	gsl_vector* A = &A_v.vector;
	
	gsl_vector_view I_v = gsl_vector_subvector(Bvec, 1 + N_A, N_I);
	gsl_vector* I = &I_v.vector;

	gsl_vector_view T_v = gsl_vector_subvector(Bvec, 1 + N_A + N_I, N_T);
	gsl_vector* T = &T_v.vector;

	// make some temporary vectors for calculations

	gsl_vector_view A_t = gsl_vector_view_array(Atemp, N_A);
	gsl_vector* A_temp = &A_t.vector;
	
	gsl_vector_view I_t = gsl_vector_view_array(Itemp, N_I);
	gsl_vector* I_temp = &I_t.vector;

	gsl_vector_view T_t = gsl_vector_view_array(Ttemp, N_T);
	gsl_vector* T_temp = &T_t.vector;

	gsl_matrix_view a_v = gsl_matrix_view_array(a_arr, N_I, N_A);
	gsl_matrix *a = &a_v.matrix;
	gsl_matrix_view alpha_v = gsl_matrix_view_array(alpha_arr, N_I, N_A);
	gsl_matrix *alpha = &alpha_v.matrix;

	gsl_matrix_view Mtemp_v = gsl_matrix_view_array(Mtemp_arr, 2, 2);
	gsl_matrix *Mtemp = &Mtemp_v.matrix;

	gsl_matrix_view g_v = gsl_matrix_view_array(g_arr, N_I, N_A);
	gsl_matrix *g = &g_v.matrix;
	gsl_matrix_view gamma_v = gsl_matrix_view_array(gamma_arr, N_T, N_I);
	gsl_matrix *gamma = &gamma_v.matrix;

	gsl_vector_view delta_A_v = gsl_vector_view_array(deltaA_arr, N_A);
	gsl_vector* delta_A = &delta_A_v.vector;
	gsl_vector_view delta_I_v = gsl_vector_view_array(deltaI_arr, N_I);
	gsl_vector* delta_I = &delta_I_v.vector;
	gsl_vector_view delta_T_v = gsl_vector_view_array(deltaT_arr, N_T);
	gsl_vector* delta_T = &delta_T_v.vector;


	// Bdotvec subvectors
	gsl_vector_view Adot = gsl_vector_subvector(Bdotvec, 1, N_A);
	gsl_vector_view Idot = gsl_vector_subvector(Bdotvec, 1 + N_A, N_I);
	gsl_vector_view Tdot = gsl_vector_subvector(Bdotvec, 1 + N_A + N_I, N_T);


	gsl_vector_set_zero(Bdotvec);

	// Nutrients
	double N_uptake = N/(N + h_N);

	gsl_vector_memcpy(A_temp, A);
	gsl_vector_scale(A_temp, N_uptake);


	gsl_vector_mul(A_temp, r);
	double feedtot = gsl_blas_dasum(A_temp);
	feedtot *= ratio;

	gsl_vector_set(Bdotvec, 0, -feedtot + delta*(N0 - N)); // - sum r_i A_i + dilution for nutrients
	
	gsl_vector_add(&Adot.vector, A_temp);  // r_i*A_i for a species
	// Autotrophs

	// reset A_temp
	gsl_vector_memcpy(A_temp, A);

	gsl_vector_pow(A_temp, N_A, n);

	// Mtemp = element-wise multiplication of a and h
	gsl_matrix_memcpy(Mtemp, h);
	gsl_matrix_mul_elements(Mtemp, a);

	// I_temp = vector of total food of I's
	gsl_blas_dgemv(CblasNoTrans, 1, Mtemp, A_temp, 0, I_temp);
	
	gsl_vector_add_constant(I_temp, 1);

	gsl_vector_pow(I_temp, N_I, -1);

	// ah -> temp grazing matrix
	gsl_matrix_memcpy(g, a);

	for(i = 0; i < N_A; i++){
		gsl_vector_view col = gsl_matrix_column(g, i);
		gsl_vector_mul(&col.vector, I_temp);
	}

	for(i = 0; i < N_I; i++){
		gsl_vector_view row = gsl_matrix_row(g, i);
		gsl_vector_mul(&row.vector, A_temp);
		feedtot = gsl_blas_dasum(&row.vector);
		gsl_vector_set(&Idot.vector, i, e_i*feedtot); // e sum_i g_ji I_j to I
	}
	gsl_vector_mul(&Idot.vector, I);

	// sum_j g_ji I_j
	gsl_blas_dgemv(CblasTrans, 1, g, I, 0, A_temp);

	gsl_vector_scale(A_temp, -1);
	gsl_vector_add(&Adot.vector, A_temp);		 // sum_j g_ji I_j

	// Intermediates
	gsl_vector_memcpy(I_temp, I); 

	gsl_vector_pow(I_temp, N_I, nu);

	// alphaetamatr = element-wise multiplication of a and h
	gsl_matrix_memcpy(Mtemp, eta);
	gsl_matrix_mul_elements(Mtemp, alpha);

	// T_temp = vector of total food of I's
	gsl_blas_dgemv(CblasNoTrans, 1, Mtemp, I_temp, 0, T_temp);

	gsl_vector_add_constant(T_temp, 1);

	gsl_vector_pow(T_temp, N_T, -1);

	// alphaeta -> temp grazing matrix
	gsl_matrix_memcpy(gamma, alpha);

	for(i = 0; i < N_I; i++){
		gsl_vector_view col = gsl_matrix_column(gamma, i);
		gsl_vector_mul(&col.vector, T_temp);
	}

	for(i = 0; i < N_T; i++){
		gsl_vector_view row = gsl_matrix_row(gamma, i);
		gsl_vector_mul(&row.vector, I_temp);
		feedtot = gsl_blas_dasum(&row.vector);
		gsl_vector_set(&Tdot.vector, i, e_t*feedtot); // e sum_i gamma_ji T_j to T
	}
	gsl_vector_mul(&Tdot.vector, T);

	// sum_j gamma_ji T_j
	gsl_blas_dgemv(CblasTrans, 1, gamma, T, 0, I_temp);

	gsl_vector_scale(I_temp, -1);
	gsl_vector_add(&Idot.vector, I_temp);		 // sum_j g_ji I_j

	// add dilution terms for species
	gsl_vector_memcpy(A_temp, A);
	gsl_vector_memcpy(I_temp, I);
	gsl_vector_memcpy(T_temp, T);

	gsl_vector_mul(A_temp, delta_A);
	gsl_vector_mul(I_temp, delta_I);
	gsl_vector_mul(T_temp, delta_T);

	gsl_vector_add(&Adot.vector, A_temp);
	gsl_vector_add(&Idot.vector, I_temp);
	gsl_vector_add(&Tdot.vector, T_temp);

  return;
}

  

//***************************************************************************************
//  write mean and CV of biomasses to an output file
//***************************************************************************************

static void Write_mean_output(double meanB[], double CV[], double meanB_TL[], double CV_TL[]) 
{
	int i;

	FILE *file1;
	file1 = fopen(mean_file,"w");

	for (i = 0; i < S + N; i++){
		if(i < 3){
		fprintf(file1,"%.9g	%.9g %.9g\n", meanB[i], CV[i], CV_TL[i]);
		}
		else{fprintf(file1,"%.9g %.9g\n",meanB[i], CV[i]);}
	}

	fclose(file1);

	return;
}
//***************************************************************************************
//  write the time series of the last run to an output file
//***************************************************************************************
static void Prepare_timeseries_file(FILE *timeseries) {
  timeseries = fopen(timeseries_file,"w");
  fclose(timeseries);
  
  return;
}

static void Write_timeseries_to_file(double B[], double t, FILE *timeseries) {
  int i;
  timeseries = fopen(timeseries_file,"a");
  
  fprintf(timeseries,"%.8g  ",t);
  for(i=0; i < S+N; i++)
    fprintf(timeseries,"%.8g  ", B[i]);
  fprintf(timeseries,"\n");
  
  fclose(timeseries);
  
  return;
}

//***************************************************************************************
//  allocate and free the memory for the globally defined arrays used by Calculate_net_rates
//***************************************************************************************
static void Allocate_global_array_memory() {
	
	r_arr = (double *) calloc(N_A, sizeof(double));

	a_arr = (double *) calloc(N_I*N_A, sizeof(double));
	alpha_arr = (double *) calloc(N_T*N_I, sizeof(double));

	h_arr = (double *) calloc(N_I*N_A, sizeof(double));
	eta_arr = (double *) calloc(N_T*N_I, sizeof(double));
	
	Atemp = (double *) calloc(N_A, sizeof(double));
	Itemp = (double *) calloc(N_I, sizeof(double));
	Ttemp = (double *) calloc(N_T, sizeof(double));

	Mtemp_arr = (double *) calloc(4, sizeof(double));
	g_arr = (double *) calloc(N_A*N_I, sizeof(double));
	gamma_arr = (double *) calloc(N_I*N_T, sizeof(double));

	deltaA_arr = (double *) calloc(N_A, sizeof(double));
	deltaI_arr = (double *) calloc(N_I, sizeof(double));
	deltaT_arr = (double *) calloc(N_T, sizeof(double));

    return;
}

static void Free_global_array_memory(){

	free(r_arr);
	free(a_arr);
	free(alpha_arr);
	free(h_arr);
	free(eta_arr);
	free(Atemp);
	free(Itemp);
	free(Ttemp);
	free(Mtemp_arr);
	free(g_arr);
	free(gamma_arr);
	free(deltaA_arr);
	free(deltaI_arr);
	free(deltaT_arr);

	return;
} 
```
