## Supplementary material for "Top predators govern multitrophic diversity effects in tritrophic food webs": Code and data: sys_solve.h

```
#include <stdio.h>
#include <math.h>
#include <time.h>
#include <string.h>
#include <gsl/gsl_matrix.h>
#include <gsl/gsl_vector.h>
#include <gsl/gsl_rng.h>            // random number generator
#include <gsl/gsl_blas.h>           // linear algebra with vecors and matrices
#include <gsl/gsl_randist.h>        // random number distributions
#include <gsl/gsl_sort_vector.h>    // vector operations
#include <cvode/cvode.h>            /* prototypes for CVODE fcts., consts. */
#include <nvector/nvector_serial.h> /* serial N_Vector types, fcts., macros */
#include <cvode/cvode_dense.h>      /* prototype for CVDense */

static void set_args(char *argv[]);
static void print_gsl_vector(gsl_vector *V, int length);
static void print_gsl_matrix(gsl_matrix *M, int n_rows, int n_cols);
static void gsl_vector_pow(gsl_vector *V, int length, double exponent);
double min(double a, double b);
static void Initialise_model(char *argv[]);
static void Generate_network_structure();
static void Initialise_biomass_densities(double B[], char *argv[]);
static void Solve_ODE(double B[], double meanB[], double CV[], double meanB_TL[], double CV_TL[]);
static void Extinction(double B[], int length);
static int Cvode_rates(double t, N_Vector y, N_Vector ydot, void *params);
static void Calculate_total_rates(double B[], double Bdot[]);
static void Calculate_net_rates(gsl_vector *Bvec, gsl_vector *Bdotvec);
static void Write_mean_output(double meanB[], double CV[], double meanB_TL[], double CV_TL[]);
static void Prepare_timeseries_file(FILE *timeseries);
static void Write_timeseries_to_file(double B[], double t, FILE *timeseries);
static void Allocate_global_array_memory();
static void Free_global_array_memory();
```
