## Supplementary material for "Top predators govern multitrophic diversity effects in tritrophic food webs": Code and data: analyze_web.py

```
#!/usr/bin/env python3
# -*- coding: utf-8 -*-
"""
Created on Fri Jun 5 2020

@author: ruben
"""
import numpy as np
import random as rd
import time
from joblib import Parallel, delayed # for parallelization
import multiprocessing
import pandas as pd   # for fast reading of files
import collections    # for the ordered dict
import subprocess     # for the execution of the C program
import os

# %%===========================================================================
# fixed params and loop properties etc
# =============================================================================
webid = 0 # which food web to check: 0: chain
                                    #1: B
                                    #2: I
                                    #3: BI
                                    #4: T
                                    #5: BT
                                    #6: IT
                                    #7: BIT


total = 1024  # total number of parameter combinations to try
duplicates = 200 # number of initial conditions to try per parameter combination

num_cores = 72  # number of cores to use for parallelization

# make sure to check first how long the simulation are expected to take. 
# "normally" the ODE solving in C takes a few seconds, but it can be up to one minute
# in exceptional cases. 

# the code will generate total*duplicates output files. this number can be 
# very large and make it hard to handle (even just deleting can take long)
# it set a maximal number make sure this number is much larger than duplicates*num_cores
# if all files should be kept and none overwritten set it to something larger 
# than total*duplicates 
max_output_files = 1e5 


# =============================================================================
# arrays for sampling random values inbetween
#                [begin,   end]
N0_a = np.array([1120/2, 1120*2])
h_N_a = np.array([10/2, 10*2])
r_a = np.array([1/2, 1*2])
a_a = 1e-3*np.array([1.04/2, 1.04*2])
h_a = np.array([1.15/2, 1.15*2])
alpha_a = 1e-4*np.array([4.28/2, 4.48*2])
eta_a = np.array([2.62/2, 2.62*2])
a_scale_a = np.array([1, 500**(1/4)])
delta_a = 1e-2*np.array([3, 6])
n_a = np.array([1, 2])
nu_a = np.array([1, 2])

B_diff_a = np.linspace(0.25, 1, 4)
I_diff_a = np.linspace(0.25, 1, 4)
T_diff_a = np.linspace(0.25, 1, 4)

# "initialize" dictionary
c_params = collections.OrderedDict()

c_params['tend'] = 1e5 # relax to attractor
c_params['N0'] = 1120
c_params['h_N'] = 10
c_params['r0'] = 1
c_params['r_inc'] = 0.5
c_params['h0'] = 1.15
c_params['h_dec'] = 0.5
c_params['eta0'] = 2.62
c_params['eta_dec'] = 0.5
c_params['a0'] = 2.39e-3
c_params['alpha0'] = 8.48e-4
c_params['a_scale'] = 1

c_params['delta'] = 0.055
c_params['e_i'] = 0.33
c_params['e_t'] = 0.33
c_params['n'] = 1.10
c_params['nu'] = 1.10
c_params['ratio'] = 0.175

c_params['teval'] = 3e4 # eval mean
c_params['TIMESERIES'] = 0 # put to 1 if timeseries should be written to file
                           # !!!!! CAREFUL THIS MAY PRODUCE A HIGH NUMBER OF VERY LARGE FILES !!!!!
c_params['POP_STATS'] = 1
c_params['id'] = 0
c_params['web_id'] = webid

c_params['N_0'] = 10
c_params['B1_0'] = 10
c_params['B2_0'] = 10
c_params['I1_0'] = 10
c_params['I2_0'] = 10
c_params['T1_0'] = 10
c_params['T2_0'] = 10

c_params['B_diff'] = 0
c_params['I_diff'] = 0
c_params['T_diff'] = 0

# %%===========================================================================
# functions
# =============================================================================

# call the C code in parallel
def run_parallel():
    Parallel(n_jobs = num_cores)(delayed(test_param_combination)(i) for i in range(0, total))

# test a certain parameter combination multiple times and add the output to the data file
def test_param_combination(i):
    with open('../../python_output_data/' + str(int(webid)) + '-data' + '_' + str(int(total))
            + '-' + str(int(duplicates)) + '.out', 'a') as out_file:
        # get new parameter combination
        p = set_params(i, total)

        # unique id per simulation
        p['id'] = i*duplicates

        # get relevant information on properties such as mean biomasses, CVs, also number of attractors
        e_flag, num, mean, mean_1, cv, cv_1, cv_2, prop = calc_prop(p)

        # write data to Python output file for further analysis later
        # the parameters
        out_file.write(str(p) + "; ") 
        # the number of attractors found
        out_file.write(str(num) + "; ")
        # which attractors lead to coexistence and which not 
        out_file.write(np.array2string(e_flag, separator=',', suppress_small=True).replace('\n', '') + "; ")
        # trophic level means per attractor
        out_file.write(np.array2string(mean, separator=',', suppress_small=True).replace('\n', '') + "; ")
        # means of B1, I1, T1 per attractor
        out_file.write(np.array2string(mean_1, separator=',', suppress_small=True).replace('\n', '') + "; ")
        # trophic level CVs per attractor
        out_file.write(np.array2string(cv, separator=',', suppress_small=True).replace('\n', '') + "; ")
        # cvs of B1, I1, T1 per attractor
        out_file.write(np.array2string(cv_1, separator=',', suppress_small=True).replace('\n', '') + "; ")
        # cvs of B2, I2, T2 per attractor
        out_file.write(np.array2string(cv_2, separator=',', suppress_small=True).replace('\n', '') + "; ")
        # proportional size of basin of attraction for each attractor
        out_file.write(np.array2string(prop, separator=',', suppress_small=True).replace('\n', '') + "\n")
    
    # progress bar
    if i % 50 == 0:
        print('***', end = ' ')
        global ts
        print("subtime: " + str(time.time() - ts) 
                + " (" + str(i) + " / " + str(total) + ")", flush = True)
        ts = time.time()


# get relevant information on properties such as mean biomasses, CVs, also number of attractors
# per parameter combination

def calc_prop(p):
    # these values are output by C 
    Nmean = []
    Bmean = []
    Imean = []
    Tmean = []
    
    B1mean = []
    I1mean = []
    T1mean = []
    
    Ncv = []
    
    B1cv = []
    I1cv = []
    T1cv = []
    
    B2cv = []
    I2cv = []
    T2cv = []
    
    Bcv_TL = []
    Icv_TL = []
    Tcv_TL = []

    ext_flag = [] # checks if all species coexist

    
    for i in range(duplicates):
        # get new initial conditions
        inits = get_inits(p['N0'])
        # make a new dictionary to pass on to C
        new_dict = {**inits, **p}
        new_dict['id'] = int((new_dict['id'] + i) % max_output_files) # create unique ID per simulation 
                                                                      # (maximum max_output_files)
        # copy new values into c_params dictionary
        if new_dict != None:
            for key in new_dict:
                c_params[key] = new_dict[key]
        # make a string out of all arguments to be passed on to C
        args = ''
        for name in c_params:
            args += ' ' + str(c_params[name])
        # run the C code and read its output
        means, cvs, TL_cvs = run_C(args, c_params['id'])        


        # make sure we only treat cases that are physically possible (with some tolerance)
        if ((means[0] + 0.175*(sum(means[1:3]) + (sum(means[3:5]) + sum(means[5:7])/0.33)/0.33)) > c_params['N0']*2):
            print("biomass overflow")
            means = [0, 0, 0, 0, 0, 0, 0]
            cvs = [0, 0, 0, 0, 0, 0, 0]
            TL_cvs = [0, 0, 0]
        
        # c outputs the standard deviation and not CV
        for i in range(len(means)):
            if means[i] != 0:
                cvs[i] = cvs[i]/means[i]
            else:
                cvs[i] = 0
        for i in range(3):
            if (means[2*i + 1] + means[2*(i + 1)]) != 0:
                TL_cvs[i] = TL_cvs[i]/(means[2*i + 1] + means[2*(i + 1)])
            else:
                TL_cvs[i] = 0
                
        # check whether all species coexist
        ext_flag.append(getext(np.asarray(means), c_params['web_id']))
        
        # put all outcomes in Python lists to handle later
        Nmean.append(means[0])
        Bmean.append(means[1] + means[2])
        Imean.append(means[3] + means[4])
        Tmean.append(means[5] + means[6])
        
        B1mean.append(means[1])
        I1mean.append(means[3])
        T1mean.append(means[5])
        
        Ncv.append(cvs[0])
        
        B1cv.append(cvs[1])
        I1cv.append(cvs[3])
        T1cv.append(cvs[5])
        
        B2cv.append(cvs[2])
        I2cv.append(cvs[4])
        T2cv.append(cvs[6])
        
        Bcv_TL.append(TL_cvs[0])
        Icv_TL.append(TL_cvs[1])
        Tcv_TL.append(TL_cvs[2])
        
    # initialize variables to check how many unique attractors we have
    n = 0 # duplicate to check all remaining others against
    m = 1 # first "new" dupilcate to check against dupilcate n
    k = 0 # currently checking
    
    # check which duplicates are likely to lead to the same attractor
    # criterium here: all means and CVs are "close enough" together
    # this only serves to simplify the output data, we do not quantitatively 
    # analyze the number of attractors 
    counts = np.ones(duplicates)
    while n < duplicates - k:
        m = 1
        while m < duplicates - k - n:
            if (np.allclose(np.array([Nmean[n], Bmean[n], Imean[n], Tmean[n]]), 
                          np.array([Nmean[n + m], Bmean[n + m], Imean[n + m], Tmean[n + m]])
                          , atol = 5, rtol = 0.1)
                and np.allclose(np.array([Ncv[n], B1cv[n], I1cv[n], T1cv[n]]), 
                          np.array([Ncv[n + m], B1cv[n + m], I1cv[n + m], T1cv[n + m]])
                          , atol = 1e-2, rtol = 0.1)
                and np.allclose(np.array([B2cv[n], I2cv[n], T2cv[n]]), 
                          np.array([B2cv[n + m], I2cv[n + m], T2cv[n + m]])
                          , atol = 1e-2, rtol = 0.1)
                and np.allclose(np.array([Bcv_TL[n], Icv_TL[n], Tcv_TL[n]]), 
                          np.array([Bcv_TL[n + m], Icv_TL[n + m], Tcv_TL[n + m]])
                          , atol = 1e-2, rtol = 0.1)):
                    del Nmean[n + m]
                    del Bmean[n + m]
                    del Imean[n + m]
                    del Tmean[n + m]
                    
                    del B1mean[n + m]
                    del I1mean[n + m]
                    del T1mean[n + m] 
        
                    del Ncv[n + m]
                    
                    del B1cv[n + m]
                    del I1cv[n + m]
                    del T1cv[n + m]
                    
                    del B2cv[n + m]
                    del I2cv[n + m]
                    del T2cv[n + m]
                    
                    del Bcv_TL[n + m]
                    del Icv_TL[n + m]
                    del Tcv_TL[n + m]
                    
                    del ext_flag[n + m]
                    
                    k += 1
                    counts[n] += 1
            else: 
                m += 1
        n += 1        
    
    attr_num = n

    # calculate relative frequency of each attractor
    freqs = counts[0:attr_num]
    props = [] # measure of the size of the basin of attraction as a proportion of the state space
               # from which the initial conditions were sampled
    i = 0
    while i < attr_num:
        props.append(freqs[i]/sum(freqs))
        i += 1

    return np.array(ext_flag), attr_num, np.array([Nmean, Bmean, Imean, Tmean]).T,  \
            np.array([B1mean, I1mean, T1mean]).T, np.array([Ncv, Bcv_TL, Icv_TL, Tcv_TL]).T, \
            np.array([B1cv, I1cv, T1cv]).T,  np.array([B2cv, I2cv, T2cv]).T, \
            np.array(props)

# check whether all species coexist
def getext(means, id):
    # where id = web id
    if id == 0:
        indices = [0, 1, 3, 5]
        
    elif id == 1:
        indices = [0, 1, 2, 3, 5]
        
    elif id == 2:
        indices = [0, 1, 3, 4, 5]
        
    elif id == 3:
        indices = [0, 1, 2, 3, 4, 5]

    elif id == 4:
        indices = [0, 1, 3, 5, 6]
        
    elif id == 5:
        indices = [0, 1, 2, 3, 5, 6]

    elif id == 6:
        indices = [0, 1, 3, 4, 5, 6]
    elif id == 7:
        indices = [0, 1, 2, 3, 4, 5, 6]   
    
    if any(means[indices] == 0):
        ext_flag = 1
    else:
        ext_flag = 0
            
    return ext_flag

# run the C code and read its output
def run_C(args, index):
    # calling the program
    command = "../c/sys_solve" + args 
    # remove a file if it already exists 
    # (otherwise C will append to it, and then Python cannot correctly import the new values)
    if os.path.exists("../../c_output_data/mean/mean" + str(c_params['id']) 
                        + '-' + str(c_params['web_id']) + ".out"):
    	os.remove("../../c_output_data/mean/mean" + str(c_params['id']) 
                        + '-' + str(c_params['web_id']) + ".out")

    try:
        # run the C code
        c_prog = subprocess.run(command, stdout = subprocess.PIPE, stderr=subprocess.PIPE,
                            shell = True, timeout = 180)

    except subprocess.TimeoutExpired:
        # if it takes too long, assume something went wrong
        print('timeout expired')
        means = np.array([0, 0, 0, 0, 0, 0, 0])
        cvs = np.array([0, 0, 0, 0, 0, 0, 0])
        TL_cvs = np.array([0, 0, 0])
        return means, cvs, TL_cvs
    try:
        # read the output created by C
        means = np.array(pd.read_csv("../../c_output_data/mean/mean" + str(c_params['id']) 
                        + '-' + str(c_params['web_id']) + ".out", sep = '\s+',
                        header = None, usecols = [0]))
        cvs = np.array(pd.read_csv("../../c_output_data/mean/mean" + str(c_params['id']) 
                        + '-' + str(c_params['web_id']) + ".out", sep = '\s+',
                        header = None, usecols = [1]))
        TL_cvs = np.array(pd.read_csv("../../c_output_data/mean/mean" + str(c_params['id']) 
                        + '-' + str(c_params['web_id']) + ".out", sep = '\s+',
                        header = None, usecols = [2]))
    except KeyboardInterrupt:
        # make sure the program can be interrupted if necessary
        return
    except:
        # make sure the code does not stop if something went wrong (should not happen)
        print('exception')
        means = np.array([0, 0, 0, 0, 0, 0, 0])
        cvs = np.array([0, 0, 0, 0, 0, 0, 0])
        TL_cvs = np.array([0, 0, 0])
    return means, cvs, TL_cvs

# generate a new random parameter combination
def set_params(current, total):
    new_params = {}
    # make sure it is random because of the parallelization
    np.random.seed(int.from_bytes(os.urandom(4), byteorder='little'))
    
    # generate valid combination of Delta_B, Delta_I, Delta_T for the desired food web
    if webid == 1:
        current = current%4
        new_params['B_diff'] = B_diff_a[int(current)]
    if webid == 2:
        current = current%4
        new_params['I_diff'] = I_diff_a[int(current)]
    if webid == 4:
        current = current%4
        new_params['T_diff'] = T_diff_a[int(current)]
    if webid == 3:
        current = current%16
        new_params['B_diff'] = B_diff_a[int(current//4)]
        new_params['I_diff'] = I_diff_a[int(current % 4)]
    if webid == 5:
        current = current%16
        new_params['B_diff'] = B_diff_a[int(current//4)]
        new_params['T_diff'] = T_diff_a[int(current % 4)]
    if webid == 6:
        current = current%16
        new_params['I_diff'] = I_diff_a[int(current//4)]
        new_params['T_diff'] = T_diff_a[int(current % 4)]
    if webid == 7:
        current = current%64
        new_params['B_diff'] = B_diff_a[int(current//16)]
        new_params['I_diff'] = I_diff_a[int((current % 16)//4)]
        new_params['T_diff'] = T_diff_a[int(current % 4)]
        
    
    
    # all other parameters are randomly drawn from uniform distributions
    indices = np.random.rand(11)

    indices[0] = indices[0]*(N0_a[1] - N0_a[0]) +  N0_a[0]
    indices[1] = indices[1]*(h_N_a[1] - h_N_a[0]) +  h_N_a[0]
    indices[2] = indices[2]*(r_a[1] - r_a[0]) +  r_a[0]
    indices[3] = indices[3]*(a_a[1] - a_a[0]) +  a_a[0]
    indices[4] = indices[4]*(h_a[1] - h_a[0]) +  h_a[0]
    indices[5] = indices[5]*(alpha_a[1] - alpha_a[0]) +  alpha_a[0]
    indices[6] = indices[6]*(eta_a[1] - eta_a[0]) +  eta_a[0]
    indices[7] = indices[7]*(delta_a[1] - delta_a[0]) +  delta_a[0]
    indices[8] = indices[8]*(n_a[1] - n_a[0]) +  n_a[0]
    indices[9] = indices[9]*(nu_a[1] - nu_a[0]) +  nu_a[0]
    indices[10] = indices[10]*(a_scale_a[1] - a_scale_a[0]) + a_scale_a[0]

    new_params['N0'] = indices[0]
    new_params['h_N'] = indices[1]
    new_params['r0'] = indices[2]
    new_params['a0'] = indices[3]
    new_params['h0'] = indices[4]
    new_params['alpha0'] = indices[5]
    new_params['eta0'] = indices[6]
    new_params['delta'] = indices[7]
    new_params['n'] = indices[8]
    new_params['nu'] = indices[9]
    new_params['a_scale'] = indices[10]

    return new_params

# generate random initial conditions
def get_inits(N0):
    init_bounds = [2*N0, 2*N0/c_params['ratio'], 2*c_params['e_i']*N0/c_params['ratio'], 
                   2*c_params['e_i']*c_params['e_t']*N0/c_params['ratio']]
    inits = {}
    # nutrients
    inits['N_0'] = init_bounds[0]*rd.random()
    
    # basal species
    B_max = init_bounds[1]*rd.random()
    B1 = B_max*rd.random()
    inits['B1_0'] = B1
    inits['B2_0'] = B_max - B1
    
    # intermediate species
    I_max = init_bounds[2]*rd.random()
    I1 = I_max*rd.random()
    inits['I1_0'] = I1
    inits['I2_0'] = I_max - I1
    
    # top species
    T_max = init_bounds[3]*rd.random()
    T1 = T_max*rd.random()
    inits['T1_0'] = T1
    inits['T2_0'] = T_max - T1
    
    if c_params['web_id'] == 0:
        inits['B1_0'] += inits['B2_0']
        inits['I1_0'] += inits['I2_0']
        inits['T1_0'] += inits['T2_0']
        
        inits['B2_0'] = 0
        inits['I2_0'] = 0
        inits['T2_0'] = 0
        
    elif c_params['web_id'] == 1:
        inits['I1_0'] += inits['I2_0']
        inits['T1_0'] += inits['T2_0']
        
        inits['I2_0'] = 0
        inits['T2_0'] = 0
        
    elif c_params['web_id'] == 2:
        inits['B1_0'] += inits['B2_0']
        inits['T1_0'] += inits['T2_0']
        
        inits['B2_0'] = 0
        inits['T2_0'] = 0
        
    elif c_params['web_id'] == 4:
        inits['B1_0'] += inits['B2_0']
        inits['I1_0'] += inits['I2_0']
        
        inits['B2_0'] = 0
        inits['I2_0'] = 0

    elif c_params['web_id'] == 3:
        inits['T1_0'] += inits['T2_0']
        
        inits['T2_0'] = 0
        
    elif c_params['web_id'] == 5:
        inits['I1_0'] += inits['I2_0']
        
        inits['I2_0'] = 0

    elif c_params['web_id'] == 6:
        inits['B1_0'] += inits['B2_0']
        
        inits['B2_0'] = 0

    return inits

# =============================================================================
# "main"
# =============================================================================


t0 = time.time()
ts = time.time()


run_parallel()
te = time.time()
print('code execution time: ', te-t0, 'sec')

 
```
