## Supplementary material for "Top predators govern multitrophic diversity effects in tritrophic food webs": Code and data: figS3-3.py

```
#!/usr/bin/env python3
# -*- coding: utf-8 -*-
"""
Created on Fri Jun 5 2020

@author: ruben
"""
# this script will generate Figure S3.3 (Appendix S3)

import numpy as np
import pandas as pd   # for fast reading of files
import collections    # for the ordered dict
import subprocess     # for the execution of the C program
import os
import matplotlib.pyplot as plt
from scipy.integrate import odeint

# %%===========================================================================
# specify parameters and initial conditions to plot
# =============================================================================

# "initialize" dictionary
c_params = collections.OrderedDict()

c_params['tend'] = 1e4 # relax to attractor !!! for this long time a very large file (~680MB) will be created
c_params['N0'] = 666.2
c_params['h_N'] = 18.92
c_params['r0'] = 1.874
c_params['r_inc'] = 0.5
c_params['h0'] = 0.7924
c_params['h_dec'] = 0.5
c_params['eta0'] = 3.209
c_params['eta_dec'] = 0.5
c_params['a0'] = 1.518e-3
c_params['alpha0'] = 3.299e-4
c_params['a_scale'] = 1.290

c_params['delta'] = 0.05710
c_params['e_i'] = 0.33
c_params['e_t'] = 0.33
c_params['n'] = 1.907

c_params['nu'] = 1.363
c_params['ratio'] = 0.175

c_params['teval'] = 3e3 # eval mean
c_params['TIMESERIES'] = 1 # put to 1 if timeseries should be written to file
c_params['POP_STATS'] = 1
c_params['id'] = 0  # change to another int if previous files should not be overwritten
c_params['web_id'] = 0 # does not matter in this case

c_params['N_0'] = 10
c_params['B1_0'] = 1000
c_params['B2_0'] = 1000
c_params['I1_0'] = 100
c_params['I2_0'] = 100
c_params['T1_0'] = 100
c_params['T2_0'] = 100

c_params['B_diff'] = 0.75
c_params['I_diff'] = 0.75
c_params['T_diff'] = 0.25

# %%===========================================================================
# create and import output files
# =============================================================================

args = ''
for name in c_params:
    args += ' ' + str(c_params[name])
        
command = "../c/sys_solve" + args 
# remove a file if it already exists 
# (otherwise C will append to it, and then Python cannot correctly import the new values)
if os.path.exists("../../c_output_data/mean/mean" + str(c_params['id']) 
                    + '-' + str(c_params['web_id']) + ".out"):
	os.remove("../../c_output_data/mean/mean" + str(c_params['id']) 
                    + '-' + str(c_params['web_id']) + ".out")
# run the C code
c_prog = subprocess.run(command, stdout = subprocess.PIPE, stderr=subprocess.PIPE,
                            shell = True)

# read the output created by C
t = np.array(pd.read_csv("../../c_output_data/timeseries/timeseries" + str(c_params['id']) 
                + '-' + str(c_params['web_id']) + ".out", sep = '\s+',
                header = None, usecols = [0]))
B = np.array(pd.read_csv("../../c_output_data/timeseries/timeseries" + str(c_params['id']) 
                + '-' + str(c_params['web_id']) + ".out", sep = '\s+',
                header = None, usecols = np.arange(7) + 1))
means = np.array(pd.read_csv("../../c_output_data/mean/mean" + str(c_params['id']) 
                + '-' + str(c_params['web_id']) + ".out", sep = '\s+',
                header = None, usecols = [0]))
cvs = np.array(pd.read_csv("../../c_output_data/mean/mean" + str(c_params['id']) 
                + '-' + str(c_params['web_id']) + ".out", sep = '\s+',
                header = None, usecols = [1]))
TL_cvs = np.array(pd.read_csv("../../c_output_data/mean/mean" + str(c_params['id']) 
                + '-' + str(c_params['web_id']) + ".out", sep = '\s+',
                header = None, usecols = [2]))

# %%===========================================================================
# plot timeseries
# =============================================================================
labels = [r'$N$', r'$B_1$',r'$B_2$', r'$I_1$', r'$I_2$', r'$T_1$', r'$T_2$']
colors = ['lightskyblue', 'seagreen', 'seagreen', 'chocolate', 'chocolate'
          , '#8D1B2C', '#8D1B2C']
linestyles = ['-', '-', '--', '-', '--', '-', '--'] 

fig = plt.figure(figsize = (8, 5))
for i in range(7):
    plt.plot(t[-15000:], B[-15000:, i], linewidth = 2, linestyle = linestyles[i],
             color = colors[i], label = labels[i])
plt.yscale('log')
plt.legend(fontsize = 14)
plt.tick_params(axis='both', which='major', labelsize=14)
plt.xlabel(r't', fontsize = 16)
plt.ylabel(r'Biomass', fontsize = 16)
plt.tight_layout()
plt.savefig('S33.pdf')
```
